# Nascent transcript O-MAP reveals the molecular architecture of an RBM20-regulated single-locus subnuclear compartment organized around the *TTN* RNA

**DOI:** 10.1101/2024.11.05.622011

**Authors:** Evan E. Kania, Sara Bianchi, Aidan Fenix, Daphnée M. Marciniak, Shuai Li, Qiaoyi Lin, Marilù Casini, Borislav Hristov, Lukasz Truszkowski, Nicholas G. Skiados, Milad Mokhtaridoost, Conor K. Camplisson, Rose Fields, Elisabeth Wyart, Silvia Marchiano, Brian J. Beliveau, Philipp G. Maass, Davide Cacchiarelli, Elisa Balmas, Devin K. Schweppe, William S. Noble, Shao-En Ong, Charles E. Murry, Alessandro Bertero, David M. Shechner

## Abstract

Eukaryotic gene expression is regulated by highly organized subnuclear compartmentalization, yet the molecular architecture of locus-specific nuclear microenvironments remains difficult to dissect. Here, we adapt oligonucleotide-mediated proximity-interactome mapping (O-MAP) to map genomic loci, RNAs, and proteins within discrete nuclear neighborhoods, using a cardiac-specific RNA factory organized around nascent *TTN* RNA as a model. We find that the *TTN*-associated microenvironment is embedded within a developmentally regulated hub of chromosomal interactions, including *cis-*interacting active domains and *trans*-interacting poised and repressed domains. In addition to chromatin contacts, the factory contains diverse RNAs in a pattern consistent with substantial post-transcriptional recruitment and is enriched for RNA-metabolic and chromatin-regulatory proteins. These include the splicing factor RBM20 and chromatin-scaffolding factor SAFB, which exert opposing effects on factory-associated gene expression. Disruption of factory architecture through loss of *TTN* RNA or RBM20 leads to widespread splicing defects and mislocalization of transcripts. Notably, many splicing defects caused by RBM20 loss are attenuated by *TTN* RNA depletion, indicating that *TTN’s* unbound transcript exerts toxic gain-of-function effects on RNA metabolism in the absence of RBM20. Collectively, this work establishes O-MAP as a powerful approach for biochemically dissecting single-locus compartments and reveals how coding RNAs can influence nuclear architecture and gene regulation.

## MAIN

Mammalian nuclei adopt a highly compartmentalized architecture that enables spatial control over most functions performed on, or by, their genomes. Core nuclear functions like transcription, RNA processing, and DNA repair^1-3^ are often coordinated in dedicated, micrometer-scale compartments that concentrate regulatory proteins and RNAs^4-7^ and spatially cluster co-regulated loci^8,9^. A single nucleus may contain dozens to thousands of instances of a given compartment type (*e.g.*, nucleoli, nuclear speckles, Cajal bodies), each built on a characteristic molecular architecture conserved within that compartment type ^9-11^. However, individual instances are thought to modify this core architecture to support the specific needs of the loci they regulate, raising the possibility that similar compartments may perform distinct, locus-specific functions. Yet, exploring this hypothesis has proven challenging, given the lack of robust technologies for biochemically characterizing the distinct microenvironments surrounding individual target loci. As a result, the functional diversity of single-locus nuclear compartments, and the mechanisms by which this diversity is programmed, remain largely unexplored.

We recently identified one such specialized compartment—an RNA factory that plays a key regulatory role during cardiac development and homeostasis^12^. In cardiomyocytes (CMs), this factory assembles around nascent primary transcripts encoding the giant protein Titin (*TTN*) and nucleates prominent foci of the splicing factor RBM20, a key regulator of *TTN* mRNA biogenesis^13,14^. Genomic loci encoding other RBM20 targets (located on several chromosomes) also localize near or within these RBM20 foci, which appears to influence their alternative splicing. These observations suggest that *TTN* RNA factories might act as "crucibles" that coordinate the biogenesis of cardiomyogenic genes^12^. Previous studies indicate that RBM20 foci are a class of nuclear speckles^15^—RNA-rich nuclear bodies that coordinate pre-mRNA splicing^5,11^. However, while cardiomyocytes contain dozens of nuclear speckles, only those located near the *TTN* locus compartmentalize RBM20 and its target genes, suggesting their functional specialization^12^. Similar models of RNA-mediated compartmentalization have been proposed for other subnuclear structures—including nucleoli, Barr bodies, paraspeckles, and nuclear stress bodies (nSBs)^16-18^. Yet, while these compartments are assembled on long noncoding RNAs, such architectural roles have rarely been attributed to pre-mRNAs^19,20^ Thus, the *TTN* RNA factory may represent an underexplored mechanism of nuclear compartmentalization in which coding loci "moonlight" as sources of chromatin-regulatory RNAs^21^.

Proper assembly of the *TTN* RNA factory may also be critical for human health. Mutations in RBM20, including those that sequester it outside of the nucleus^22^, cause a malignant form of dilated cardiomyopathy (DCM) characterized by conduction system defects, ventricular arrhythmia, and rapid progression to heart failure^23,24^. Current therapies can only manage these symptoms; the sole effective treatment is heart transplantation^23,24^. Much of the cardiac dysfunction in RBM20–DCM can be ascribed to defects in *TTN* splicing, which produce an elastic, fetal-like protein isoform that severely weakens left ventricular function^13^. However, other facets of disease etiology, including calcium current aberrations, cannot readily be explained by the missplicing of RBM20 target transcripts^25^. We therefore hypothesized that nuclear depletion of RBM20 also disrupts *TTN* RNA factory architecture, with downstream consequences for chromatin organization and gene expression.

To explore this hypothesis, we sought an experimental strategy that enables targeted biochemical characterization of discrete subnuclear compartments like the *TTN* RNA factory—an approach for which few robust tools are currently available. Most current approaches for characterizing nuclear architecture have focused on the looping and folding of chromatin at the nanometer-scale^26,27^ and are not well suited for probing the higher-order, micron-scale compartmental interactions that are pervasive in nuclear organization^28^, including *trans-*chromosomal contacts^29-31^. Newer genomics approaches that probe such structures are technically demanding due to complex split-pool indexing (SPRITE^28^) or laser capture and cryosectioning manipulations (GAM^32^) requiring dozens of steps, and have limited dynamic range^33^. And, because these techniques profile all compartments *en masse,* they might lack the sensitivity to detect interactions specific to individual loci. In principle, such compartment-specific interactions could be captured using pulldown-based strategies—for example, by immunoprecipitation of RBM20, or using antisense oligonucleotide-based capture of the *TTN* pre-mRNA or DNA^34-36^. However, these methods would be difficult to apply to compartments like the *TTN* RNA factory, which is low in abundance and may be too fragile to withstand biochemical purification.

We reasoned that these barriers could be overcome using *in situ* proximity biotinylation. In this approach, promiscuous biotinylating enzymes (*e.g.* Horseradish Peroxidase (HRP) or APEX), are used to covalently tag biomolecules within a defined subcellular compartment, enabling unbiased identification of local molecular interactions without complex fractionation or pulldown^37^. Proximity labeling has been previously adapted to study nuclear structure—for example, by using CRISPR/Cas9 to localize APEX to target genomic loci^38,39^, or antibody-HRP conjugates to label all compartments of a given type^40^. However, these approaches are often constrained by large pools of mislocalized or off-targeted enzymes^41^ and thus lack the targeting specificity required to resolve interactions within individual, single-locus compartments.

To overcome this challenge, we adapted **O**ligonucleotide-mediated proximity-interactome **MAP**ping (**O–MAP**)^41^—an RNA-targeted proximity-biotinylation approach—to biochemically characterize discrete subnuclear compartments with high spatial specificity. By directing O-MAP to nascent introns of the *TTN* primary transcript, rather than the mature mRNA, we profile the chromatin loci, transcripts, and proteins that co-compartmentalize with nascent *TTN* and examine how this architecture is remodeled upon loss of RBM20. This strategy enables us to move beyond a descriptive view of RNA factories to address their molecular organization and regulation. In doing so, we define how RBM20 and *TTN* RNA contribute to compartment organization and broader RNA regulatory programs during cardiac development, with potential relevance to RBM20-associated dilated cardiomyopathy. To our knowledge, this work provides the first biochemical characterization of a single-locus subnuclear compartment, and it establishes a framework for dissecting how nascent transcripts can contribute to nuclear architecture in development and disease.

## RESULTS

### Nascent transcript O-MAP enables precise in situ proximity-biotinylation of the TTN RNA Factory

O-MAP uses programmable DNA oligonucleotide probes (similar to those used in RNA-FISH)^42,43^ to recruit proximity-labeling enzymes to an RNA of interest within its native context^41^ (**Fig. 1a**). In O-MAP, samples are chemically fixed, and oligo probes are hybridized to the target RNA *in situ*. These probes are appended with a universal "landing pad" sequence that, in a subsequent hybridization step, recruits a secondary oligo that is chemically conjugated to Horseradish Peroxidase (HRP). Upon addition of biotin-tyramide and hydrogen peroxide, HRP generates short-lived biotinyl radicals that diffuse outward (∼50–75 nm)^44^ and covalently tag nearby molecules, enabling their capture and analysis.

**Figure 1.**
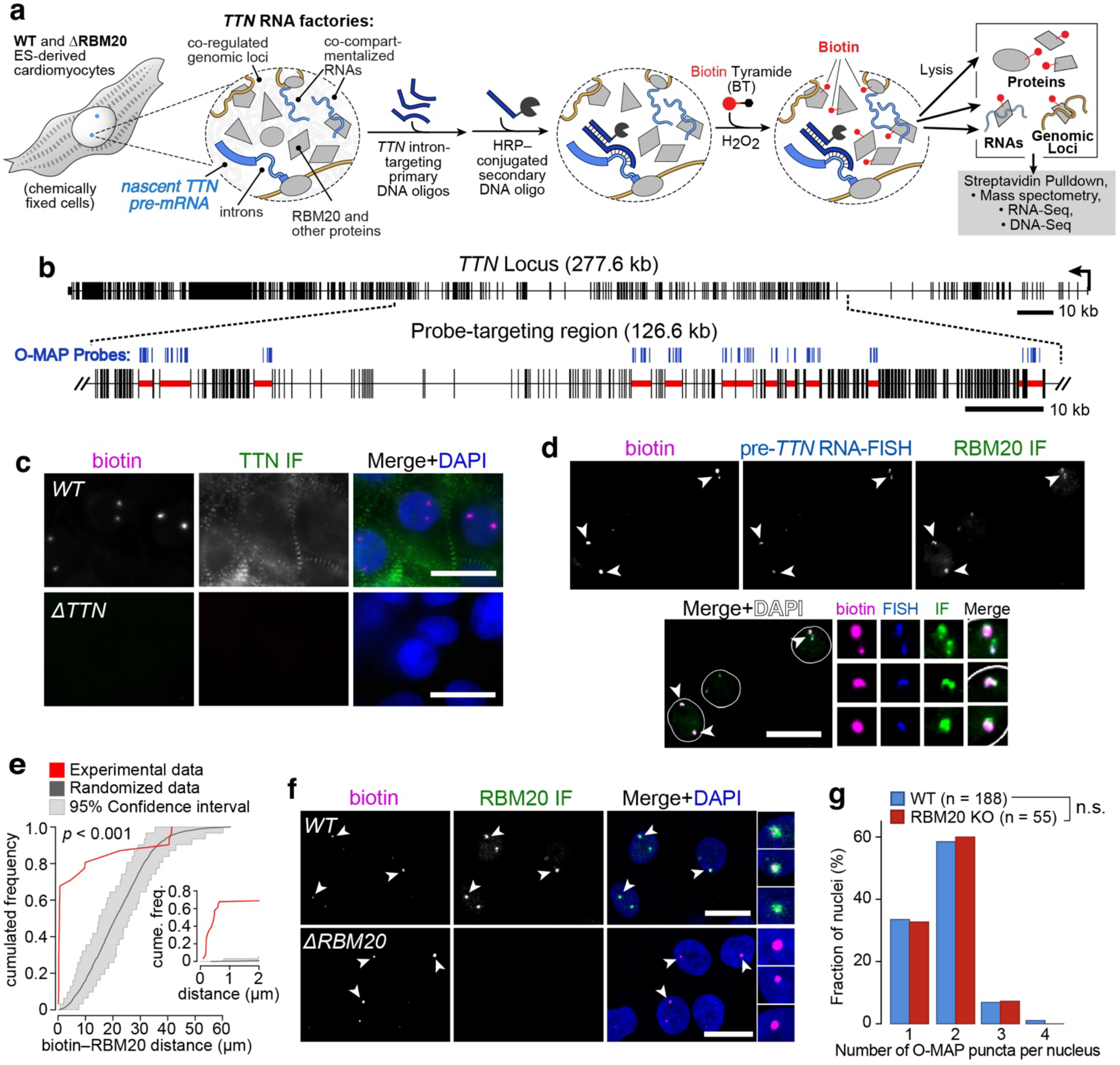
Microenvironment-mapping of the *TTN* RNA factory by nascent transcript O-MAP. **a,** Approach. WT and RBM20 KO hESC-derived cardiomyocytes (CMs)^12^ were chemically fixed, and pools of antisense DNA probes were hybridized to the introns on the *TTN* pre-mRNA. These probes recruit a common HRP-conjugated oligo that catalyzes *in situ* proximity-biotinylation. Biotinylated material was enriched by streptavidin pulldown, and interacting factors (proteins, RNAs, genomic loci) were isolated for downstream analyses^41^. **b,** O-MAP probes (*blue*) targeted *TTN* introns previously identified as RBM20 targets (*red highlight*)^15^. **c**–**f,** O-MAP enables highly precise *in situ* biotinylation in *TTN* RNA factories. **c,** In WT CMs (*top*) O–MAP induced two prominent biotinylation puncta per diploid nucleus; this signal is ablated in cells lacking the *TTN* promoter (ΔTTN, *bottom*)^12^. IF: immunofluorescence. **d,** Combined O-MAP, RNA-FISH, and RBM20 immunofluorescence demonstrates that O-MAP specifically labels *TTN* RNA factories. Arrowheads denote colocalization, highlighted in zoomed images (*bottom right*). **e**, Object-based quantification of the pre-*TTN* O-MAP—RBM20 colocalization distances, as observed in **d**. The distribution of the distances from experimental images (*red*) falls outside the confidence interval of the distribution obtained for 100 images in which object locations are randomly shuffled (*gray*). Inset highlights data below 2 µm. **f,** Representative images show that pre-*TTN* O-MAP puncta remain readily detectable in RBM20 KO cells. **g,** O-MAP puncta were unchanged by loss of RBM20. n.s.: not significant, two-sided Mann–Whitney U test. All scale bars: 20 µm; zoomed images are ∼3x.

We sought to apply O-MAP to the *TTN* RNA factory by targeting the RNA "scaffold" around which this factory nucleates: the primary *TTN* RNA^12^. To pinpoint these unspliced nuclear transcripts (and avoid mature cytoplasmic mRNA), we designed O-MAP probes against intronic sequences within the *TTN* gene, focusing on introns known to be bound and regulated by RBM20^15^ (**Fig. 1b**; **Supplementary Table 1**). Because TTN spans more than 280 kb, probes placed elsewhere along the transcript could sample a partially distinct local neighborhood. We applied these probes in an established cell culture model that generates highly pure (>90% cardiac troponin T-positive) cardiomyocytes by *in vitro* differentiation from RUES2 human embryonic stem cells (hESC-CMs)^12^. Parallel experiments in an isogenic, RBM20 knockout line^12^ (RBM20 KO hESC-CMs) enabled us to also map disruptions in pre-*TTN* factory architecture upon RBM20 depletion. The RBM20 KO cell line was generated via CRISPR/Cas9, using sgRNAs designed towards RBM20 exon 2, as described previously^12^. Although such homozygous RBM20 deletions are not observed in human DCM patients^45-47^, this line provides a biochemically clean model to study the effect of RBM20 nuclear depletion (which is observed with most disease-associated variants), while avoiding potentially confounding effects from mutant RBM20 aggregation in the cytoplasm^22,48-50^.

We began by validating the targeting precision of our O-MAP probe set. In wild-type (WT) hESC-CMs, pre-*TTN* O-MAP induced two prominent (∼1 µm diameter) biotinylation puncta within each diploid nucleus— corresponding to the two alleles of *TTN—*with undetectable background signal elsewhere (**Fig. 1c**). O-MAP puncta were not observed in hESC-CMs in which both *TTN* alleles had been CRISPR-edited to remove the promoter of full-length Titin^12^ (Δ*TTN* Pro), demonstrating the high specificity of our approach (**Fig. 1c**, *bottom*). In WT cells, biotinylation puncta exclusively overlapped with nascent *TTN* transcripts, as gauged by intron-targeting RNA-FISH^41^, indicating that O-MAP had targeted pre-*TTN* with high precision (**Fig. 1d**). These puncta also colocalized with foci of the RBM20 protein, as gauged by immunofluorescence, with an average O-MAP–RBM20 displacement near the resolution limit of conventional microscopy (overlap *p-*value < 0.01; *n*=100 cells; **Fig. 1d, e**). Importantly, in RBM20 KO hESC-CMs pre-*TTN* O-MAP puncta remained readily detectable and appeared qualitatively comparable to those of WT cells (**Fig. 1f–g**). This suggests that any differences observed between WT and RBM20 KO cells are likely not artifacts of differential O-MAP labeling of the *TTN* microenvironment, but rather reflect *bona fide* remodeling of this microenvironment and/or of RBM20-dependent *TTN* biogenesis upon RBM20 loss.

### The TTN RNA factory forms a hub of cis- and trans-chromosomal interactions

The *TTN* RNA factory is thought to organize a network of higher-order chromatin interactions that recruits and compartmentalizes loci from multiple chromosomes, potentially facilitating their co-regulation^12^. We sought to elucidate this network using O-MAP-ChIP, which captures chromatin loci near a target RNA (here, the unspliced *TTN* pre-mRNA) by pulling down the biotinylated proteins to which these loci are bound^41^ (**Fig. 2a**). Data are compared with control O-MAP-ChIP samples in which the primary *TTN-*targeted probes were omitted ("Mock").

**Figure 2.**
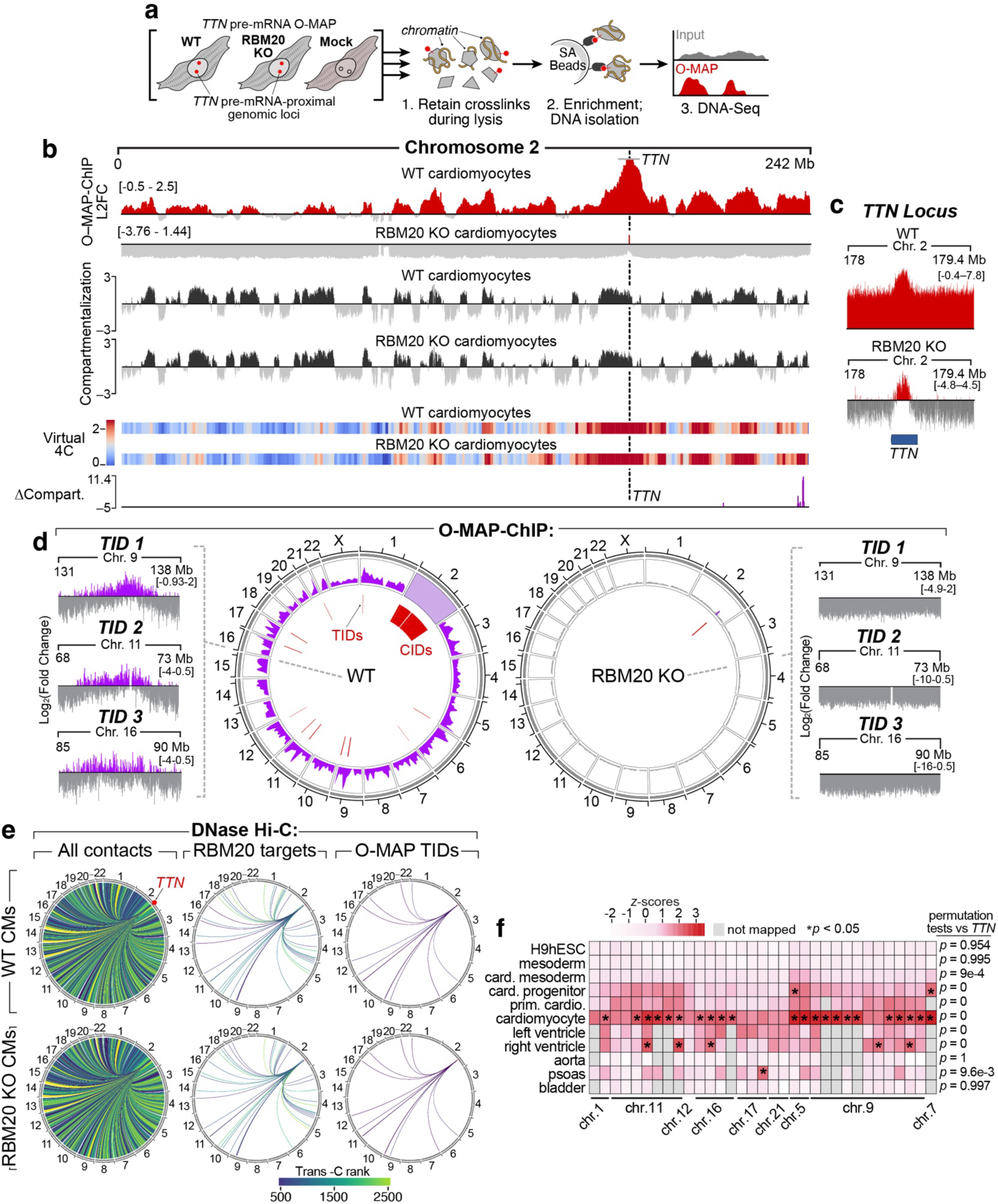
The *TTN* RNA factory forms a developmentally regulated hub of *cis* and *trans* chromosome interactions. **a,** Approach. pre-*TTN* O-MAP-ChIP^41^ was applied to WT and RBM20 KO CMs, and in negative control ("mock") samples omitting primary probes. **b,** In WT cells, O-MAP-ChIP revealed Mb-scale *cis* interacting domains (CIDs) along chromosome 2 (*top*), broadly paralleling that chromosome’s A/B compartmentalization and long-range pairwise interactions with the *TTN* locus (*middle*). Compartmentalization and interlocus interactions were largely unchanged in RBM20 KO cells (*bottom*). O-MAP data are L2FC: log_2_(fold change, enriched/input). A/B compartmental data are the quartile-normalized, p-value-adjusted eigenvectors derived from the DNase Hi-C correlation matrix^12^. Virtual 4C data are ICE- and inter-sample normalized interactions anchored on the *TTN* promoter. Changes in compartmentalization between samples (Δcompart.) are the -log10(p-value) adjusted ratio (RBM20 KO/WT). **c**, In RBM20 KO cells, nearly all pre-*TTN* proximal chromatin enrichment detected by O-MAP-ChIP is lost, with only a single statistically significant peak (FDR 0.05, EDD peak score > 40), observed with the *TTN* locus itself (*black bar*). **d,** Genome-wide views of all O-MAP-ChIP data, and highlighting representative novel TIDs, in WT (*left*) and RBM20 KO CMs (*right*). Data are L2FC (enriched/input). **e,** Significant interactions between *TTN* and O-MAP TIDs are observable in Hi-C data. Trans-C analysis of WT (*top*) and RBM20 KO (*bottom*) DNase-HiC data, using the *TTN* locus as seed. Note the relative strength of O-MAP TIDs and previously established interactions with RBM20 target loci. Relative interaction rankings were largely preserved upon RBM20 loss. **f,** *Signature* analysis of publicly available Hi-C datasets of various cell types reveals the developmental trajectory of *TTN*-TID interactions. *p*-values (starred) derived by permutation testing vs. all *trans*-contacts genome-wide, and all contacts associated with the *TTN-*locus.

In WT hESC-CMs, O-MAP-ChIP revealed the prominent enrichment of megabase-scale domains spanning the entirety of chromosome 2 (on which the *TTN* gene is itself located), which we term *Cis*-Interacting Domains (CIDs; **Figs. 2b** and **c**, *top*). Notably, CID periodicity approximates the A/B compartmentalization^2^ of chromosome 2 mapped by DNase Hi-C. This spatial segregation distinguishes the A compartment—which is generally associated with open, transcriptionally active chromatin—from the B compartment, which is generally enriched for heterochromatic, transcriptionally inactive chromatin, with individual CIDs corresponding to the open A compartment. (average Pearson’s correlations: O-MAP vs Hi-C *r* = 0.463; Mock vs Hi-C *r* = –0.113, **Fig. 2b**, *middle*, and **Supplementary Figure 1**). This suggests that, at the chromatin level, nascent *TTN* transcripts predominantly engage with other active genes on chromosome 2, consistent with the prevailing "chromosome territory" model of nuclear organization^51^. However, we also observed the marked enrichment of 19 megabase-scale (average ∼5.15 Mb) domains located on nine other chromosomes, which we term *trans*-Interacting Domains (TIDs; **Fig. 2d**, *left* and **Supplementary Table 2**). Such *trans*-contacts are challenging to study by conventional proximity-ligation-based analysis tools (*e.g.*, Hi-C)^2,29^, which highlights O-MAP’s unique ability to probe compartment-level chromatin architecture.

Remarkably, in the absence of RBM20, the *TTN* pre-mRNA appears to lose all forms of higher-order chromatin associations. In RBM20 KO hESC-CMs, pre-*TTN* O-MAP-ChIP revealed a complete absence of CIDs and TIDs, though a single notable peak centered on the *TTN* locus confirmed that its transcripts were still affiliated with the site of their transcription (**Figs. 2c**, *bottom,* and **2d**, *right*). This apparent loss of long-range and *trans* contacts suggests either that RBM20 is required to assemble the factory’s chromatin architecture, or that, in its absence, nascent *TTN* transcripts cannot maintain their association with this architecture. To distinguish between these possibilities, we used DNase Hi-C to map genome-wide chromatin folding patterns in RBM20 KO hESC-CMs. This revealed that higher-order chromatin architecture of chromosome 2 is largely unaltered by RBM20 loss: we observed no significant changes in its global A/B compartmentalization (average Pearson’s correlation between WT and RBM20 KO eigenvectors = 0.957), nor in long-range contacts centered on the *TTN* locus (**Figs. 2b**, *bottom* and **Supplementary Fig. 1**; average Stratum-adjusted correlation coefficient^52^, WT *vs* RBM20 KO = 0.985). These findings suggest that the observed loss of *TTN*-chromatin contacts reflects a defect in transcript association with chromatin architecture, rather than widespread disruption of higher-order folding of chromosome 2.

We next examined the distance of O-MAP TIDs and the *TTN* locus using multicolor DNA SABER-FISH^53^ (**Supplementary Fig. 2** and **Supplementary Table 3**). In WT cells, nearly all tested loci (seven of eight TIDs probed) compartmentalized with *TTN*, with median locus–*TTN* distances of 0.65 µm (CID) and 1.2–1.8 µm (TIDs), comparable to the spatial scale of nuclear speckles and nuclear stress bodies^7^. Although these average distances lie beyond the estimated O-MAP labeling radius^44,54,55^, the dynamic organization of the *TTN* RNA factory likely generates a sub-population of cells in which close-range encounters are sufficient to be captured by O-MAP (**Supplementary Discussion**). Notably, in RBM20 KO cells, these higher-order interactions were substantially relaxed, with median locus–*TTN* distances increasing to 1.3 µm (CID) and 1.9–2.7 µm (TIDs)— roughly equivalent to that of a negative control locus on chromosome 4. These imaging analyses provide orthogonal support for O-MAP-defined TID proximity and reveal that RBM20 loss relaxes *TTN*-centered higher-order spatial organization.

We next examined the spatial organization of newly identified TIDs in DNase Hi-C data. For this, we applied the trans-C pipeline, which uses a random-walk with restart algorithm to identify cliques of *trans*-chromosomal contacts centered on specified seed loci^29^. Application of this pipeline in WT hESC-CMs revealed prominent associations between the *TTN* locus and all O-MAP TIDs, with interaction strengths exceeding those observed with RBM20 target loci^12,29^ (**Fig. 2e**, *top;* receiver operating characteristic areas under the curve (AUCs): O-MAP TIDs: 0.928; RBM20-targets: 0.662; **Supplementary Fig. 3a**). In RBM20 KO hESC-CMs these *trans*-contacts were largely preserved, indicating that *TTN*-TIDs remain among the predominant *trans*-chromosomal interactions of the *TTN* locus despite the increased spatial separation observed by DNA-FISH.

The prominence of *TTN*-TID associations in cardiomyocytes prompted us to ask whether this architecture is developmentally regulated or present in other tissue contexts. For this, we applied the *Signature* pipeline^31^, which identifies *cis*- and *trans-*chromosomal interactions genome-wide, to a cohort of Hi-C datasets that span cardiomyogenesis and differentiated muscle-related tissues (*i.e.*, aorta, bladder, psoas, ventricles; **Supplementary Fig. 3b**–**d**)^57-59^. This analysis revealed a pronounced developmental trajectory for the majority of *TTN*–TID and *inter*-TID interactions (22/33; 66.67%), which progressively strengthened along the cardiac lineage and peaked in mature cardiomyocytes, suggesting that *TTN*-TIDs form a unique subnuclear clique (**Fig. 2f**). Some TIDs (*i.e.*, domains on chromosomes 9 and 11) overlap with previously reported "anchor loci," which exhibit strong *trans*-chromosomal proximity across diverse cell types and are frequently localized to nuclear speckles^31^. Notably, although many *inter*-TID contacts remained invariant following RBM20 loss, interactions between loci on chromosomes 9, 17, and 21 were significantly weakened (Mann-Whitney test, *p* < 0.032; **Supplementary Fig. 3f**). Together with our trans-C analysis, these data indicate that the core *TTN*-centered contact network can persist without RBM20, whereas RBM20 contributes to its compaction and to the close association of nascent *TTN* RNA with that network.

### TTN RNA factory integrity fine-tunes the chromatin landscape and transcriptional programs of TID genes

We next sought to identify epigenomic signatures that might control chromatin localization to the *TTN* RNA factory and examine how this localization influences gene expression. ChromHMM analysis^60^ using established epigenomic datasets for RUES2 hESC-CMs^61,62^ revealed that, in aggregate, *TTN-*interacting loci were enriched in genic enhancer domains (EnhG1 and EnhG2; **Fig. 3a**, *top*). However, this enrichment was largely driven by CIDs, consistent with their localization in the "A compartment" which is broadly thought to support transcription^63^ (**Figs. 2b, 3a**, *middle*). Indeed, pre-*TTN* O-MAP puncta frequently overlapped with prominent foci of H3K4me3— an activating histone modification—irrespective of RBM20 expression (**Supplementary Fig. 4a**–**b**). In contrast, TID loci were predominantly enriched in bivalent (TssBiv; EnhBiv) and repressive chromatin states (ReprPC; ReprPCWk; **Fig. 3a**, *bottom*), challenging our initial model that *TTN* factories function exclusively as crucibles of RBM20-mediated RNA biogenesis^12^. Notably, TID-enclosed genes are enriched in annotations for cardiac disorders (**Fig. 3b**) and include developmentally regulated transcription factors with established roles in stem cell differentiation and cardiomyogenesis. These include regulators of ventricular development and cardiac conduction system maturation, processes frequently perturbed in RBM20-DCM^45,64^ (Metascape metric *q* < 0.05; **Supplementary Fig. 4c**). We thus hypothesized that the fine-tuning of TID epigenetic and transcriptional programs, mediated in part by RBM20, may contribute to proper cardiomyocyte development.

**Figure 3.**
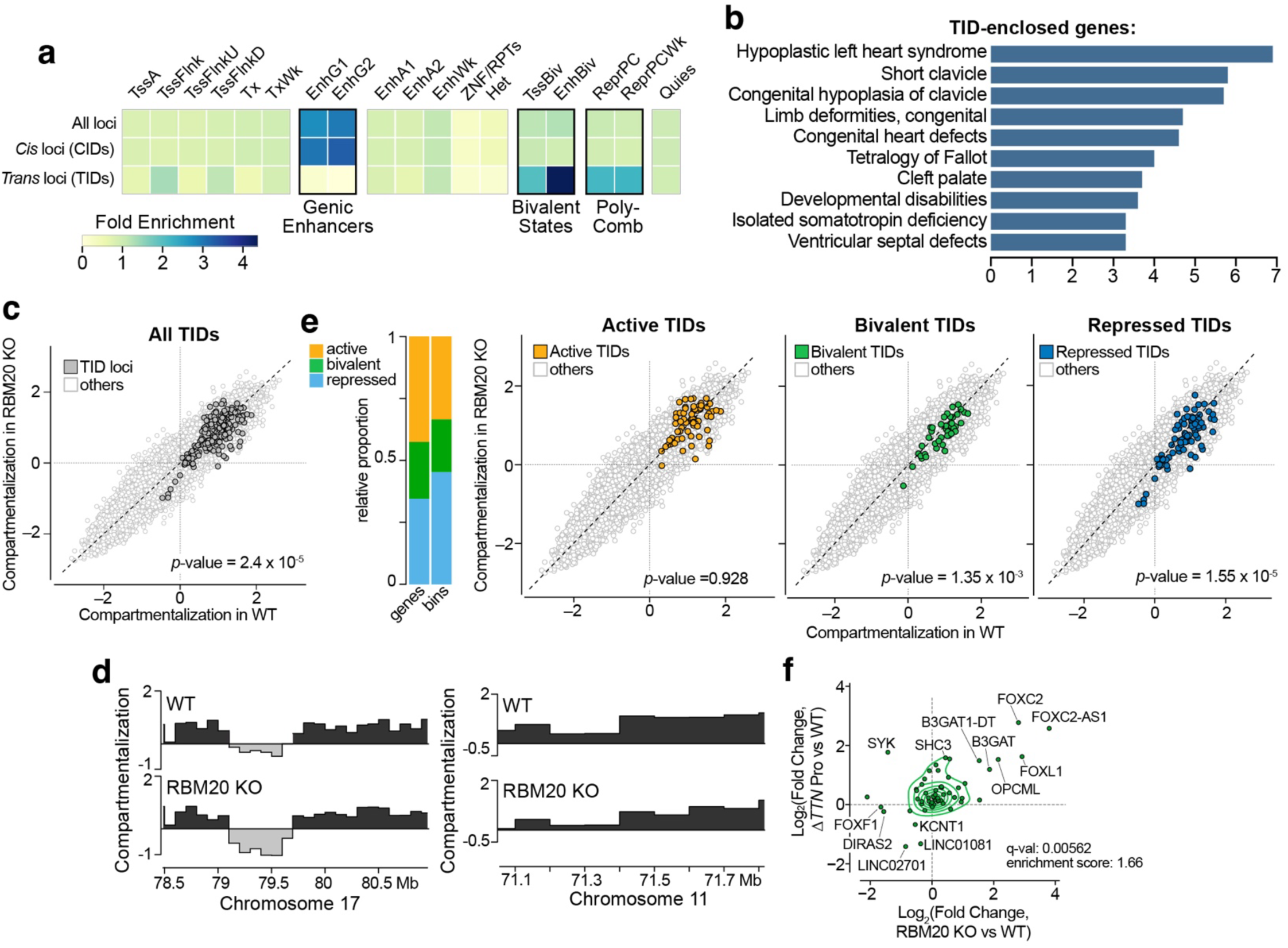
Disruption of *TTN* RNA factory integrity modulates TID chromatin architecture and gene expression programs. **a,** ChromHMM analysis revealed distinct chromatin signatures in pre-*TTN*-associated chromatin domains. Genic enhancers were enriched in *cis*-interacting domains (CIDs), whereas bivalent- and polycomb-repressed domains were enriched in *trans*-interacting domains (TIDs). **b,** Gene Ontology (GO) analysis^56^ of TID-enclosed genes. The top ten disease pathways enriched by DisGeNET are shown, highlighting prominent enrichment of cardiac disorders. **c–d,** Loss of RBM20 induces mild, though pervasive, shifts in TID compartmental character. **c,** Scatter plot comparing DNase Hi-C eigenvectors in WT and RBM20 KO CMs, for all genomic bins, excluding chromosome 2. TID bins exhibit a significant downward shift upon RBM20 loss. *p*-value: two-sided Wilcoxon signed-rank test. **d,** Examples of two TIDs exhibiting increased B-type (*left*) or diminished A-type (*right*) compartmentalization in RBM20 KO cells. **e,** TID loci with active, bivalent, and repressed chromatin signatures in WT CMs exhibit distinct responses to RBM20 loss. *Left*: distribution of TID bins, classified according to epigenetic status. *Right*: same data as in (**c**), highlighting the behavior of each TID class. Only bivalent and repressive TIDs exhibit significant compartmental shifts. **f,** Genes in bivalent TID domains undergo concordant transcriptional shifts upon loss of RBM20 and of the *TTN* transcript (by promoter deletion, "Δ*TTN* Pro"). Enrichment score represents the fraction of TID-enclosed genes that are upregulated in both RBM20 KO and ΔTTN Pro cells relative to WT, normalized to the genome-wide fraction of genes undergoing the same concordant upregulation. q-value: Fisher’s two-sided test. See also **Supplementary** Fig. 5c–**d**.

To explore this hypothesis, we examined how the compartmental character and gene expression of TID loci change upon loss of RBM20. While CID compartmentalization was largely invariant between WT and RBM20 KO hESC-CMs (*p* = 0.491; **Supplementary Fig. 4d**), TID loci collectively exhibited a modest but significant lowering of A-compartment character upon RBM20 loss (**Fig. 3c,d**; average difference in Hi-C eigenvectors, RBM20 KO–WT = –0.105; *p* = 2.4 x 10^-5^; two-sided Wilcoxon signed-rank test). This shift was primarily driven by a weakening of A-character in WT-defined A-compartment TIDs, and strengthening B-character among WT-defined B-compartment TIDs, with relatively few TID loci undergoing complete transitions between compartments (3.53% of 100 kb TID bins; **Fig. 3d**). We noted that the magnitude of this effect varied according to the epigenetic classification of each TID in WT cells. Using ChromHMM annotations, we grouped TIDs into sets with predominantly activating (EnhA1, EnhA2, Tx, TssA), bivalent (EnhBiv, and TssBiv), and repressive states (Het, Quies, ReprPCWk, ReprPC; **Fig. 3e***, left*), which exhibited correspondingly high, intermediate, and low aggregate transcription levels in WT CMs from multiple genetic backgrounds (**Supplementary Fig. 4e**–**g**). Notably, while the compartmentalization of active TID loci was unchanged between WT and RBM20 KO CMs (average Δeigenvector = –0.0153; *p =* 0.928, Wilcoxon signed-rank test), bivalent and repressive TIDs showed significant shifts towards more B-compartmental character (average Δeigenvector = –0.1278 and –0.1616; *p* = 1.35x10^-3^ and 1.55x10^-5^, respectively; Wilcoxon signed-rank test, **Fig. 3e***, right*). This modest though widespread shift in compartmental character was accompanied by correspondingly subtle changes in gene expression. These were most apparent in bivalent TIDs, where 13.3% and 10% of enclosed genes underwent at least two-fold transcriptional up- or down-regulation, respectively, including several transcription factors with established roles in cardiac development^65-67^ (**Supplementary Fig. 4c, f–g**). In contrast, alterations to pre-mRNA splicing were markedly more pronounced, as discussed below. These findings are consistent with RBM20 contributing to *TTN* RNA factory organization and the compartmental character of TID loci. Alternatively, these effects may arise indirectly through RBM20’s established roles in pre-mRNA splicing, including regulation of *TTN* itself, rather than from disruption of factory architecture *per se*.

To distinguish between these models, we sought an orthogonal strategy to disrupt factory architecture independently of RBM20 depletion. To this end, we used cardiomyocytes derived from an established hESC line that lacks the *TTN* promoter (*ΔTTN* Pro, **Fig. 1c**). This mutation abolishes production of the nascent RNA scaffold on which the factory assembles^12^, thereby preventing formation of prominent RBM20 puncta while maintaining RBM20 expression at near-WT levels (**Supplementary Fig. 5a**–**b**). Remarkably, in Δ*TTN* Pro cells, a substantial subset of bivalent TID genes exhibited transcriptional dysregulation similar to that observed upon RBM20 loss (20% upregulated at least two-fold; **Fig. 3f** and **Supplementary Fig. 5d**; q-value comparing (Δ*TTN* Pro/WT) to (RBM20 KO/WT) = 0.0056; Fisher’s two-sided test). These cells likewise recapitulated many of the splicing defects observed in RBM20 KO cells, as detailed below. Importantly, these effects were substantially attenuated in control *TTN* knockout (*TTN* KO) cells, which express nascent *TTN* transcripts but lack functional protein due to a premature stop codon that disrupts translation but not mRNA stability^12^ (**Supplementary Fig. 5a, e**; q-value = 0.273; Fisher’s two-sided test). This serves as an important tool to distinguish *TTN* RNA-dependent effects (*i.e.*, specific to the Δ*TTN* Pro line) from those caused by loss of Titin protein and the resulting changes in cardiac contractility (also observed in this line)^12^. We note that these results do not exclude additional effects of RBM20 loss from contributing to the broader gene-expression phenotype—including the downstream consequences of altered pre-mRNA splicing on transcription factors and other key genetic regulators. However, our findings implicate the *TTN* RNA factory, jointly organized by RBM20 and nascent *TTN* RNA, in fine-tuning the compartmental character and transcriptional output of TID-associated genes, particularly in bivalent domains.

### The TTN RNA factory is a node of post-transcriptional RNA compartmentalization

Many classes of nuclear bodies (*e.g.,* Cajal bodies, paraspeckles, nuclear stress bodies) function in part by compartmentalizing and co-regulating specific populations of target transcripts^7^. Inspired by this, and guided by our initial hypothesis that the *TTN* RNA factory serves as a hub of RBM20-mediated splicing, we sought to characterize the factory’s transcriptome and its remodeling upon RBM20 loss. To achieve this, we applied pre-*TTN* O-MAP-Seq^41^ in both WT and RBM20 KO hESC-CMs (**Fig. 4a**). This analysis identified hundreds of RNAs that co-compartmentalize with nascent *TTN* in each cell type with significant though modest enrichment over their respective input (average fold–enrichment, O-MAP/input: 2.21 (WT), 2.32 (RBM20 KO); **Supplementary Table 4**). That *TTN* associates with a defined cohort of transcripts in RBM20 KO cells contrasts sharply with our O-MAP-ChIP results, wherein loss of RBM20 abolished the *TTN* pre-mRNA*’*s affiliation with chromatin beyond its own locus (**Fig. 2**). Since we expect that co-transcriptionally localized transcripts would exhibit enrichment at both the RNA and chromatin levels, this suggests that *TTN* does not solely compartmentalize with transcripts encoded at nearby genomic loci, but that its RNA interactions are, at least in part, programmed independently of chromosomal proximity. Indeed, few *TTN*-proximal transcripts were encoded by either CID or TID loci (∼7.2% and 0.6%, respectively, in WT cells; 4.7% and 0.6% in RBM20 KO cells; **Fig. 4b**), by any locus on chromosome 2 (∼7% in WT; 4.7% in RBM20 KO), or by loci previously defined as RBM20 targets (∼0.25% and 0.62% of enriched transcripts, representing ∼13.3% and 8.8% of established RBM20 target loci^29^; **Supplementary Fig. 6a,b**). Collectively, these data, together with the prominent enrichment of exons, relative to introns, suggest that most transcripts captured by O-MAP were recruited to the *TTN* RNA factory post-transcriptionally, and may represent a distinct set of interactions from those discovered previously^12^.

**Figure 4.**
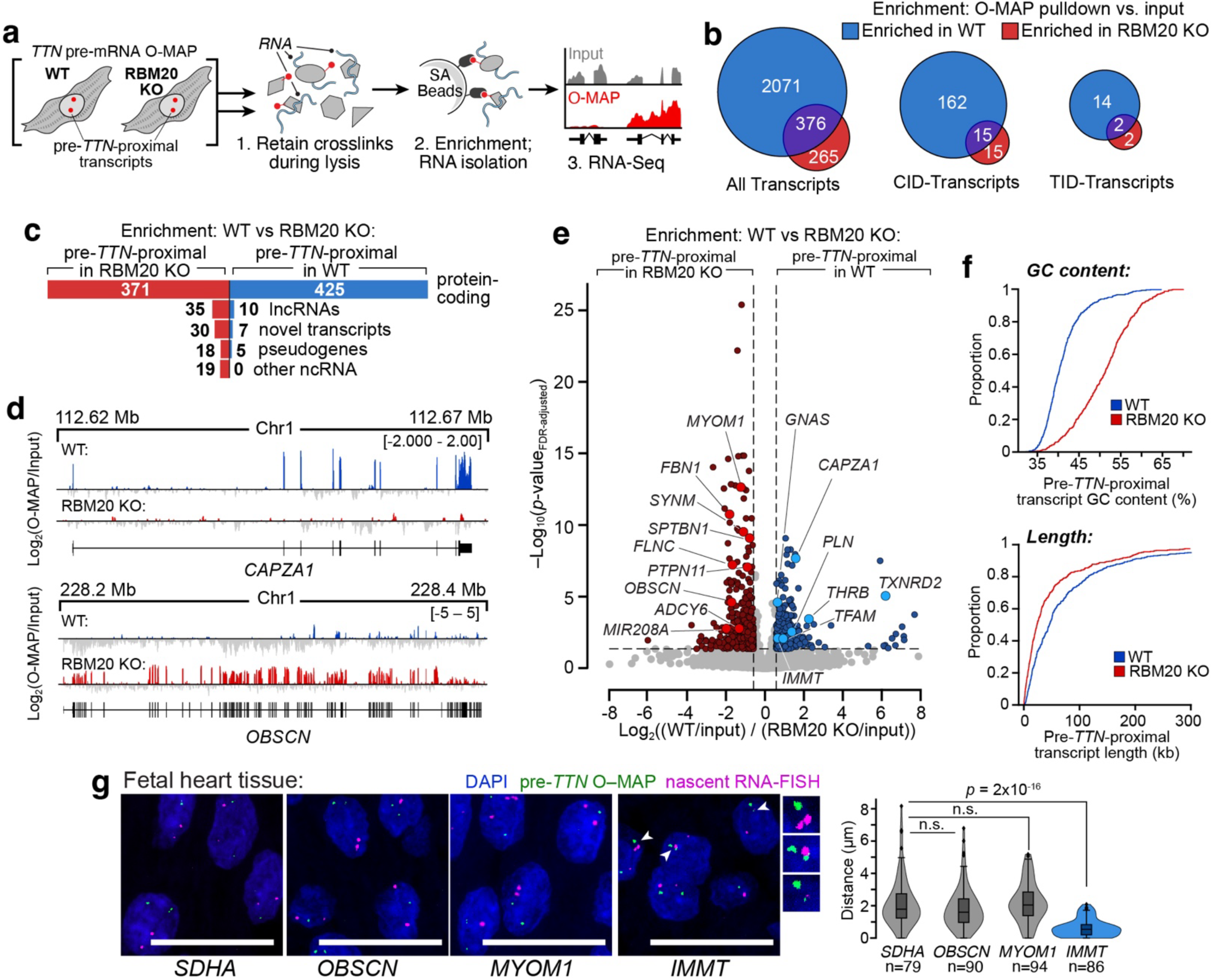
The *TTN* RNA factory compartmentalizes hundreds of mRNAs post-transcriptionally. **a,** Experimental scheme. Transcripts within *TTN* RNA factories were mapped by O-MAP-seq^41^, applied in parallel to WT and RBM20 KO CMs. As in O-MAP-ChIP (Fig. 2), transcripts were enriched by pulling down the biotinylated proteins to which they are bound and quantified by high-throughput sequencing. **b,** Most pre-*TTN*-proximal transcripts are encoded from loci outside the *TTN* RNA factory. Venn diagrams summarizing all (*left*), CID-enclosed (*middle*), and TID-enclosed (*right*) enriched transcripts, in both WT and RBM20 KO cells. Enrichment was measured relative to matched samples of whole-cell RNA. **c**, Overview of the pre-*TTN*-proximal transcriptome. Most transcripts are protein-coding, irrespective of RBM20 status. **d**, Examples of pre-*TTN*-proximal transcripts specifically enriched in WT (*top*) and RBM20 KO (*bottom*) CMs. *CAPZA1* and *OBSCN* are known RBM20 targets^45^. **e,** Volcano plot showing differential enrichment of pre-*TTN*-proximal transcripts in WT and RBM20 KO cells (*n*=4 biological replicates). Notable WT-specific transcripts encoding Ca^2+^–handling and metabolic proteins are highlighted in blue; RBM20 KO-specific transcripts encoding proteins implicated in DCM etiology are highlighted in red. **f,** Cumulative Distribution Function (CDF) plots of the nucleotide composition (*top*) and length (*bottom*) of pre-*TTN-*proximal transcripts. RBM20 KO-specific transcripts have markedly higher GC content and are modestly shorter than their WT-specific counterparts. **g**, *IMMT*, but not *MYOM1* or *OBSCN*, co-compartmentalizes with pre-*TTN* in fetal heart tissue. pre-*TTN* visualized by O-MAP Imaging^41^; other transcripts were visualized by RNA-FISH; quantified data are plotted on the right. Internal box and whisker plots indicate the median, 25^th^, and 75^th^ percentiles, and the 5th–95th percentile range. Data pooled from ≥ 3 biological replicates. *p*-values: Kruskal-Wallis test followed by Dunn’s multiple comparisons vs. the *SDHA* negative controls. n.s.: *p* > 0.05; ****: *p* < 0.001. All scale bars: 20 µm; zoomed images are ∼2.5x.

We next examined how the *TTN*-proximal transcriptome is altered upon RBM20 loss. We compared the O-MAP-enriched transcripts from WT and RBM20 KO hESC-CMs, with each dataset normalized against its corresponding input (i.e., WT/input vs. KO/input). This analysis identified 447 and 473 RNAs that preferentially localize near nascent *TTN* in each cell type (**Fig. 4c**). These differentially localized transcripts were predominantly coding, though RBM20 KO cells exhibited increased enrichment of several classes of noncoding RNA (collectively ∼22% of RBM20 KO-specific transcripts, *vs*. ∼5% in WT, **Fig. 4c**). Notably, only a minority of RBM20 target transcripts (mapped previously by eCLIP)^22^ were differentially enriched in either cell type (**Supplementary Fig. 6b,c**), with rare examples observed in both WT and RBM20 KO cells (*e.g.,* **Fig. 4d**). Likewise, TID-encoded genes were not differentially enriched in either cell type (**Supplementary Fig. 6c**). Collectively, these data suggest that RBM20-dependent remodeling of the pre-*TTN*-proximal transcriptome is not determined primarily by direct RBM20 binding or by changes in local chromatin architecture but instead reflects other determinants of RNA localization. Indeed, transcripts’ differential localization exhibited clear correlations with both length and sequence composition, as pre-*TTN*-proximal RNAs were markedly longer (by ∼34 kb) and AU-rich (∼10% lower GC-content) in WT hESC-CMs than in RBM20 KO cells (**Fig. 4f**), with statistically significant differences in GC-content spanning all transcript regions—most prominently in exons, introns, and 3′–UTRs (**Supplementary Fig. 6d**). These structural preferences may reflect differential recruitment by co-compartmentalized RNA-binding proteins.

We next sought to experimentally validate that O-MAP-enriched transcripts compartmentalize within *TTN* RNA factories by applying a two-color imaging approach, combining SABER-FISH^53^ to visualize candidate *TTN-* proximal transcripts with either intron- or exon-targeting probes (labeling nascent and mature transcripts, respectively; **Supplementary Table 1**) with O-MAP-Imaging^41,68^ to visualize nascent *TTN.* These orthogonal imaging experiments largely confirmed the predictions of our O-MAP-Seq data. For example, nascent *IMMT* preferentially compartmentalized with nascent *TTN* in WT CMs, whereas nascent *MYOM1* and *OBSCN* did so preferentially in RBM20 KO CMs (**Supplementary Fig. 7a**). These findings were recapitulated in fetal human heart tissue—selected to match the early cardiac developmental trajectory of our *in vitro* differentiated cardiomyocyte lines^12^—suggesting that the interactions discovered in our cultured cardiomyocyte model system are also formed *in vivo* (**Fig. 4g**). Mature transcript pools also exhibited the same overall trend, wherein the puncta nearest to nascent *TTN* displayed the same genotype-specific localization patterns (**Supplementary Fig. 7b**). In all cases, transcripts compartmentalized within ∼0.5–1.4 µm of nascent *TTN*, similar to those observed in our O-MAP *trans*-interacting chromatin domains (**Supplementary Fig. 2**), and comparable in size to many nuclear bodies (*e.g.*, nuclear speckles)^11^. These findings provide orthogonal support that O-MAP-Seq enrichment corresponds to altered single-cell RNA proximity and illustrate O-MAP’s use for probing compartment-level associations.

### RBM20 loss rewires pre-mRNA splicing programs at the TTN RNA factory and beyond

The selective localization of transcripts to the *TTN* RNA factory prompted us to examine whether this compartment is associated with distinct alternative splicing programs, as several classes of nuclear bodies have been implicated in shaping isoform choice or selectively compartmentalizing specific transcript species^41,69-71^. We therefore asked two related questions: do transcripts residing within the *TTN* RNA factory exhibit distinct splicing behavior, and does disruption of factory architecture reshape splicing programs more broadly?

Alternative splicing events were modestly enriched among genotype-selective *TTN*-proximal transcripts,defined as the differential O-MAP/input comparison in **Supplementary Fig. 6** and **Supplementary Table 4**. Nearly 20% of WT- and RBM20 KO-specific O-MAP-enriched RNAs exhibited differential splicing— compared to ∼15% of transcripts genome-wide—driven predominantly by skipped exon (SE), mutually exclusive exon (MXE), and retained intron (RI) events (*p =* 2.6 x 10^-5^; two-sided *z*-test; **Fig. 5a,b**). Notably, approximately 60% of alternative splicing events in genotype-selective *TTN*-proximal RNAs occurred in the RBM20 KO-selective set (**Fig. 5c**, *left*), with retained intron (RI) events showing the strongest bias (85% unique to the knockout line; **Fig. 5c**, *right*). These RBM20-dependent differences were also evident in splicing effect sizes, defined as differences in Percent Spliced In (ΔPSIs), since RBM20 KO-specific, pre-*TTN-*proximal transcripts included some of the most pronounced splicing changes observed (0.974%, compared to average ΔPSIs of 0.731% for WT-specific, factory-resident transcripts, and 0.915–0.945% for non-resident transcripts overall, **Fig. 5d** and **Supplementary Fig. 8a**–**c**). Together, these data indicate that RBM20 loss disproportionately reshapes the splicing landscape of *TTN*-proximal transcripts, even though most of these transcripts are not direct RBM20 targets (**Supplementary Figs. 6a** and **8b**). A rare exception is *IMMT*, a WT-specific *TTN-*proximal transcript in which RBM20 loss dysregulates a SE event coincident with its depletion from the factory (**Fig. 5e**). This splicing defect was recapitulated in WTC-11 induced pluripotent stem cell-derived cardiomyocytes (hiPSC-CMs) bearing the DCM-associated RBM20 R636S mutation, indicating that the observed remodeling of the pre-*TTN*-proximal transcriptome extends beyond our RUES2-derived model (**Supplementary Fig. 8d** and **Supplementary Table 5**). Collectively, these data suggest that loss of RBM20 not only reshapes the composition of the *TTN* RNA factory transcriptome (**Fig. 4**), but also shifts it toward an enrichment of mis-spliced RNAs.

**Figure 5.**
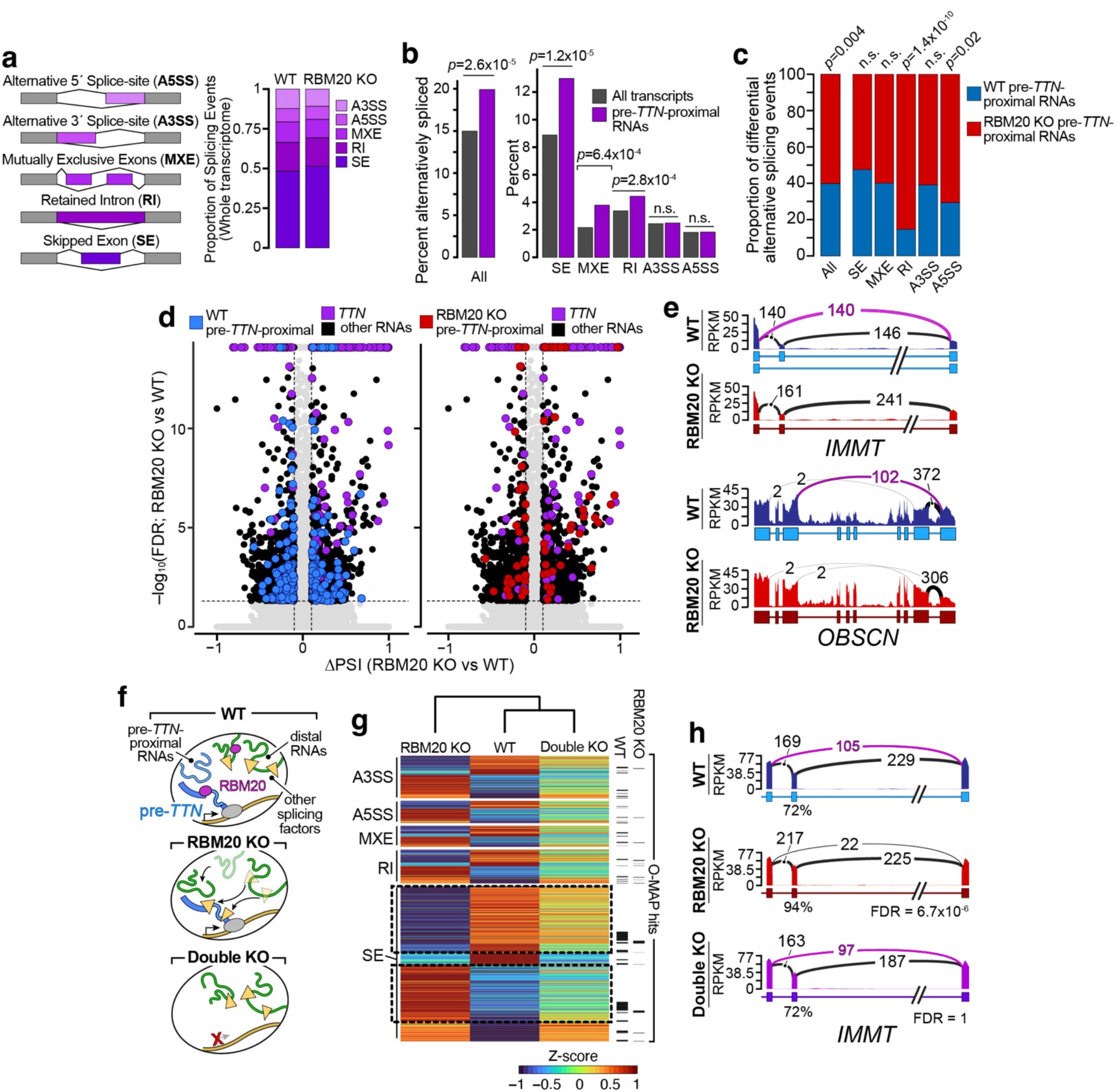
*TTN RNA* factory compartmentalizes and regulates pre-mRNA alternative splicing. **a,** Loss of RBM20 induces widespread splicing dysregulation. *Left:* schematic of major alternative splicing classes. *Right:* distribution of differential splicing events between WT and RBM20 KO CMs, transcriptome wide. **b–c,** Alternative splicing is enriched among pre-*TTN-*proximal transcripts. **b,** Differential splicing between WT and RBM20 KO CMs, quantified for all transcripts (*grey*) and pre-*TTN-*proximal transcripts (*purple*). Pre-*TTN-*proximal transcripts exhibit increased alternative splicing overall (*middle*), driven primarily by SE, MXE, and RI events (*right*; *two-sided z-test).* **c,** Dysregulated splicing is more prevalent in RBM20 KO-selective pre-*TTN*-proximal transcripts (Fig. 4e). Proportions of differential splicing events are shown by event class; *p* values: *two-sided z-test*. **d,** RBM20 KO-specific pre-TTN–proximal RNAs exhibit increased splicing dysregulation. Volcano plots summarizing all significant splicing changes (ΔPSI = (PSI_WT_ – PSI_RBM20 KO_); positive values indicate greater exon inclusion in WT cells). Pre-*TTN*–proximal RNAs and *TTN* are highlighted. **e,** Examples of altered splicing in WT-specific SE event (*IMMT*) and RBM20 KO-specific MXE event (*OBSCN*) in pre-*TTN*-proximal transcripts. Sashimi plots visualizing splicing patterns in each cell type. Splicing events highlighted in magenta were targeted for qRT–PCR validation (**Supplementary** Fig. 8d). **f,** Model: loss of RBM20 exposes pre-*TTN*, which may sequester splicing factors and thereby perturb splicing of other transcripts; depleting the *TTN* pre-mRNA may partially rescue these transcripts. **g,** *TTN* RNA depletion attenuates a subset of splicing changes induced by RBM20 loss. Heatmap of z-score-normalized PSI values for events differentially spliced across WT, RBM20 KO, and Double KO (RBM20 KO/Δ*TTN* Pro) CMs. Events significantly altered between RBM20 KO and WT are displayed and grouped by event type. Pre-*TTN*-proximal hits identified by O-MAP-Seq (**Supplementary Table 4**) are indicated. Highlighted boxes denote SE events rescued in the double knockout (RBM20 KO/ΔTTN Pro) line. **h,** Sashimi plots of the same *IMMT* SE event shown in **f,** demonstrating rescue in the Double KO line. FDRs calculated relative to WT (likelihood-ratio test, rMATS)^72^. Alternative splicing events defined as FDR < 0.05 and |ΔPSI| > 0.1.

We next examined how disruption of *TTN* RNA factory architecture by RBM20 KO reshapes splicing programs across the broader transcriptome, beyond factory-resident RNAs. Globally, we observed 3,477 differential splicing events, with skipped exon (SE) events exhibiting the largest shift (3.15%; **Fig. 5a; Supplementary Fig. 8a**). While this pattern is consistent with RBM20’s established role in promoting exon exclusion^73^, known RBM20-target transcripts^22^ comprised only a minor fraction of SE events (4.05%) and of differential splicing events overall (3.74%; **Supplementary Fig. 8b**–**c**). This suggests that the majority of splicing changes arise from indirect or downstream effects of RBM20 loss, potentially linked to disruption of factory architecture. We hypothesized that this dysregulation may be driven in part by the *TTN* RNA itself. In the absence of RBM20, this long, highly abundant, and aberrantly spliced transcript might serve as a thermodynamic sink for RNA biogenesis factors. This could deplete these factors from their canonical targets and thereby driving widespread missplicing (**Fig. 5h**). A related RNA-binding protein (RBP)-sequestration model has been proposed in repeat-expansion disorders, most notably myotonic dystrophy 1 (Ref. ^74^).

To test this model, we asked whether depletion of the nascent *TTN* transcript could rescue the broader splicing defects induced by loss of RBM20. We used targeted genome editing to delete the *TTN* promoter^12^ in both WT and RBM20 KO WTC-11 hiPSCs, thereby generating a new Δ*TTN* Pro line and a corresponding "Double KO" (RBM20 KO/*ΔTTN* Pro) line. Each line could be efficiently differentiated into cardiomyocytes that exhibited the expected changes in RBM20 expression (or localization) and Titin expression, enabling us to assess how these genetic perturbations reshape global splicing programs (**Supplementary Fig. 9**). Importantly, compared to WT CMs, the Δ*TTN* Pro line showed many splicing changes concordant with those induced by RBM20 loss (R^2^ = 0.357; F-test *p-*value < 2.2 x 10^-16^), with significant correlations observed across all major classes of alternative splicing events (R^2^ = 0.332—0.476 across classes; F-test *p-*values < 2.2 x 10^-16^; **Supplementary Fig. 9d**–**e**). These results were parallelled in our previously described hESC-CM datasets (**Supplementary Fig. 10a–b**). Moreover, only Δ*TTN* Pro hESC-CMs showed a significant enrichment of dysregulated RBM20 targets, confirming that depletion of the *TTN* transcript *per se*—rather than its protein product—more faithfully phenocopies RBM20 loss (**Supplementary Fig. 10**). These observations are consistent with our model that both RBM20 KO and Δ*TTN* Pro perturbations converge on a shared mechanism involving disruption of factory architecture (**Fig. 3f** and **Supplementary Fig. 5**).

Intriguingly, in the Double KO line ablation of the *TTN* transcript attenuated a substantial majority (80.22%) of the splicing defects induced by RBM20 loss, with global splicing patterns clustering more closely with WT CMs than with RBM20 KO cells (**Fig. 5g**). Epistasis analysis revealed an asymmetric interaction between these perturbations, whereby deletion of the *TTN* promoter is epistatic to RBM20 loss: restoring RBM20 in the absence of *TTN* RNA had little effect on global splicing, whereas restoring *TTN* in the RBM20-deficient background induced broad splicing changes (**Supplementary Fig. 11a**–**b**). These effects were observed across all major classes of alternative splicing, but were most pronounced for SE events, which accounted for nearly 56% of rescued splicing events (**Fig. 5g**, *boxed* and **Supplementary Fig 11c**). Importantly, this encompassed numerous WT factory-resident transcripts (110 transcripts; **Fig. 5g**, *right*), including *IMMT*, which exhibited near-complete restoration of its skipped-exon event in the Double KO line (**Fig. 5h**). Analysis of RBP motifs enriched in Double KO-rescued transcripts nominated candidate factors that may explain this phenomenon (**Supplementary Fig. 11c**). Collectively, these findings suggest that the pre-*TTN* transcript is itself a major downstream driver of the widespread splicing defects induced by RBM20 loss.

### The TTN RNA factory proteome couples chromatin organization and RNA biogenesis

Having mapped the *TTN* RNA factory’s chromatin architecture and compartmentalized transcriptome, we next sought to characterize its proteome. To enable this analysis, we re-optimized our mass spectrometry pipeline, O-MAP-MS, originally developed for more abundant RNA targets, to accommodate the inherently restricted signal from pre-*TTN* puncta (**Figs. 1c** and **6a**). Applying this optimized workflow and filtering for proteins with established nuclear localization, we identified 254 and 180 candidate factors enriched in the *TTN-*proximal proteomes of WT and RBM20 KO cardiomyocytes, respectively (**Fig. 6b** and **Supplementary Table 6**). As anticipated, RBM20 was selectively enriched in WT cells. Notably, the WT *TTN-*proximal proteome was dominated by factors involved in chromosome organization and compartmentalization (**Figs. 6b–c**), including nearly all members of the core cohesin complex—the master regulator of chromatin looping and folding^75^—and factors like HMGB2 and HMGB3, which play pervasive roles in chromatin organization^76^. Among the most highly enriched proteins was SAFB (**Fig. 6d**), along with modest enrichment of its homologues, SAFB2 and SLTM— factors originally identified as components of the putative "nuclear matrix," and more recently implicated in RNA-directed chromatin compartmentalization^1,77^. We also observed the prominent enrichment of DNA repair factors—including FEN1, PCNA, and APEX1—and all members of the MCM complex, which plays a "moonlighting" role in chromatin folding by antagonizing cohesin-mediated loop-extrusion (**Fig. 6b,c)**^78^. These chromatin-regulatory proteins were joined by a cluster of RNA-metabolic and RNP-biogenesis proteins, including both factors that play broad roles in general RNA metabolism (*e.g.*, hnRNPAB; EXOSC2), and those that regulate specific cohorts of target transcripts (*e.g.*, SRSF9; QKI; PUM1). Notably, QKI—a developmentally regulated splicing factor with putative roles in mRNA nuclear export^79^—linked to cardiac RNA processing and DCM-associated phenotypes^80^, potentially through its interactions with *TTN*-derived noncoding transcripts^81^. We also observed widespread enrichment of cognate QKI binding motifs within WT-specific factory-resident transcripts, mainly driven by enrichment in introns, with pronounced depletion in the corresponding RBM20 KO-specific transcriptome (**Supplementary Fig. 12**). This suggests potentially coordinated compartmentalization of RBPs and their target transcripts within the WT factory.

**Figure 6.**
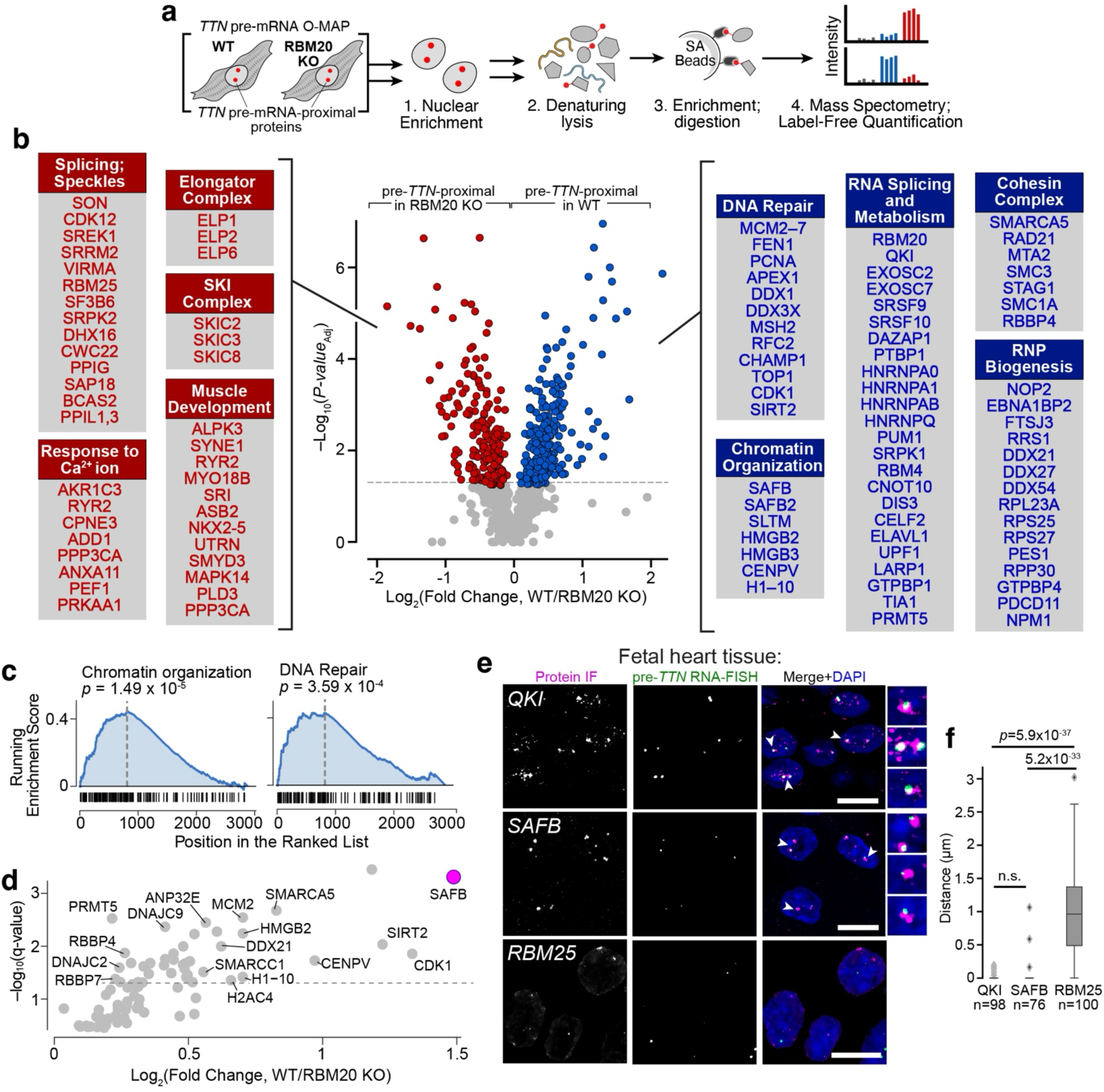
The TTN RNA factory proteome couples chromatin architecture to RNA compartmentalization. ***a,*** Experimental scheme. pre-*TTN* O-MAP was performed in parallel in WT and RBM20 KO CMs. Following nuclear isolation, biotinylated proteins were enriched under denaturing conditions and analyzed by mass spectrometry with label-free quantification (LFQ). **b,** Volcano plot summarizing pre-*TTN* O-MAP-MS hits specific for WT (*blue; right*) and RBM20 KO (*red; left*) CMs. Prominent functional classes and representative protein hits are shown for each. **c,** Gene Set Enrichment Analysis (GSEA) based on Log_2_(Fold change, WT/RBM20 KO) values, highlighting the prominent enrichment of "Chromatin Organization" and "DNA Repair" factors, as defined by Gene Ontology of Biological Processes (GOBP). *p-values:* gene-rank permutation test. **d,** Volcano plot for proteins related to the chromatin organization GOBP term; SAFB highlighted in magenta. Data are the same as plotted in **b. e,** Immunofluorescence for two WT-specific hits, QKI and SAFB, and negative control, RBM25, in WT human fetal heart tissue sections. Nascent *TTN* was visualized by RNA-FISH. Arrowheads denote conspicuous colocalization, highlighted in ∼3x-zoomed images (*right*). All scale bars, 20 μm. **f**, quantification of 3D-distances between pre-TTN and each protein hit. Box and whisker plots indicate median, 25th, and 75th percentile, and the 5th–95th percentile range. Data pooled from ≥ 3 independent biological replicates. p-values by Kruskal-Wallis test followed by Dunn’s multiple comparisons, vs RBM25.

Critically, loss of RBM20 dramatically remodels the pre-*TTN-*proximal proteome (**Fig. 6b**, *left*). In RBM20 KO cells, O-MAP-MS revealed no enrichment of chromatin-architectural factors, consistent with our observation that nascent *TTN* requires RBM20 to maintain chromatin association (**Fig. 2d**). Instead, in the absence of RBM20, the *TTN*-proximal proteome was enriched for a varied assortment of core splicing factors (*e.g.*, the promiscuous splicing regulator RBM25)^82^ and nuclear speckle components (*e.g.*, SON, SRRM2). However, we found no evidence for concordance between these RBPs and the RBM20 KO-specific pre-*TTN*-proximal transcriptome, as observed in WT cells (**Supplementary Fig. 12**). Together, these findings suggest that RBM20 loss erodes the specialized chromatin-associated proteome of the *TTN* RNA factory, shifting it to a speckle-enriched state that may contribute to pathogenicity *via* gain-of-function.

We next sought to validate these hits by combined immunofluorescence and pre-*TTN* RNA-FISH in human fetal heart tissue, focusing on SAFB and QKI. Strikingly, both factors formed prominent nuclear puncta that colocalized with nascent *TTN* transcripts (**Fig. 6e,f**), resembling the pronounced compartmentalization of RBM20 within the *TTN* RNA factory (**Fig. 1d**). In contrast, RBM25 (enriched in RBM20 KO O-MAP) showed no clear colocalization with *TTN* in these samples (**Fig. 6e,f**). The marked enrichment of SAFB and QKI in WT *TTN* RNA factories suggests that these factors may help couple *TTN* mRNA biogenesis to the regulation of gene expression of other factory-resident loci.

To assess the potential role of SAFB and QKI within *TTN* RNA factories, we examined the molecular dependencies governing their recruitment to the compartment and their influence on factory-associated gene expression in cultured CMs. We first asked whether their factory localization required RBM20, nascent *TTN* RNA, or both. In WT hESC-CMs, both SAFB and QKI formed distinct nuclear puncta surrounding nascent *TTN* transcripts, similar to those observed in fetal tissues (**Figs. 7a–b**, *top*). This compartmentalization was strictly RBM20-dependent: loss of RBM20 caused both SAFB and QKI to disperse from nascent *TTN*, yielding a diffuse nucleoplasmic pattern (**Fig. 7a–b**, *bottom; quantified at right*). These results were recapitulated in hiPSC-CMs, where prominent puncta observed in WT cells were replaced by diffuse nucleoplasmic localization upon loss of RBM20 or deletion of the *TTN* promoter (**Fig. 7c–d**). Structured Illumination Microscopy (SIM) further revealed that disruption of factory architecture—by ablating either RBM20 or nascent *TTN* transcripts—dramatically increased the number of SAFB and QKI puncta, yielding more numerous, smaller, and dimmer foci. This effect was nearly equivalent in Δ*TTN* Pro hiPSC-CMs and RBM20 KO cells, and was not rescued in the Double KO line (**Fig. 7c–d**, *right*). Together, these data suggest that recruitment of SAFB and QKI to the factory requires the localized assembly of RBM20 coordinated by nascent *TTN* RNA.

**Figure 7.**
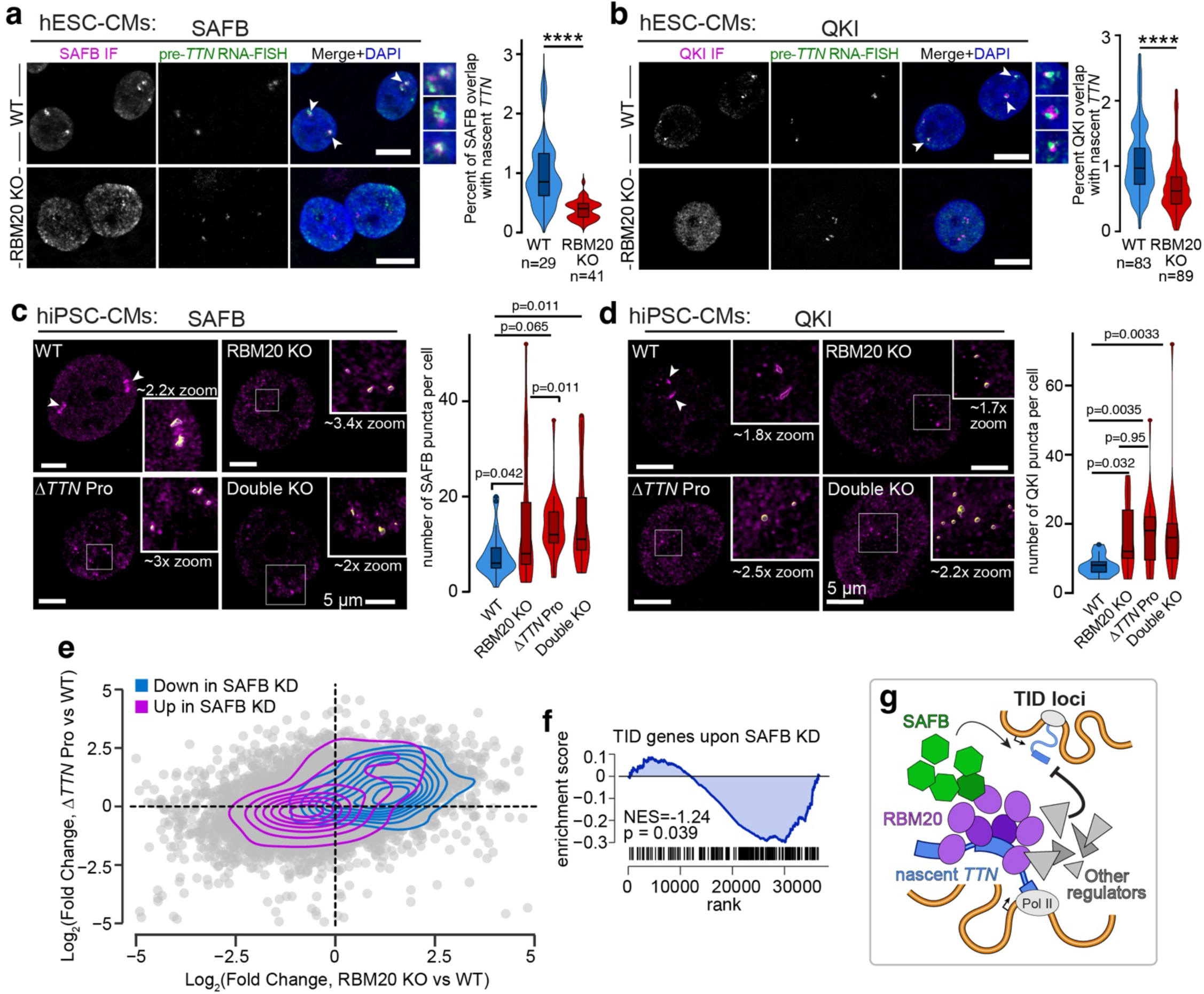
Compartmentalization of SAFB and QKI requires both RBM20 and nascent TTN RNA and regulates TID gene expression. **a–b,** SAFB and QKI concentrate within TTN RNA factories in an RBM20-dependent manner. Immunofluorescence of SAFB (**a**) and QKI (**b**) in hESC-derived CMs. Nascent TTN was visualized by RNA-FISH. Arrowheads denote colocalization, highlighted in ∼2.8x zoomed images (right). Maximum-intensity projections of confocal Z-stacks. Scale bars, 10 µm. The overlap of each protein’s IF signal with nascent TTN is quantified in violin plots to the right. **** p < 0.001, *Welch’s t test.* **c–d**, Structured Illumination Microscopy (SIM) analysis of SAFB (**c**) and QKI (**d**) mislocalization following loss of RBM20, nascent TTN RNA, or both, in hiPSC-CMs. Insets highlight WT factories or residual puncta observed in RBM20 KO, ΔTTN Pro, or Double KO cells (yellow); approximate zoom factors indicated. Right: number of foci per cell, depicted as violin and boxplots, indicating the median, 25th, and 75th percentiles. p-values were computed using a mixed model with differentiation batch as a random effect and genotype as a fixed effect (* if p-value < 0.05). **e,** Gene expression changes following SAFB knockdown show an inverse relationship to those observed upon RBM20 or TTN promoter loss. Scatterplot of the gene expression changes upon orthogonal perturbations of the *TTN* microenvironment (RBM20 KO; *ΔTTN* promoter), viewed as Log_2_(Fold change), relative to WT RUES2 CMs. The distributions of genes significantly upregulated following SAFB knockdown (log_2_(fold change) > 2.0; padj < 0.05) are shown as 2D density plots. **f,** TID gene expression is particularly sensitive to SAFB loss. Gene Set Enrichment Analysis (GSEA) of expression changes in TID-localized genes, relative to a scrambled knockdown control (**Supplementary** Fig. 12). Rightward shift and negative enrichment scores indicate a drop in gene expression upon knockdown. **g,** Model summarizing the role of nascent TTN and RBM20 in maintaining TID bivalency. Nascent *TTN* transcripts nucleate a cluster of RBM20 that in turn recruits SAFB and other chromatin- and RNA-regulatory factors (Fig. 6b), collectively exerting opposing effects on TID gene expression.

Given its prominent enrichment and striking colocalization with nascent *TTN* transcripts, we next sought to determine whether SAFB plays a functional role within *TTN* RNA factories by examining the consequences of its depletion. Using lentiviral short hairpin RNA (shRNA) delivery, we achieved modest SAFB knockdown in WT hESC-CMs (∼2-fold reduction; **Supplementary Fig. 13a–d**). SAFB depletion did not appear to alter QKI localization, suggesting that QKI recruitment is RBM20-dependent but SAFB-independent (**Supplementary Fig. 13e**). As anticipated, SAFB knockdown induced widespread changes in gene expression, with 162 and 101 genes significantly up- or downregulated, respectively (|log2 fold change| > 2 and p_adj._ < 0.05 from limma moderated t-test). Surprisingly, these expression patterns exhibited an inverse relationship with the transcriptional dysregulation observed upon disruption of factory architecture, wherein genes that were upregulated in both RBM20 KO and Δ*TTN* Pro hESC-CMs were downregulated by SAFB depletion (**Fig. 7e**). This relationship was particularly pronounced for TID-localized genes, which were highly enriched among SAFB-regulated transcripts and markedly downregulated upon SAFB knockdown (**Fig. 7f**). These findings suggest that SAFB may act downstream and in opposition to RBM20, promoting the expression of TID-encoded genes and potentially contributing to their bivalent expression signatures (**Fig. 7g**). Notably, SAFB has been implicated in coordinating subnuclear compartmentalization and RNA-directed chromatin organization in several developmental and disease contexts^77,83^. Although it is classically thought to stimulate heterochromatinization, emerging data have demonstrated that SAFB can also associate with nascent transcripts to stimulate gene expression^84^. Our data suggest that it may play a similar role within the *TTN* RNA factory.

Together, these findings establish O-MAP-MS as a powerful approach for resolving the proteomes of single-locus nuclear compartments while uncovering RBM20-dependent regulatory factors—including SAFB— that couple *TTN* RNA biogenesis to local chromatin organization and gene expression.

## DISCUSSION

In this study, we adapted the RNA-targeted proximity-biotinylation platform O-MAP to enable biochemical dissection of discrete subnuclear compartments by targeting the nascent transcripts around which they assemble. This enabled us to map the molecular architecture of the *TTN* RNA factory and propose, to our knowledge, the first molecular model of a single-locus subnuclear compartment (**Fig. 8**, *left*).

**Figure 8.**
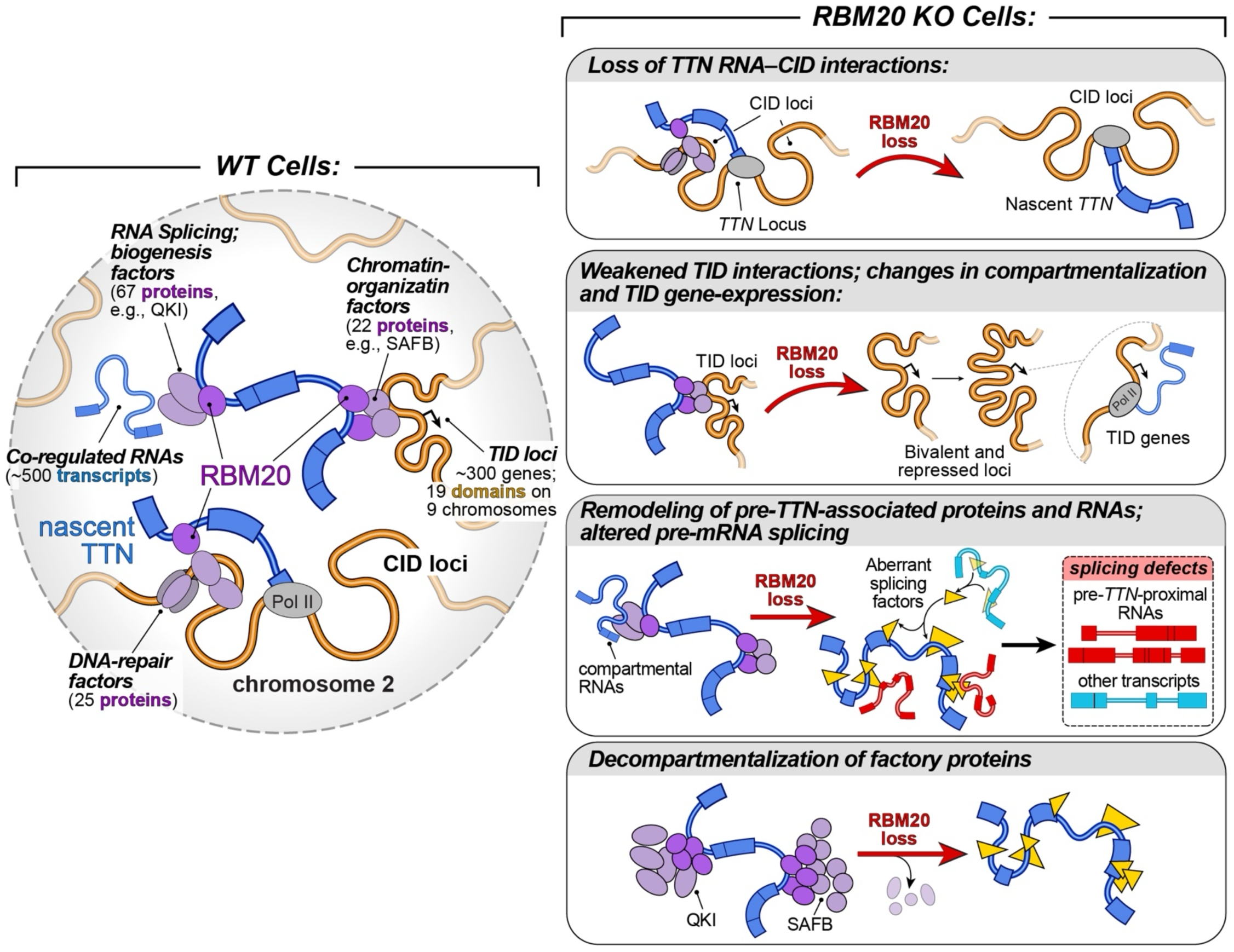
Molecular architecture of the *TTN* RNA factory. *Left*: overview of the compartment-wide interactions discovered by O-MAP. For simplicity, factory components are depicted as directly binding to pre-*TTN* and RBM20, though these may interact through higher-order assemblies or biomolecular condensates. Our data cannot discriminate between interactions with nascent transcripts and incompletely spliced intermediate species, both of which are depicted. *Right*: upon RBM20 loss, *TTN* can no longer remain affiliated with the clique of factory CIDs and TIDs, and associates with an aberrant cohort of transcripts and proteins.

In WT cells, the factory organizes *TTN* mRNA biogenesis within a specialized, cardiomyocyte-specific chromatin neighborhood comprising numerous *cis*-interacting domains (CIDs) on chromosome 2, together with a clique of *trans*-interacting domains (TIDs) interspersed across other chromosomes (**Fig. 2b–d**). These domains assemble along a developmentally regulated trajectory, coalescing throughout the cardiomyocyte lineage and peaking in mature CMs to form a micron-wide compartment centered on nascent *TTN* transcripts (**Fig. 2f** and **Supplementary Fig. 3**). Functionally, this architecture appears to shape the epigenetic status and expression of TID-localized genes, many of which are maintained in a poised or transcriptionally repressed state (**Fig. 3**). Notably, a prominent subset of TID loci bear a bivalent signature—an epigenetic class frequently associated with genes that control stem cell differentiation and cell-fate determination^85^. These loci were especially sensitive to the disruption of factory architecture (**Fig. 3e–f**), suggesting that the factory may help to fine-tune their regulation. Based on our proteomic data, we hypothesize that this regulation is driven by a cohort of chromatin-architectural proteins, including the cohesin complex^75^ and scaffolding factors like HMGB2^76^ and SAFB^77^—each highly enriched within the factory (**Fig. 6b–d**). Consistent with this, depletion of SAFB, which forms prominent factory-localized puncta in WT cells (**Figs. 6e** and **7a,c**), had a pronounced inhibitory effect on TID-localized gene expression (**Fig. 7e**). This suggests that SAFB may play a stimulatory role in the poised expression of TID genes, similar to the transcription-activating functions recently described in embryonic stem cells^84^. In parallel, the factory also appears to serve as an RNA hub that recruits and sequesters a large ensemble of transcripts post-transcriptionally (**Fig. 4** and **Supplementary Fig. 6**), potentially influencing their processing (**Fig. 5** and **Supplementary Figs. 8, 10,** and **11**). We hypothesize that this RNA recruitment may be orchestrated by the dozens of RBPs observed in our proteomic data—including factors like QKI and SAFB, which form prominent factory-localized puncta dependent on both *TTN* expression and RBM20 (**Fig. 7b,d** and **Supplementary Fig. 12**). A similar RNA-compartmentalization phenomenon has been observed in other nuclear bodies—including nuclear speckles^69^, paraspeckles^86^, and nucleoli^87,88^—though its functional significance in the *TTN* factory remains unclear.

Our revised model of *TTN* RNA factory architecture resembles a class of subnuclear compartments termed nuclear stress bodies (nSBs)^7^. Like the *TTN* factory, nSBs are assembled on long, repetitive transcripts and accrete prominent nuclear foci of SAFB. These foci serve as recruitment sites for defined sets of RBPs— many of which are shared by the *TTN* RNA factory (*e.g.*, SRSF1, SRSF9)—which appear to drive post-transcriptional sequestration of many transcripts^7,89^. It is thus possible that the *TTN* RNA factory and nSBs represent two foundational examples of a broader mechanism by which cells assemble functionally distinct chromatin compartments around nascent transcripts. Moreover, the fact that this mechanism can utilize nascent pre-mRNAs like *TTN*, and thereby co-opt a coding gene as a chromatin-regulatory RNA, raises the question of whether additional mRNAs may play similar roles in other developmental and disease contexts.

In the absence of RBM20, nearly every facet of the nascent *TTN* transcript*’s* microenvironment and interactome is remodeled (**Fig. 8**, *right*). In RBM20 KO cells, nascent *TTN* transcripts appear incapable of specifically affiliating with genomic loci beyond the *TTN* locus itself (**Fig. 2d**). Our data strongly suggest that this is not due to a gross defect in chromatin organization, as Hi-C interactions between the *TTN* locus and its clique of CIDs and TIDs remain largely preserved even in the absence of RBM20 (**Fig. 2e** and **Supplementary Figs. 1**, **3**). Instead, we favor a model in which RBM20 is required to retain nascent *TTN* transcripts within the factory and that, upon RBM20 depletion, these “untethered” transcripts associate with a new set of RNAs and proteins, and compartmentalize into an aberrant subnuclear microenvironment—a feature that may be pathogenic.

The untethering of *TTN* from its factory in RBM20 KO cells appears to have subtle, though pervasive effects on the A/B compartmentalization and transcription of TID-localized genes (**Fig. 3d–f** and **Supplementary Figs. 4**–**5**)—epigenetic effects that may partly explain why restoring RBM20 function in differentiated DCM model cells fails to completely rescue disease-associated changes in gene expression^50^. Concomitantly, RBM20-depleted *TTN* transcripts relocalize to a compartment with a distinct proteomic composition that is depleted of core compartmental factors like SAFB and QKI (**Figs. 6b** and **7**). It also co-compartmentalizes with a population of RNAs that are disproportionately mis-spliced, even though few are direct RBM20 targets (**Figs. 5d–e** and **Supplementary Figs. 6, 8**). That these splicing defects—and indeed, a substantial fraction of the global mis-splicing events induced by RBM20 loss—are rescued by further deleting the *TTN* promoter implicates the untethered pre-*TTN* transcript as an active contributor to the downstream splicing phenotype (**Fig. 5f–h** and **Supplementary Figs. 9**–**10**). This model suggests a previously unappreciated pathogenic mechanisms that, in future studies can be explored in more clinically relevant contexts, including models of *bona fide* disease-causal genetic variants. Conceptually, this mechanism resembles a mode of RNA-mediated toxicity proposed in repeat-expansion disorders, which raises the possibility that analogous RNA-driven mechanisms may contribute more broadly to other spliceopathies^90^.

The model emerging from this study significantly enhances our understanding of the molecular architecture and functional scope of the *TTN* RNA factory, and reveals how disruption of this architecture might contribute to RBM-DCM. Prior work, and our own trans-C data (**Fig. 2e**), have demonstrated that the factory organizes a clique of RBM20-target genes, thereby influencing their biogenesis^12,29^. Our results expand on these findings, revealing that the *TTN* factory also organizes hubs of repressed and poised chromatin, as well as post-transcriptional RNA compartmentalization. A deeper investigation into the RNA structural elements and RNA-RBP interactions that drive this compartmentalization may yield new therapeutic avenues for RBM20-DCM—for example, pharmacotherapies that stabilize features of WT factory architecture (*e.g.*, SAFB, QKI), or which antagonize pathological interactions induced by RBM20 depletion (*e.g.*, nascent *TTN* toxicity). Such effects should likewise be pursued in the context model systems that better emulate the etiology of RBM20-DCM. Indeed, while our knockout model has enabled us to cleanly dissect the influences of RBM20 function on factory assembly and maintenance, biallelic complete loss of RBM20 has not been reported as a disease-causing genotype in patients^91,92^. Future studies that apply O-MAP to *bona fide* patient models (*e.g.*, heterozygous R636S point mutants)^91,92^ may disentangle the complex and subtle influences of heterozygous RBM20 dysfunction and toxic gain-of-function effects^22^.

This characterization study also highlights O-MAP’s unique ability to elucidate novel facets of nuclear architecture, establishing a method that complements the DNA proximity-ligation-based tools that are mainstays of the field^2^. Although powerful, these methods are not designed to directly probe the µm-scale, compartment-level interactions that define the *TTN* RNA factory^29^ and many other subnuclear compartments^1,6^, and cannot capture compartmental RNAs or proteins^6^. By comparison, nascent transcript O-MAP has enabled us to capture higher-order TID interactions (**Fig. 2**), as well as the ensembles of co-compartmentalized RNAs (**Fig. 4**) and proteins (**Fig. 6**), revealing a host of novel interactions and regulatory phenomena. Our approach also spatially refines compartment-level analysis in a way that is unattainable with established biochemical methods, such as fractionation^93^, affinity purification of tagged compartmental proteins^34^, or targeted *in situ* proximity-biotinylation of these proteins^4^. By targeting the common biophysical or biochemical features of a given compartment type, these methods survey all such compartments simultaneously and therefore cannot identify the unique molecular features that define individual subnuclear neighborhoods. O-MAP overcomes this longstanding technical roadblock, enabling us to characterize, for the first time, the distinct molecular microenvironment surrounding a single genomic locus of interest. However, several important limitations should be considered. First, because O-MAP labels molecules over length scales of tens of nanometers^44^, it resolves compartment-level molecular organization rather than direct (Å-scale) molecular interactions. This enrichment cannot be converted into a precise physical distance: in a heterogeneous population, enrichment can arise if a subset of target molecules/cells enters the labeling radius, whereas failure to enrich does not define a specific separation threshold (**Supplementary Discussion**). Consequently, the proteins, RNAs, and genomic loci identified by O-MAP should be interpreted as constituents of a shared molecular microenvironment rather than direct binding partners. Second, successful application of O-MAP depends on the availability of a uniquely defined, targetable RNA species that faithfully marks the compartment of interest. Transcriptionally silent loci and highly repetitive genomic regions may therefore be better suited to complementary approaches such as DNA O-MAP^94^. Likewise, because RNA biogenesis can produce multiple transcript populations that bear the same targeted sequence yet reside in distinct subnuclear microenvironments, care must be taken to ensure that the targeted RNA uniquely identifies the compartment of interest. Despite these limitations, we anticipate that our nascent transcript O-MAP strategy can be applied to dissect nuclear architecture at the single-neighborhood level across a wide range of biological settings.

## Supporting information

Compiled Supplementary Materials

## DATA AVAILABILITY AND ACCESSION CODES

Mass spectrometry data have been deposited to the MassIVE repository with the accession number MSV000096046, and to PRIDE, with the accession number PXD056627. RNA- and DNA-pulldown sequencing data have been deposited to the Gene Expression Omnibus under SuperSeries accession number GSE234811. Bulk RNA-sequencing data have been deposited to ArrayExpress under the accessions E-MTAB-XXX (hESC-CMs KO data), E-MTAB-XXX (hiPSCs-CMs KO data), and E-MTAB-XXX (hESC-CMs KD data). Hi-C raw sequencing data have been deposited to ENA under the Studies Report accession PRJEB104212. Intermediate processed sequencing data are available on Zenodo under the 18722155 record. Raw imaging files are available from the corresponding authors upon request. This manuscript made use of public datasets from ENCODE, accession numbers ENCSR584MHY and ENCFF135PCO.

## CODE AVAILABILITY

Settings used for routine computational analyses (*e.g.* high-throughput sequencing alignment and quantification; Hi-C analyses) are described in detail in (**Methods**). Custom code and Dockerized workflows used for the analyses reported here are available on the Bertero and Shechner Labs’ GitHub repositories: (1) https://github.com/SheqLab/ (2) https://github.com/sara-bianchi/TTN_O_MAP_data_analysis

## ACKNOWLEDGEMENTS

We thank A. Tsue for technical assistance with O-MAP, H. Reinecke and J. Klaiman for assistance with cardiomyocyte differentiation, N. Peters and members of J. Scott’s and B. Beliveau’s laboratory for assistance with imaging, S. Henikoff for the generous donation of reagents, members of Y. Sancak’s lab for their mitochondrial expertise, I. Ulitsky for insight into O-MAP-Seq data analysis, E. Van Nostrand and L. Kuriyan for their insight into QKI targeting, and members of the Shechner, Bertero, and Murry labs for their support, thoughtful advice, and critiques. This work was supported by National Institutes of Health Grants R01HL160825 (to C.E.M., and D.M.S.), R01GM138799 (to E.E.K. and D.M.S.), T32GM007750 (to E.E.K.), UM1HG011531 (to B.H., and W.S.N.), R35GM137916 (to C.K.C. and B.J.B.), and the Canadian Institutes of Health Research Project Grant PJT 173542 and the NSERC Discovery Grant RGPIN-2020-04180, to M.M. and P.G.M. D.M.S. was further supported by a Chan-Zuckerberg Institute Collaborative Pairs award and McKnight Foundation Neurobiology of Brain Disorders award. E.E.K. was further supported by the NSF DEB2016186 and AHA 902616; A.B. received funding from the European Research Council (ERC) under the European Union’s Horizon Europe research and innovation programme (Grant agreement No. 101076026; Project Acronym TRANS-3); C.E.M also was supported by a grant from the Robert B. McMillen Foundation. D.C. was supported by grants from the Italian Ministry of Health (Piano Operativo Salute Traiettoria 3, T3-AN-09, “Genomed”; Ricerca Finalizzata 2021, “genOMICA”; MCNT2 2023, “EUCARDIS”). Imaging at the UW Keck Center was supported by NIH S10 OD016240 and the UW Student Technology Fee (STF). This work used an EASY-nLC1200 UHPLC and Thermo Scientific Orbitrap Fusion Lumos Tribrid mass spectrometer purchased with funding from a NIH S10OD021502. NGS data analysis was facilitated through the use of advanced computational, storage, and networking infrastructure provided by the Hyak supercomputer system and funded by the STF at the University of Washington, and supported by the UW Student Technology Fund.

## AUTHOR CONTRIBUTIONS

E.E.K. contributed to experimental design, performed O-MAP experiments and hit validation, and oversaw initial data analysis, assisted by S.L. and N.S. S.B. oversaw most computational analyses, assisted by Q.L., B.H, and W.S.N. *Signature* analyses were performed by M.M. and P.G.M. Cardiomyocyte differentiation was overseen by A.F., A.B., C.E.M, and S.M. M.C., L.T., and E.W. generated and characterized new knockout hiPSC-CM lines. E.B. and D.C. generated RNA-Seq and Hi-C data. D.M.M., R.F., D.K.S., and S-E.O. assisted with mass-spectrometry experiments. C.K.C. and B.J.B. assisted with DNA-FISH experiments and image analysis. D.M.S., A.B., and C.E.M. initially conceived of this work and obtained research funding; D.M.S. and A.B. directed research. All authors contributed to writing.

## CORRESPONDING AUTHORS

Correspondence to: Charles Murry; Alessandro Bertero; David M. Shechner

## MATERIALS AND METHODS

### Cell culture and tissue sections

RUES2 hESCs (RUESe002-A; WiCell) and WTC-11 hiPSCs (a gift of Bruce R. Conklin; also available from Coriell GM25256) were maintained on Matrigel (Corning; 354277) in Essential 8 media (Thermo Fisher; A1517001). Media was exchanged every other day. Cells were passaged with Versene (Thermo Fisher; 15040066) into low confluent wells supplemented with 10 µM Y-27632 (Stem Cell Technologies; 72302).

hESC lines were differentiated into cardiomyocytes (hESC-CMs) as described previously^12,22^. Briefly, 4.5–9 x 10^4^ cells were plated onto 12-well plates pre-coated with 80 µg/µl growth factor-reduced Matrigel in mTeSR1 (StemCell Technologies; 85850) and 10 µM Y-27632, for 24 hours. Cells were then treated with 1 μM CHIR-99021 (Cayman Chemical; 13122). On day 0, media was exchanged for RPMI-1640 with glutamine supplemented with 500 μg/ml Bovine Serum Albumin (BSA), 213 μg/ml ascorbic acid (RBA media; Millipore Sigma), and 5 μM CHIR-99021, and maintained for 48 h. On day 2, culture media was replaced with RBA media supplemented with 2 μM WNT-C59 (Selleck Chemicals; S7037) and maintained for 48 h. On day 4, media was replaced with RBA. On day 6, cells were cultured in RPMI supplemented with B27 (Thermo Fisher; 17504044), with media exchanges every two days. To increase cell viability after cryopreservation and thawing, on day 13 cells underwent a 45-minute heat shock at 42 °C, using pre-warmed media. On day 14, cells were dissociated with TrypLE (Thermo Fisher; 12563011) with gentle trituration and were strained with 100 µm filters (Fisher Scientific; 22363549). Dissociated cells were frozen in CryoStor media (1 million cells/100 μL CryoStor; Millipore Sigma; C2874) and frozen slowly at -80 °C using a Mr. Frosty Freezing Container (Thermo Fisher; 5100-0001). The next day cells were stored in liquid nitrogen.

hiPSC lines were differentiated into cardiomyocytes using a similar protocol. Briefly, hiPSCs were seeded at a density of 1.5–3 × 10⁵ cells per well in Geltrex-coated 12-well plates in hE8 medium supplemented with 2 µM thiazovivin. After 24 h, cells were primed with basal medium containing 1 µM CHIR99021 before initiating differentiation (day 0) with RBA medium supplemented with 5 µM CHIR99021. Subsequent patterning with WNT-C59 (2 µM, day 2), RBA refresh (day 4), and maintenance in RPMI + B27 from day 6 onward were performed as described above. Between days 12–15, cardiomyocytes were enriched by sodium lactate metabolic selection, followed by a 5-day recovery period in RPMI + B27 prior to replating and downstream analyses.

Formaldehyde-fixed and paraffin embedded fetal heart tissue sections were acquired from the National Disease Research exchange. The University of Washington Pathology Research Services Laboratory was used to cut blocks into 5 µm sections.

### Generating mutant hiPSC-CM cell lines

WTC-11 hiPSCs containing GCaMP6f (Gladstone Institutes) were used in all experiments. An RBM20 KO sub-line from the same genotype was generated previously by TALEN-mediated deletion of 8 bp in RBM20 exon 9 (Ref. ^95^). To generate the Δ*TTN* Pro line, we removed the 1.5-kbp long region containing the promoter and first *TTN* exon, via CRISPR-Cas9 using the following sgRNA targeting sequences: upstream 5′-GTGAAACTCTACTTAGAGGG–3′; downstream 5′–TTTAAGGGAATCAACTGCTG–3′. These sequences were cloned into plasmids containing Cas9 and mCherry expressed from the same promoter. The same editing was performed in parallel on both "WT" WTC-11 GCaMP6F and RBM20 KO sub-lines, to generate Δ*TTN* Pro and Double KO cell lines, respectively.

Cells at 50-60% confluency were treated for 4 hours with E8 medium (Thermo Fisher Scientific; A1517001) supplemented with 2 µM Thiazovivin, washed briefly with magnesium- and calcium-free PBS and harvested with Accutase (Thermo Fisher Scientific; 00-4555-56). After suspending in growth medium, cells were counted and 1 x 10^6^ cells were suspended in P3 Primary Cell solution containing Supplement 1 (Lonza; V4XP-3024) and 10 µg of DNA. Cells were electroporated using the CA 137 program in the 4D Nucleofector X Unit (Lonza; AAF-1003X) and seeded in a well of a 24-well plate in the E8 medium containing CEPT supplement: 50 nM Chroman1 (MedChem Express; HY-15392), 5 µM Emricasan (Selleckchem; S7775), Polyamine supplement 1:1000 (Sigma-Aldrich; P8483), and 0.7 µM Trans-ISRIB (Tocris; 5284). The medium was changed to fresh E8 (without CEPT) after 24 hours.

Cells were detached using TryplE:0.5 mM EDTA solution (3:1 by volume) and passed through a 40 µm cell strainer (Pluriselect; 43-10100-40). Individual mCherry-positive cells were isolated by sorting using an SH800S Sony cell Sorter equipped with a 100 µm Chip, using a 3-drop setting, and seeded onto Matrigel-coated 96-well plates, containing E8 + CEPT + 1% Penicillin/Streptomycin (Euroclone; ECB3001D). After 3 days, the medium was changed to E8 + 1% Penicillin/Streptomycin, with subsequent daily media changes. The resulting colonies were assayed by genomic PCR probing the targeted region (**Supplementary Fig. 9**) using the following primers: Forward 5′–CTTTGGGATGAACAACTTTCTGTGT–3′; Reverse 5′–ATGCCATCTATGGGGGTGGAA–3′. Successfully edited cell lines were verified for genomic integrity by low resolution karyotyping by Illumina microarray (Life & Brain company; Bonn, Germany). SNP data were analyzed for copy number variations in GenomeStudio 2.0.5 with the cnvPartition 3.2.1 plugin, using default parameters except for minimal probe count of 6 and Confidence Threshold of 70. CNVs were then filtered for confidence threshold (above 85) and the size of the CNV (above 350 kb, or above 100 kb known CNV hotspots in hPSCs^96^, and compared to the profile observed in low passage number WTC-11 cells. No novel CNVs were identified on any of the gene-edited sublines.

### pre-TTN O-MAP main protocol

RUES2-CMs were used for both O-MAP omics and O-MAP imaging. For omics, CM stocks were thawed onto 12-well plates coated in Matrigel at 1.2 million cells/well in RPMI/B27, supplemented with 10 µM Y-27632 and 10% FBS. For imaging, 2.5 x 10^5^ cells were plated onto 10mm glass bottom plates (Cellvis; D35-10-1.5-N). The next day media was exchanged for RPMI/B27 and exchanged every two days thereafter, until post-differentiation day 21. The core O-MAP protocol follows the same two-day protocol as previously described^41^, but with slight optimizations for cardiomyocytes. Briefly, during the fixation step, formaldehyde concentration was increased to 4% (v/v; increased from the standard 2%), and endogenous peroxidase inactivation proceeded for 20 minutes (increased from the standard 10 minutes). The remainder of the protocol is largely unchanged from that described previously^41^, and is summarized here.

### pre-TTN O-MAP day 1

Cells were quickly washed with 1x DPBS (Thermo Fisher; 14190250) and fixed in 4% (v/v) formaldehyde (Electron Microscopy Sciences; 15710) in 1x PBS (Sigma; 6506) for 10 minutes at room temperature, without agitation. The fixation solution was aspirated and quenched by two five-minute washes with 250 mM glycine in 1x PBS, with rocking (3 rpm on a platform rocker; Everlast Rocker 247). Cells were then briefly washed with DPBS and permeabilized with 0.5% (v/v) triton-X 100 (Thermo Fisher; 28314) in PBS for 10 min with rocking. Three five-minute DPBS washes were then performed to prevent over-permeabilization. To inactivate endogenous peroxidases, samples were then incubated with 0.5% H_2_O_2_ (v/v, in PBS) for 20 minutes with rocking. Two five-minute washes with DPBS followed to prevent H_2_O_2_ damage to cells. Cells were then equilibrated in Formamide Wash Buffer (40% (v/v) deionized formamide (VWR, 0314); 2X SSC (Thermo Fisher; AM9765); 0.1% Tween-20 (Thermo Fisher; 28320) for five minutes with rocking. Wash buffer was then aspirated, and to each well was added 50 µL of Primary Probe Mix, comprising 0.1 µM aggregated primary oligo probe pool (**Supplementary Table 1**), in 1x Hybridization Buffer (40% (v/v) deionized formamide; 2x SSC; 0.1% (v/v) Tween-20; 10% (w/v) dextran sulfate (SIGMA; D8906); in nuclease-free water). A clean 12-well glass coverslip was then applied to spread the Primary Probe Mix across the samples. A kimwipe soaked in nuclease-free water was placed between the wells, and plates were sealed with Parafilm to create a hybridization chamber. Plates were then incubated for 8 hours at 42°C without rocking.

### pre-TTN O-MAP day 2

After primary hybridization, ∼1 mL pre-warmed 30% Formamide Wash Buffer was added to the samples. Coverslips were then removed, and cells were allowed to wash for five minutes with rocking. Two additional five-minute washes with fresh 30% Formamide Wash Buffer followed. To each well was added 50 µL of O-MAP Secondary Probe Mix, comprising 0.1 µM HRP-oligo conjugate (**Supplementary Table 1**) in 30% Hybridization Buffer. Samples were covered with a clean coverslip and incubated for one hour at 37°C in the dark. For pre-*TTN* O-MAP imaging, endogenous biotin blocking was performed (see "*pre-TTN O-MAP Imaging*," below). Otherwise, to each well 1 mL of PBST (0.1% (v/v) Tween-20, in 1X PBS) was added. Coverslips were then removed, and samples washed for 15 minutes in PBST at room temperature, with gentle rocking. Samples were washed three times in PBST (15 minutes each), and PBST was aspirated. To initiate *in situ* biotinylation, samples were treated with 1 mL of Labeling Solution (0.8 μM biotinyl-tyramide (Sigma; SML2135); 1 mM H_2_O_2_, in 1x PBST) for 45 minutes at room temperature, protected from light. The biotinylation reaction was quenched by the addition of 1 mL of PBST-q (10 nM sodium azide (Sigma; 71289), 10 nM sodium ascorbate (Sigma; A4034) in 1X PBST), for three five-minute washes, with gentle rocking.

### pre-TTN O-MAP Imaging

After secondary hybridization with HRP-conjugated oligo, samples were washed three times in PBST; five minutes per wash. To block the signal from endogenous biotin, samples were incubated for 30 minutes with 1 mL 1% nuclease-free BSA (VWR; 97061-420) in PBST, at room temperature with rocking, followed by a 15-minute incubation with 1 mL Neutravidin solution (10 μg/mL neutravidin (Thermo Fisher; 31000), 1% (w/v) nuclease-free BSA, in 1x PBST), at room temperature with rocking. After three five-minute washes in PBST, samples were incubated with 1 mL of biotin solution (10 μg/mL D-biotin (Thermo Fisher; B20656) in 1x PBST) to saturate neutravidin. Samples were then washed three times in PBST before proceeding with *in situ* biotinylation and quenching as described above. Following biotinylation, samples were incubated with 1 mL of 1 µg/mL neutravidin-Dy550 (Thermo Fisher; 84606) in 1% BSA PBST for one hour at room temperature with rocking. Samples were then washed three times with PBST, stained with DAPI, and either imaged immediately or mounted in Vectashield (Vector Labs; H-1900-10) and stored at 4°C before use.

### Combined pre-TTN O-MAP, RNA-FISH, and immunofluorescence

In combined O-MAP/RNA-FISH experiments (*e.g.*, **Fig. 1d**) primary probes were split into two sub-pools, equipped with orthogonal "landing pad" sequences that enable simultaneous recruitment of a fluorescent imaging oligo (for RNA-FISH) and HRP-conjugate (for O-MAP). These were introduced in a single primary hybridization step. Cells were processed as described in ***pre-*TTN *O-MAP day 1***, followed by endogenous biotin-blocking and *in situ* biotinylation, as described in ***pre-*TTN *O-MAP imaging***, above. After quenching biotinylation, samples were equilibrated 30% Formamide Wash buffer (1 mL, three five-minute washes, Room temperature, with rocking). This buffer was aspirated and each well was treating with 50 µL of FISH secondary probe mix, comprising 0.1 µM Alexa Fluor 647-conjugated secondary oligo (IDT; **Supplementary Table 1**) in 30% Formamide Hybridization Buffer. Samples were covered with a clean coverslip, incubated for one hour at 37°C in the dark, then washed three times with PBST. Before proceeding to immunofluorescence, samples were blocked with 4–5% (w/v) nuclease-free BSA in 1X PBST for 30 minutes. Samples were then incubated with rabbit-RBM20 (Thermo Fisher; PA5-57404, used at 1:800 dilution), or other primary antibodies listed below, in 4–5% BSA in 1X PBST for 1 hour 45 minutes (RBM20) or overnight (all others). Samples were washed three times in 1X PBST (5 minutes per wash), and then incubated with Atto488-conjugated goat anti-rabbit (Thermo Fisher; R37116, used at 1:300 dilution), Alexa Fluor 568-conjugated anti-rabbit (Thermo Fisher; A-11011) or Alexa Fluor 488-conjugated anti-mouse, (Thermo Fisher; A-11001) in 1% BSA in 1XPBST, supplemented with 1 µg/mL neutravidin-Dy550, where appropriate. Samples were washed four times in 1 x PBST, stained with DAPI, and imaged immediately in 1 x PBST.

When performing pairwise fluorescent imaging (*e.g.*, combined O-MAP/RNA-FISH; O-MAP /immunofluorescence, RNA-FISH/immunofluorescence) the O-MAP protocol preceded the RNA-FISH or immunofluorescence protocol and RNA-FISH preceded the immunofluorescence protocol. Antibodies used: rabbit-TTN (Myomedix; Z1Z2, used at 1:250 dilution); rabbit-QKI (Sigma; HPA019123, used at 1:100–1:125 dilution); rabbit-SAFB (Sigma; HPA020076, used at 1:50–1:100 dilution); rabbit-H3K4Me3 (Active Motif; 39159, used at 1:500 dilution); mouse anti-sarcomeric α-actinin (Abcam, ab9465; used at 1:100 dilution).

### pre-TTN O-MAP-ChIP

O-MAP-ChIP was performed essentially as previously described^41^, but with 24 sonication cycles during the cell-lysis/chromatin-shearing step. Briefly, O-MAP-labeled cells (3.6 x 10^6^ per replicate; two biological replicates per experimental condition) were harvested in PBSTq using cell scrapers, pelleted at 3,000 x g for 10 minutes at 4°C, and supernatant was aspirated. Cell pellets were flash-frozen in liquid nitrogen and stored at –80°C until use. Samples were thawed on ice and resuspended by gentle pipetting in 1mL CLB (20 mM Tris pH 8.0, 85 mM KCl, 0.5% (v/v) NP-40) supplemented with 1x Halt EDTA-Free protease inhibitor cocktail (Thermo Fisher; 87786) and 10 mM sodium azide, for 10 minutes. Lysates were clarified by centrifugation at 3,000 x g for five minutes at 4 °C, and supernatant was aspirated. Pellets were resuspended in fresh CLB, pelleted and aspirated twice more, for a total of three washes. Pellets were then lysed by gentle pipetting in 1 mL of NLB (10 mM Tris-HCl pH 7.5, 1% (v/v) NP-40, 0.5% (w/v) sodium Deoxycholate, 0.1% (w/v) SDS) supplemented with 1x Halt EDTA-Free protease inhibitor cocktail and 10 mM sodium azide, for 10 minutes. Lysates were then placed into an ice-cold thermal block and sheared with a Branson Digital Sonifier outfitted with a double stepped microtip, at 10-12 Watts over 30 s intervals (0.7 s on; 1.3 s off), with 30 s resting steps between intervals. A total of 24 intervals were performed per sample, and after every 5 intervals the thermal block was placed on ice for 2 minutes to maintain temperature. This resulted in an average shearing size of approximately 200–400 bp, as gauged by agarose gel electrophoresis.

Sheared samples were clarified by centrifugation at 15,000 x g for 10 minutes at 4°C, and supernatants were moved to fresh tubes. 10 µL was removed from each sample and saved as "input," and to the remaining was added 100 µL slurry of streptavidin-coated magnetic beads (Thermo Fisher; 88817), that had been pre-equilibrated by two washes in NLB. Samples rotated end-over-end for 2 hours at room-temperature. Beads were collected by magnetic separation, aspirated, and washed with the following series of buffers (1 mL each, 5-minutes per wash, at room-temperature, with end-over-end rotation): (1) NLB, supplemented with 5 mM EDTA, 10 mM sodium azide and protease inhibitors (1x Halt EDTA-free Protease Inhibitor Cocktail) and 150 mM NaCl; (2) NLB, supplemented with 5 mM EDTA, 10mM sodium azide and protease inhibitors, (3–4) two washes in High Salt Buffer (1 M KCl, 10 mM Tris-HCl pH 7.5, 5 mM EDTA), (5–6) two washes in Urea Buffer (2 M Urea, 10 mM Tris-HCl pH 7.5, 5 mM EDTA), (7) 10 mM Tris-HCl pH 7.5, 1% (w/v) SDS, (8–9) two washes in TE Buffer (10 mM Tris-HCl pH 7.5, 1 mM EDTA). Washed beads were resuspended in 100 µL of Elution Buffer (2% (v/v) N-lauryl sarcosine (VWR; 100218-364), 10 mM EDTA, 5 mM DTT, in 1x PBS, supplemented with 200 μg proteinase K (Thermo Fisher; AM2548)). Samples were then incubated at 65°C with shaking at 700 rpm for one hour in a Mixer HC, transferred to 0.2 mL tubes, and incubated overnight at 65°C in a thermocycler. Samples were transferred to 1.5 mL tubes and extracted by addition of an equal volume of phenol pH 6.6, followed by two equal volume extractions with absolute chloroform. Samples were supplemented with 1 µg GlycoBlue (Thermo Fisher; AM9516) and NaCl to 300 mM final, and precipitated with ethanol, overnight at –20°C. Samples were then pelleted by centrifugation at 15,000 x g for 30 minutes at 4 °C, washed twice with ice-cold 80% ethanol and resuspended in 20 µL nuclease-free water. Residual RNA was removed by supplementation of 10 µg RNase Cocktail Enzyme Mix (Thermo Fisher; AM2286) and incubation at 37 °C for 1 hour. Finally, DNA was purified by phenol extraction and ethanol precipitation as described above, and resuspended in 20 µL of nuclease-free water.

### Pre-TTN O-MAP-ChIP library preparation and sequencing

DNA samples were processed for sequencing and sequenced as previously described^41^. Briefly, samples were quantified using a NanoDrop One and 300 ng of DNA from each sample was used for library preparation, using the NEBNext Ultra II DNA Library Prep Kit and NEBNext Multiplex Oligos for Illumina (NEB; E7645 and E7335, respectively), according to the manufacturer’s instructions. Two biological replicates were used per condition, and each library was assigned a unique index. Library concentrations were quantified using the NEBNext Library Quantification Kit for Illumina (NEB; E7360), and independently validated using an Agilent 4200 TapeStation with a “DNA High Sensitivity” kit (Fred Hutch Genomics Core). Libraries were pooled in equimolar concentrations to 20 nM aggregate in nuclease-free water, with no more than eight libraries per pool. Each pool was subjected to 150 cycles of paired-end sequencing on an Illumina HiSeq 4000, run in high output mode (Azenta Life Sciences).

### pre-TTN O-MAP-ChIP data analysis

Deep sequencing reads were trimmed using TrimGalore! (https://www.bioizformatics.babraham.ac.uk/projects/trim_galore/) with parameters -q 30 --phred33, and mapped to the GRCh38 genome using Bowtie2 version 2.4.47 (Ref.^97^). Duplicate reads were removed with the Picard MarkDuplicates function (http://broadinstitute.github.io/picard). Coverage bigWig files were generated using deepTools version 3.5.19 (Ref. ^98^) with a binSize of 1 and normalizing with RPKM. Log_2_(Fold Change) bigWig files were generated by comparing O-MAP ChIP data to matched input samples. Regions of enrichment were called using Enriched Domain Detector (EDD)^99^, using default settings and blacklisting regions from the ENCODE Blacklist^100^.

To visualize CIDs and TIDs, data were plotted as log_2_(fold change, pulldown/input), averaged across two replicates, using the pyGenomeTracks module^101^. The number of bins for each plot was selected according to the size of the visualization range, often using 10 kb binning on the whole chromosome 2, and 1 kb binning for individual loci. ChIP tracks of individual samples were converted from the bigWig to the BedGraph format using the “bigWigToBedGraph” module (ucsc-bedgraphtobigwig Bioconda package), and imported into R for subsequent analyses, focusing on positive peaks. The genome was binned at 1 Mb and the number of positive peaks was evaluationed for each bin, after removing peaks also observed in mock O-MAP datasets. The distribution of positive peaks per bin was plotted as a density plot and a maximum accepted threshold of 100 peaks was selected. The density of positive peaks in the remaining bins of the pre-*TTN* O-MAP samples was plotted with the “genomicDensity” function and the positions of the EDD domains were plotted with the “genomicRect” function (circos R package)^102^.

### DNase Hi-C library preparation and data collection

DNase Hi-C libraries were generated broadly according to the *in-situ* DNase Hi-C protocol of Ramani *et al.*<u>^103^</u>, with minor modifications. Briefly, 2–5×10^6^ cells were crosslinked in 2% formaldehyde for 10 min, quenched with glycine, washed in ice-cold PBS, and collected. Samples were lysed in NP-40-containing buffer, followed by Dounce homogenization to obtain intact nuclei. Nuclei were then conditioned in 0.5× DNase I digestion buffer containing 0.2% SDS, followed by Triton X-100/RNase A treatment, and chromatin was fragmented *in situ* with DNase I. Digestion was standardized to 4 U DNase I for 7 min at room temperature to limit release of free DNA while preserving efficient fragmentation. Pre- and post-digestion aliquots were de-crosslinked and assessed by PAGE, and only preparations showing minimal pre-existing shearing and a bulk fragment distribution below ∼1 kb were processed further. Intact nuclei were immobilized on carboxylated magnetic beads and processed sequentially by *in situ* end repair, dA-tailing and overnight ligation of pre-annealed biotinylated bridge adaptors, followed by phosphorylation and dilute *in situ* proximity ligation. To monitor library quality during the workflow, supernatants were collected after each enzymatic step were retained as nuclear-integrity controls, and an end-repair/dA-tail positive-control ligation was processed in parallel to verify efficient end processing. After reversal of crosslinks, proteinase K digestion and DNA recovery by isopropanol precipitation and bead cleanup, DNA was sonicated before streptavidin capture using a Bioruptor Pico (3 cycles, 30 s on/90 s off in 100 µl), yielding material predominantly in the ∼200–400-bp range. Biotinylated ligation products were then isolated on streptavidin beads and subjected on-bead to end repair, dA-tailing and Illumina Y-adaptor ligation. Libraries were amplified only after empirical cycle optimization using trial PCRs (10–14 cycles), and for high-complexity libraries all remaining bead-bound material was used for the final amplification. Two biological replicates per condition were sequenced on the DNBSEQ-T7 platform (MGI) using PE100 v3 flow cells with 100-bp paired-end reads (R1, 100 cycles; R2, 100 cycles) and dual 8-bp indexes (I1, 8 cycles; I2, 8 cycles), targeting ∼1 billion read pairs per library.

### Hi-C data analysis

Raw reads were demultiplexed with bcl2fastq v2.20.0, concatenated, and aligned to hg38 genome (UCSC) with HiC-pro^104^ (v3.1.0). Since the Hi-C library fragmentation was obtained using DNase digestion, the parameters GENOME_FRAGMENT and LIGATION_SITE were left empty. The hg38 *GENOME_SIZE* was downloaded from the UCSC genome browser and manually edited to maintain autosomes only. BIN_SIZE was set to 100kb, 150kb and 1 Mb; all other parameters were used with default settings.

Differential compartment analyses were performed with dcHiC^105^, using ICE-normalized matrices at 100 kb and 150 kb resolution, the corresponding bed files, and normalization-biases files as inputs. The top eigenvectors were identified from *cis* interaction maps using the dchicf.r R script, with “pcatype” set to cis and “PC” set to 3. The best eigenvector for each chromosome was then selected with “pcatype” in *select* mode, and differential compartments analysis between WT and RBM20 KO samples was performed by setting “pcatype” to *analyze*. Compartment tracks and -log_10_(*p*-value) tracks were visualized with the pyGenomeTracks module. Compartmental differences at CIDs and TIDs were evaluated separately, after isolating bins on chromosome 2 for the CIDs and for the remaining chromosomes for the TIDs. Chromosome 21 was excluded from analyses because a principal component that captured A/B compartmentalization could not be identified amidst the background signal from long- and short-arm separation. For all other chromosomes, eigenvalue distributions for both CID and TID bins (defined using WT O-MAP ChIP data, defined using EDD^99^) were compared between the WT and RBM20 KO conditions using the “wilcox.test” function (stats R package) with “paired” set to *TRUE* and “alternative” set to *two.sided*. Data were visualized using the ggplot2 R package.

To correlate Hi-C data from RUES2 CMs at different developmental stages (**Supplementary Fig. 1**), we used data from (Ref. 12), accessed via the 4DN Data Portal (Experiment Set codes 4DNESGTHHJAC, 4DNES6G787KF, 4DNESSU7RRNR, 4DNESW72J62T) and pre-processed at 150 kb resolution as described above. Compartmental Pearson correlations were then computed from the selected eigenvector tracks using the “rcorr” function (Hmisc R package). Stratum-Adjusted Correlation Coefficient (SCC) analysis^52^ was performed on *cis* interaction maps (150kb resolution) after converting the ICE normalized matrices from the HiC-pro format to the cool format with the python module hicConvertFormat.py (HiCExplorer 3.4.1)^106-108^. Cool matrices were imported into R with the “cool2matrix” function (HiCrep R package) and SCC scores measured using the “get.scc” function (HiCrep R package) with all the settings set to default. Both correlation analyses were visualized using the “pheatmap” R function (pheatmap R package).

Due to the high degree of interreplicate reproducibility, raw contact maps were combined using python modules in HiCExplorer. First, each matrix (1 Mb resolution) was converted to h5 format using the hicConvertFormat.py module. Replicates were then combined based on genotype and converted to mcool format using the hicSumMatrices.py, and hicConvertFormat.py modules, respectively. Combined mcool matrices were then converted back to the HiC-pro format with the preprocess.py module (dcHiC) and used as input with the corresponding bed files for HiC-pro normalization, with the “step” parameter set to ice_norm and other parameters set to default. The resulting combined, ICE normalized matrices were then converted to mcool format with hicConvertFormat.py.

Virtual 4C analysis was performed using the “virtual4C” function in the HiContacts R package^109^, using a Granges object of the hg38 genome coordinate centered on the *TTN* Promoter (chr2:178806793–178808363). The same analysis was performed on both the merged mcool ICE normalized matrices (1 Mb resolution) and on the matrices of individual samples (100 kb resolution), following conversion to the HiC-pro format in mcool using the hicConvertFormat.py module^101^. The resulting merged, virtual 4C tracks were then normalized genome-wide to the total number of *TTN* interactions within each sample, scaled by 10^6^, and exported as bed files using the pyGenomeTracks module.

To compare HiC data, pre-*TTN* O-MAP-ChIP and mock O-MAP-ChIP (**Supplementary Fig. 1**), all datasets were converted to genomic signal tracks and aggregated into 100-kb bins across chromosome 2. For each bin, the mean interaction value was calculated using the bedmap function in Bedops v2.4.41. The resulting binned tracks were imported to R and their correlation with the compartment tracks and virtual 4C tracks on chromosome 2 (each at 100 kb resolution) was evaluated with the “rcorr” function. The resulting correlations were visualized with the “pheatmap” function.

### Trans-C

Trans-C analyses^29^ were performed on raw merged HiC mcool matrices at 1 Mb resolution. First, each matrix was normalized with the following functions in the base.py module: “ice_normalize_matrix” to perform ICE normalization, “normalize_chr_territories” to normalize over the distances of different chromosomal territories, and “donut_filter” to enhance the signal of isolated interactions over the background. The normalized matrices were then used as inputs for the “do_random_walk” function (rand_walk.py module) setting “alpha” to 0.5 and using the bin containing the *TTN* gene as seed. To generate the background random model, the same analysis was repeated 1000 times, using a different starting seed at each iteration. The resulting ordered lists of bins were then imported to R for further processing. Circular plots (**Fig. 2e**) were built using the “genomicLink” function (circos R package), coloring each bin based on its ranking in *TTN*-seeded random walk. TID-interactors were selected as the central bin of each TID, while the interactors in the RBM20 target list were selected as the bins containing the transcription start sites (TSSs) of genes in established RBM20-target loci^12^. AUROC analyses (**Supplementary Fig. 3**) were performed with the “roc.curve” function (PRROC R package)^110^ using the above lists of O-MAP/EDD peaks and established RBM20 target loci as positive bins, and all other bins as negatives. The same analytical pipeline was applied to all random model data to obtain null distributions of AUC values. Statistical significance was then empirically calculated via a permutation test, counting how many times the AUC of the random model exceeded that obtained using the *TTN* seed, and dividing by 1000.

### Signature analysis and permutation testing

We applied the *Signature* pipeline^31^ at 1 Mb resolution for *trans*-chromosomal proximities, and at 50 kb for *cis-* chromosomal proximities, respectively, to 11 independent Hi-C datasets derived from a time-course across cardiomyogenesis^57^, from aorta, bladder, psoas, left and right ventricles^58,59^, and from WT and RBM20 KO cells. We quantified genome-wide *cis*- and *trans*-chromosomal interaction strengths using statistical significance estimates, as previously described^31^. We treated the *TTN* locus (chr2:178,000,000–179,000,000; hg38) as a fixed anchor bin and extracted its *trans*-chromosomal proximities with the predefined set of TID bins (n = 33) and with all significant proximities along chromosome 2. To assess whether the *TTN*–TID proximity subset exhibits stronger interaction weights than expected relative to the *cis-* and *trans*-chromosomal background of the *TTN* locus, we performed a permutation-based enrichment test. Specifically, for each dataset, we randomly sampled the *cis-* and *trans*-chromosomal target bins while keeping the *TTN* anchor bin fixed. For each permutation, we computed the mean *z*-score of the *TTN* proximities with the randomly selected bins. This procedure was repeated 10,000 times to generate a null distribution of mean interaction strength. An empirical one-sided *p*-value was calculated as the proportion of permutations in which the mean *z*-score of the random bin set was equal to or greater than the observed mean *z*-score of the *TTN*–TID set. This framework tests whether the *TTN*–TID group shows significant enrichment in interaction strength compared to a random expectation, conditioned on the same anchor locus.

### DNA SABER-FISH Probe Design

DNA SABER-FISH^53^ probes (**Supplementary Table 3**) were designed using the online PaintSHOP application^111^, with a target pool size of 1000 probes per locus. Briefly, segments of DNA from the center of each target locus (see, "*pre-TTN O-MAP-ChIP data analysis*," below) were uploaded to PaintSHOP as a .BED file. DNA Probe Design parameters were as follows: Repeat=allowed; Max Off Target Score=100; Max K-mer count=5; Min Prob Value=0; Optimize Set=show; Number of probes per target=1000, Enable=off. For the Append Sequences function the following parameters were used: Design Scheme=DNA Probe Design (5′ Outer Primer Sequence=Append; Orientation=Forward; Format=Same for all probes; Select Sequence Set=Custom; ATCCTAGCCCATACGGCAATG), (5′ Bridge Sequence=Append; Orientation=Forward, Format=Unique for each target; Select Sequence Set=Kishi et al. 2019 Bridges); (5′ Inner Primer Sequence=Append, Orientation=Forward; Format=Same for all probes; Select Sequence Set=Custom; TGAATAGCAGCGGTGGCAAAC); (3′ Appending Scheme=Primer/Bridge/Universal); (3′ Outer Primer Sequence=Append; Orientation=Reverse Complement; Format=Same for all probes; Select Sequence Set=Custom; GTATCGTGCAAGGGTGAATGC).

### DNA-FISH Probe Amplification

Probes were purchased from Twist Bioscience as a single pool (Oligo pool, Tier 3L ssDNA, >6,001—12,000 oligos (151-200nt)), and Z resuspended in Tris-EDTA, to 6.0 ng/µL final. The probes were amplified by PCR, using Phusion HF (Thermo Fisher; F531S) and primers Forward 5′–ATCCTAGCCCATACGGCAATG–3′ and Reverse 5′–TAATACGACTCACTATAGGGGTATCGTGCAAGGGTGAATGC–3′, which adds a T7 promoter sequence. This reaction ran on a ProFlex PCR System (Thermo Fisher) using the following parameters: Initial denaturation: 98°C, 5s; Denaturation: 98°C, 5s; Annealing: 58°C, 30s; Extension: 72°C, 15s; Final extension: 72°C, 5 min. Denaturation, annealing, and extension were performed for 10 cycles. DNA products were then purified using the Monarch PCR and DNA Cleanup Kit (NEB; T1030S), following the manufacturer’s protocol, and eluted into a final volume of 30 µL. These purified products (10 µL per reaction) were then used as templates for *in vitro* RNA synthesis using the HiScribe T7 High Yield RNA Synthesis Kit (NEB; E2040S), in a final reaction volume of 30 µL, following the manufacturer’s provided protocol. Reactions were performed at 37°C for 5–16 hours, in a thermocycler. Single-stranded DNA FISH probes were then synthesized by Reverse-Transcription using the Maxima H Minus RT enzyme (Thermo Fisher; EP0751). RT reactions (70 µL, total) contained the entire 30 µL of unpurified T7 transcription reaction, supplemented with 1 mM of each dNTP, 1x Maxima H Minus RT Buffer, 14.28 µM RT primer, 4 µL RNaseOUT (Thermo Fisher; 10777019), and 1000 U Maxima H Minus RT enzyme. Reactions were incubated at 50°C for two hours in a thermocycler, and template RNA was removed by addition of 50 µL Alkaline Lysis Buffer (500 mM NaOH; 250 mM EDTA), followed by incubation for 12 minutes at 95°C, in a thermocycler. The final DNA probe pool was purified using the Oligo Clean and Concentrator Kit (Zymo Research; D4060) following the manufacturer’s instructions, eluted into 20 µL nuclease-free water, at stored at –20°C until use.

### Polymerase Exchange Reaction (PER)

To amplify the DNA SABER-FISH signal, 5′–Bridge oligos (IDT; **Supplementary Table 1**) were extended as 10 nt concatemers (enabling them to recruit more fluorescent imager oligos), using the Polymerase Exchange Reaction (PER), as described previously^41,53^. To remove contaminating dGTP, a 40 µL reaction comprising 1x PBS; 12.5 mM MgSO_4_; 375 µM each of dTTP, dATP, and dCTP; 125 nM each Clean.G oligo and the corresponding template hairpin (**Supplementary Table 1**); and 0.5 µL Bst LF polymerase (NEB; M0275S) were pre-incubated at 37°C for 15 minutes in a thermocycler. To these, 10 µL of the corresponding bridge oligo (4 µM, final) were added, and samples were incubated at 37°C for 2 hours, followed by 20 minutes at 80°C to inactive the polymerase. PER extension was confirmed by polyacrylamide gel electrophoresis (hand-poured 10% TBE-Urea, stained with 1x SYBER Gold, Thermo Fisher S11494). Extended probes were purified using the Oligo Clean and Concentrator Kit, according to manufacturer’s instructions, and eluted in 10 uL nuclease-free water.

### DNA SABER-FISH

DNA SABER-FISH probe pools were concentrated prior to use, by combining 1.2 µM of the purified ssDNA pool and 1.2 µM of the corresponding PER-extended bridge oligo, and speed-vacuuming to dryness (Savant; DNA Speed Vac DNA120). Dried oligo pellets were stored at –20°C prior to use. Pellets were then resuspended in a buffer comprising 50% (v/v) Formamide, 2X SSC, and 0.1% Tween-20, supplemented with 0.4 µg/mL RNase-A. These Hybridization Solution samples were placed on ice while samples were processed for DNA-FISH.

Cells were briefly washed in 1 mL 1X DPBS then fixed with 1 mL fixative (4% (v/v) PFA in 1X PBS), for 10 minutes at room temperature, without agitation. Fixative was aspirated and cells were washed with 1 mL DPBS for 1 minute, and then permeabilized with 1 mL 0.5% (v/v) Triton X-100 in 1X PBS for 10 minutes, at room temperature. Cells were then washed with 1mL 1X DPBS for two-minutes, and then incubated in 1 mL of 0.1 N HCl in 1X PBS for five minutes at room temperature. Cells were then washed twice with 2X SSCT (1 mL; 1 minute per wash; room temperature) and once with 50% Formamide Wash Buffer (1 mL; 2 minutes; room temperature). Samples were then aspirated, and incubated with 1 mL pre-warmed 50% Formamide Wash Buffer, and incubated at 60°C for 20 minutes. Wash buffer was aspirated and 50 µL of hybridization solution (*see above*) was added. Samples were denatured at 78°C for 4 minutes, resting on a pre-equilibrated thermal block in a water bath, sealed, and incubated at 37°C overnight.

The next day, samples were briefly rinsed twice with pre-warmed 60°C 2x SSCT, followed by four 5-minute washes with pre-warmed 60°C 2x SSCT, and two 5-minute washes with room-temperature 2x SSCT (1 mL per wash). Samples were then rinsed with 1X PBS, aspirated, and treated with 50 µL of Fluorophore solution (1 µM each AlexaFluor 647- and Atto 488-conjugated probes, in 1X PBS, **Supplementary Table 1**). Samples were protected from light and incubated for one-hour at 37°C. Samples were then rinsed with 1 mL 1X PBST, followed by two 5-minute washes in 1X PBST, and two 5-minute 1X PBS washes (1 mL each). Samples were imaged immediately, in 1X PBS.

### FFPE Tissue Section Preparation for RNA-FISH and Immunofluorescence

Tissue sections were prepared using the FFPE Sample Preparation and Pretreatment protocol from the RNAscope 2.5 assay (ACD Bio), using glass Coplin jars. Briefly, to deparaffinize sections slides were placed in Xylenes (Sigma; 534056) for 5-minutes at RT, followed by a second 5-minute Xylene incubation, in a fresh jar. Sliders were then immediately placed in fresh 100% EtOH for 1 minute at RT with agitation, followed by another 1-minute incubation in 100% EtOH, in a fresh jar. Slides were then dried for 3 minutes on a Thermostat slide warmer (No. 26020) with its thermostat set to 4.5. Then, ∼5–8 drops of RNAscope Hydrogen Peroxide solution were added to cover the entire section, and samples were incubated for 10 min at room temperature. Slides were then washed twice for 1 minute in nuclease-free water, followed by submerging in boiling RNAscope 1X Target Retreival Reagents solution for 10-minutes. Slides were immediately placed in room temperature nuclease-free water for 1 minute, followed by another 1-minute water wash, and a 1-minute EtOH wash. Slides were dried for 3 minutes on a Thermostat slide warmer, as above. An Immedge hydrophobic barrier pen (Vector labs; H-4000) was then used to create a hydrophobic barrier around each section, and samples were dried for 3-minutes at RT. Five drops of RNAscope Protease Plus was then added to entirely cover each section. Sections were then incubated in hybridization chambers at 40°C for 40 minutes, washed twice with room temperature nuclease-free water (1 min per wash), and twice for two-minutes with 150 µL 30% Formamide Wash Buffer. 150 µL of primary RNA-FISH probes were then added. Thereafter, combined RNA-FISH and immunofluorescence were performed as described above.

### Chromatin state analysis

Epigenomic analysis of O-MAP-ChIP-enriched DNA loci was performed using ChromHMM version 1.22 (Ref. ^60^). The genomic coordinates of enriched peaks were intersected with the ChromHMM-defined states, and fold enrichments were calculated using the OverlapEnrichment function, using a chromatin state file for RUES2 cardiomyocytes^112^ (Accelerating Medicines in Partnerships; accession DSR313BZF). This function compares the observed overlap with the expected overlap based on the genomic representation of each state. This analysis determined whether the identified peaks were enriched in specific epigenomic signatures, such as active promoters, enhancers, transcribed or repressed regions. Fold enrichment of each epigenomic signature for all DNA loci was plotted as a heatmap using seaborn version 0.10.11 (Ref. ^113^). Transcription factor data were pulled from the Human Transcription Factor database version 1.011 (Ref. ^114^), and those encoded in pre-*TTN* O-MAP TIDs were subjected gene ontology analysis using Metascape^56^.

The chromatin states associated with TID subclasses (**Fig. 3**) were defined using ChromHMM annotations. The genome was partitioned into 100 kb bins, and the fraction of bases assigned to each ChromHMM state was calculated for each bin. These values were normalized to the genome-wide average for each state. ChromHMM states were grouped into active (EnhA1, EnhA2, Tx, TssA), repressive (Het, Quies, ReprPCWk, ReprPC) and bivalent (EnhBiv, TssBiv) categories. For each bin, the mean normalized enrichment of states within each category was calculated. Bins were classified as bivalent if their bivalent enrichment exceeded the genome-wide expectation (>1); remaining bins were assigned as active or repressive according to the larger normalized category enrichment. TID-encoded genes were assigned a chromatin state based on the bin containing their transcription start site (TSS). Gene expression changes were evaluated in both RUES2 (hESC) and WTC11 (hiPSC) datasets using bulk RNA-seq, based on the distribution of log₂(TPM) values within each chromatin state, as defined in WT cells. Chromatin state-specific gene ontology (GO) enrichment analysis was performed using the "goana" function in the limma R package. Top biological process terms were extracted using the topGO R package, and GO terms with fewer than three ancestors or fewer than four genes in the input list were excluded from downstream analysis.

### Bulk RNA-Sequencing and gene-expression analysis

Whole-cell RNA was isolated from hESC-CMs and hiPSC-CMs by using the QIAGEN RNEasy kit following the manufacturer’s protocol. Sequencing libraries were prepared from 500 ng input RNA per sample, using the TruSeq Stranded mRNA Prep Ligation kit (Illumina; 20020596), with IDT for Illumina TruSeq RNA UDI indexes (Illumina; 20020591), and depleting ribosomal RNA using Ribo-Zero Plus kit (Illumina; 20040525). Libraries were quantified using the Qubit dsDNA High Sensitivity assay kit (Invitrogen; Q332301), pooled, and sequenced on a DNBSEQ-T7 platform (MGI) using PE100 v3 flow cells with 100- (hESC-CM) or 150-bp (hiPSC-CM) paired-end reads.

Raw sequencing reads were demultiplexed with bcl2fastq v2.20.0 and adapters were removed using Trim-galore v0.6.10. Reads were aligned to the GRCh38.113 reference genome using STAR^115^ v2.7.11b. Genome indices were first generated with runMode = genomeGenerate (default parameters), and trimmed reads were then aligned with quantMode = geneCounts and outSAMtype = BAM. Subsequent file-format conversions were performed using standard R tools, and Bedops (v2.4.41), generating normalized coverage tracks for data visualization. Gene expression analysis was performed in R using the ReadsPerGene.out.tab output generated by STAR, removing genes with low expression (≤ 3 counts in all samples), and differential expression between conditions was quantified using the limma–voom framework^116^. Density distributions were visualized using ggplot2. Gene-set enrichment in the quadrant analysis pipeline (**Fig. 3f** and **Supplementary Fig. 5d**) was assessed by Fisher’s exact test with Benjamini–Hochberg correction, using genes that passed the corresponding RNA-seq expression filter as the background universe, and results were visualized as heatmaps. TPM normalization of filtered counts was performed using transcript lengths derived from the alignment GTF file using gtftools in -l mode^117^.

In targeted knockdown experiments (**Fig. 7** and **Supplementary Fig. 13**), whole-cell RNA was extracted using the Qiagen RNeasy Mini Kit (Qiagen; 74104), following the manufacturer’s instructions. RNA was quantified using a NanoDrop One Spectrophotometer (Thermo Fisher) and its quality confirmed on an Agilent TapeStation, using the "RNA High Sensitivity" assay (Fred Hutch Genomics core). Library synthesis, quality control, and sequencing were performed by Azenta Life Sciences/Genewiz, using their standard rRNA-depletion workflow, and sequencing on an Illumina NextSeq500 sequencing platform, operating in paired-end mode with a target depth of 100 million reads per sample. Gene expression analysis proceeded largely as described above, but was performed in R using the DESeq2 framework^118^, and defining differentially expressed genes using a threshold of |log₂ fold change| > 2.0 and adjusted p-value < 0.05. Gene-set enrichment analysis was performed using fgsea with TID-associated genes as the enrichment set. Gene-expression changes between perturbations were compared by linear regression of log₂ fold-change values.

### pre-TTN O-MAP-Seq

O-MAP labeled cells (4.8 x 10^6^ cells; four biological replicates per experimental condition) were harvested by scraping in PBSTq. Samples were then pelleted by centrifugation at 3,000 x g for ten minutes at 4°C, aspirated, flash-frozen in liquid nitrogen, and stored at –80°C until use. Samples were thawed on ice and resuspended by gentle pipetting in 1000 μL ice cold RIPA Buffer (50 mM Tris-HCl pH 7.5, 150 mM NaCl, 0.1% (w/v) SDS, 0.5% (w/v) sodium Deoxycholate, 1% (v/v) Triton X-100, 5 mM EDTA, 0.5 mM DTT), supplemented with 1x EDTA-Free Halt Protease Inhibitor Cocktail, 0.1 U/μL RNase-OUT, and 10mM sodium azide, and rocked end-over-end at 4°C for five minutes. Cells were placed in an ice-cold thermal block and then sheared using a Branson Digital Sonifier 250 outfitted with a double stepped microtip, at 10–12 Watts for 30 s intervals (0.7 s on; 1.3 s off), with 30 s resting steps between intervals, five intervals total. After every five intervals the thermal block was re-equilibrated on ice for two-minutes. Following sonication, lysates were clarified by centrifugation at 15,000 x g for ten minutes at 4°C. Supernatants were placed in fresh tubes and diluted with 1 mL Nuclear Lysis Buffer (NLB: 25 mM Tris-HCl pH 7.5, 150 mM KCl, 0.5% (v/v) NP-40, 5 mM EDTA, 0.5 mM DTT), supplemented with 1X Halt Protease Inhibitor Cocktail, 0.1 U/μL RNase-OUT and 10 mM sodium azide. From each sample 5% was removed as “input,” and to the remainder were added 100 µL of a Pierce Streptavidin Magnetic Bead slurry that had been equilibrated by two washes in 1:1 RIPA:NLB, supplemented with 10mM sodium azide, 0.1 U/μL RNase-OUT, and 1X Halt Protease Inhibitor Cocktail. Samples were incubated for 2 hours at room temperature with end-over-end agitation. Beads were collected by magnetization, and then washed with the following series of buffers (1 mL each, 5 min per wash at room-temperature with end-over-end agitation). All buffers were supplemented with 1x EDTA-Free Halt protease inhibitor cocktail, 0.1 U/μL RNase-OUT, and 0.5 mM DTT, unless otherwise noted: (1) RIPA, supplemented with 10 mM sodium azide; (2) RIPA alone (3) High Salt Buffer (1 M KCl, 50 mM Tris-HCl pH 7.5, 5 mM EDTA) (4) Urea Buffer (2 M Urea, 50mM Tris-HCl pH 7.5, 5 mM EDTA) (5) 1:1 RIPA:NLB, without protease inhibitors (6) NLB, without protease inhibitors, (7–8) two washes in TE buffer (10 mM Tris-HCl pH 7.5, 1 mM EDTA), without protease inhibitors.

RNA was isolated from both input and O-MAP-enriched samples by proteolysis in 100 μL Elution Buffer (2% (v/v) N-lauryl sarcoside, 10 mM EDTA, 5 mM DTT, 200 μg proteinase K, in 1xPBS). Reactions were shaken at 700 rpm in a Mixer HC (USA Scientific) for 1 hour at 42°C, followed by 1 hour at 60°C. RNA was then extracted once with 1 volume of phenol pH 4.3, and twice thereafter with an equal volume of absolute chloroform. Reactions were supplemented with 1µL Glycoblue and NaCl to 300 mM, and ethanol precipitated at –20°C overnight. RNA was harvested by centrifugation at 15,000 x g for 30 minutes at 4°C, washed twice with 70% ethanol, and resuspended in 40 µL nuclease-free water. To each sample, 5 µL DNA Digestion Buffer and 5 µL DNase I (Zymo Research; R1013) was added, gently mixed by inversion, and then incubated for 15-minutes at room temperature. RNA was then purified using an RNA Clean and Concentrator-5 Kit (Zymo Research; R1013), according to manufacturer’s instructions. RNA was eluted in 15 µL nuclease-free water, and its concentration was measured using a Nanodrop One (Thermo Fisher).

### pre-TTN O-MAP-Seq library preparation and sequencing

RNA samples were processed for sequencing as described previously^41^ Briefly, ribosomal RNA was first depleted by RNase-H digestion, using a pool of DNA oligonucleotides antisense to rRNA, as previously described^41,119^. 1 µg of the antisense probe pool was added to 1 µg of RNA in 200 mM NaCl, 100 mM Tris-HCl, pH 7.4, at a final volume of 10 µL. Samples were heated at 95°C for 2 minutes and then slowly cooled to 45°C at a rate of – 0.1°C/s, using a ProFlex PCR system. Then, reactions were supplemented with 10 µL of RNase H mix (10 U Hybridase Thermostable RNase H (Lucigen; H39500), 20 mM MgCl2) that was pre-heated to 45°C. Samples were incubated at 45°C for 30 minutes then placed on ice. RNA was purified by acidic phenol-chloroform extraction and ethanol precipitation. Samples were resuspended in 40 µL of nuclease-free water, and 5 µL of DNA Digestion Buffer and 1 µL of DNase I (Zymo Research; R1013) were added. Samples were mixed by gentle inversion and then incubated at room temperature for 15 minutes. RNA was purified again by the RNA Clean and Concentrator-5 Kit, eluting into 10 µL nuclease-free water. Samples were quantified on a Nanodrop One. Sequencing libraries were synthesized from 300 ng RNA using the NEBNext Ultra II Directional RNA Library Prep Kit and NEBNext Multiplex Oligos for Illumina (NEB; E7760 and E7735), according to manufacturer’s instructions. Four biological replicates were used for each condition and each library was given a unique index. Libraries were quantified using the NEBNext Library Quantification Kit for Illumina, following manufacturer’s instructions, and library quality was confirmed using an Agilent 4200 TapeStation with a “DNA High Sensitivity” kit (Fred Hutch Genomics Core). Libraries were pooled in equimolar concentrations to 20 nM aggregate concentration in nuclease-free water, with no more than 8 libraries per pool. The pool underwent 150 cycles of paired-end sequencing, on one lane of an Illumina HiSeq 4000 per pool, run in high output mode (Azenta Life Sciences).

### O-MAP-Seq Data analysis

Raw RNA-seq FASTQ files were aligned to GRCH38 using HISAT2 version 2.2.11 (Ref. ^120^), in paired-end mode with default parameters. The resulting SAM files were converted to BAM format and sorted using Samtools version 1.15.1 (Ref. ^121^). Bigwig files for visualizing strand specific information were created using deepTools version 3.5.1 (Ref. ^98^) with parameters: --filterRNAstrand forward/reverse --binSize 1 --normalizeusing BPM. Mapped reads were quantified using StringTie version 2.2.1 (Ref. ^122^) and the StringTie output was prepared for differential expression analysis using the prepDE.py function. The resulting count matrices were used for differential expression analysis using DESeq2 (Ref. ^118^) with an FDR cutoff of 0.05. For differential analysis between WT and RBM20 KO phenotypes, the DESeq2 likelihood ratio test was used to calculate (streptavidin-enriched transcripts for WT / streptavidin-enriched transcripts for RBM20 KO) / (whole-cell transcriptome for WT / whole-cell transcriptome for RBM20 KO). For computing differentially enriched transcripts between WT and RBM20 KO cells, DESeq2 was performed for WT whole-cell transcriptome compared to RBM20 KO whole-cell transcriptome using standard settings.

To identify RBP motifs that were differentially enriched between WT- and RBM20 KO-specific, factory-localized transcripts (**Supplementary Fig. 12a**), we applied Transite^123^, using the Transcript Set Motif Analysis (TSMA) framework with the human Transite motif database. The following parameters were applied: --matrix-based --Homo sapiens --svg --Transite motif database --maximum binding sites 5. Position-weight matrices were downloaded from the RBP DB database (http://rbpdb.ccbr.utoronto.ca/) and filtered to a maximal length of 20bp. RBP-recognition motifs (**Supplementary Fig. 12a**) were further filtered to limit the potential overlaps within motifs of the same set. First, for each pair of motifs the p-value of their similarity was computed using TOM-TOM^124^, and for each motif the information content of the corresponding PWM was calculated using the Shannon’s entropy: 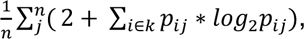 where *n* is the motif length, k = [A, T, C, G] and *p_ij_* is the probability of each base i in the specified position j. Highly similar motifs (p < 0.001) were iteratively collapsed using a greedy procedure that retained the motif with higher information content while merging the associated RBP annotations. The resulting non-redundant motif set was used for enrichment analysis with the Transite package. Enrichment was calculated as the difference in relative motif frequency between foreground and background transcript sets, and p-values were corrected using the Benjamini–Hochberg method.

For GC-content analysis, O-MAP-enriched transcripts coordinates (5′– and 3′–UTRs, introns, and exons) were extracted from the Ensembl GTF file GRCh38.113. For alternatively spliced transcript isoforms, each position was classified as exonic if it is included in at least one splicing pattern. For each transcript, the sequence of each segment was retrieved using the "getSeq" function (biomaRt package), and used to compute the average GC content, using the "letterFrequency" function (Biostrings package). P-values (comparing WT and RBM20 KO) were calculated with the "ks.test" R function (stats package).

Volcano plots were generated using EnhancedVolcano version 1.12.0 (https://github.com/kevinblighe/EnhancedVolcano). All statistical analyses (Fisher’s exact test, hypergeometric distribution test, or Student’s t-test, where appropriate) were performed in R or in python using the ggplot2 (Ref. ^125^), seaborn^113^, or matplotlib^126^ modules. Gene ontology analysis was performed using Metascape^56^, Enrichr^127^, and GSEA^128^. Gene set enrichment analyses were performed using fgsea^129^.

### Alternative splicing analysis

The rMATS package^72^ was used for alternative splicing analysis, utilizing the BAM files generated by HISAT2 with the parameters: -t paired --readLength 150. For O-MAP-Seq data (**Fig. 4a** and **Supplementary Fig. 6**), low coverage splicing events were filtered to include only transcripts with an average of 5 or more reads between replicates of each group. Differential events were further filtered for an FDR of 0.05 and an absolute change in PSI between the two groups > 10% (ΔPSI > 0.1). For additional datasets, the same rMATS framework was applied with dataset-specific junction count thresholds (≥16, ≥19, or ≥12 reads across samples for the RUES2, WTC11, and knockdown datasets, respectively). In knockdown experiments, ΔPSI values were also compared with previously reported RBM20 and QKI knockout datasets and integrated with published QKI eCLIP binding data. Sashimi plots were generated using rmats2sashimiplot (https://github.com/Xinglab/rmats2sashimiplot) using the BAM files generated by STAR coordinates and the rMATS events file for splicing event information. For the double-KO analysis, events described as attenuated were those for which the double-KO ΔPSI shifted toward WT relative to the RBM20 KO effect.

### Quantitative RT–PCR

Quantitative RT–PCR (qRT–PCR) was performed as previously described^130^. Growth medum was aspirated, and RNA was extracted with TRIZol reagent (Invitrogen; 15596026; 1 mL per well of well of a 12-well plate), according to the manufacturer’s protocol. Contaminating DNA was removed with DNase I and RNA was purified using RNA Clean and Concentrator-5 Kit as described above (*see, "pre-*TTN *O-MAP-seq"*), eluting into 30 µL nuclease-free water. RNA samples were reverse transcribed using SuperScript IV Reverse Transcriptase (Thermo Fisher; 18090010), priming with random hexamers according to the manufacturer’s protocol. cDNA was diluted with nuclease-free water, mixed with gene specific or isoform specific primers, supplemented with Rox-normalized PowerUp SYBR Green Master Mix (Thermo Fisher; A25777), and aliquoted into 384-well plates, 10 µL per well. qPCR was performed on a QuantStudio 5 Real-Time PCR System (Thermo Fisher; A34322) using four biological replicates, each measured in technical quadruplicate. Gene expression was calculated using the 2-ΔΔCt method using the housekeeping gene RPLP0. Splicing isoform gene expression was calculated using the 2-ΔCt method as previously described2; 2^-(Ct spliced isoform – Ct constitutive isoform)^131^. Primer sequences for qPCR are listed in (**Supplementary Table 5**)

### pre-TTN O-MAP-MS sample preparation

O-MAP labeled cells (2.4x10^6^ per replicate; five biological replicates per experimental condition) were harvested in PBSTq without sodium ascorbate using cell scrapers, and pelleted at 3,000 x g for 10 minutes at 4°C. PBSTq was aspirated and cells were flash-frozen in liquid nitrogen and stored at –80°C until use. Samples were thawed on ice and resuspended by gentle pipetting in 1mL CLB (20 mM Tris pH 8.0, 85 mM KCl, 0.5% (v/v) NP-40) supplemented with 1x Halt EDTA-Free protease inhibitor cocktail and 10 mM sodium azide, for 10 minutes. Lysates were clarified by centrifugation at 3,000 x g for five minutes at 4 °C followed by aspiration of supernatant. Samples were resuspended with CLB, pelleted and aspirated twice more. Pellets were then lysed by gentle pipetting in 1 mL of NLB (10 mM Tris-HCl pH 7.5, 1% (v/v) NP-40, 0.5% (w/v) sodium Deoxycholate, 0.1% (w/v) SDS) supplemented with 1x Halt EDTA-Free protease inhibitor cocktail and 10 mM sodium azide, for 10 minutes. Lysates were then placed into an ice-cold thermal block and sheared with a Branson Digital Sonifier outfitted with a double stepped microtip, at 10–12 Watts over 30 s intervals (0.7 s on; 1.3 s off), with 30 s resting steps between intervals. A total of 4 intervals were performed per sample; in between samples the thermal block was re-equilibrated on ice.

Samples were supplemented with 47.47 µL of 20% (w/v) SDS, to a final concentration of 1%, and boiled at 95°C for one hour to reverse formaldehyde crosslinks. Samples were passively cooled to 50 °C before placing on ice, and then sonicated for one more cycle, using the same setting described above. Samples were then clarified by centrifugation at 15,000 x g for 10-minutes at 4°C, and supernatants were moved to fresh tubes. To each tube was added 75 µL of Pierce Streptavidin-coated magnetic beads that had been equilibrated by two NLB washes. Samples were rotated end-over-end for 2 hours at room-temperature, and beads were collected by magnetization and aspirated. Beads were then washed with 250 µl of NLB, supplemented with 5 mM EDTA, 10 mM sodium azide, 1x Halt EDTA-free Protease Inhibitor Cocktail, and 150 mM NaCl. Beads were then moved to fresh PCR-strip tubes (Simport Scientific; T320-2N) to minimize sample loss. The following series of washes were then performed (200 µl each, 1 minute per wash, at room-temperature, with end-over-end rotation): (1–4) four washes of NLB, supplemented with 5 mM EDTA, 10mM sodium azide and protease inhibitors; (5–8) four washes of 1X PBS in nuclease-free water; (9–16) four washes of 1 M KCl, 10 mM Tris-HCl pH 7.5, 5 mM EDTA; (17–24) eight washes of 2 M Urea, 10 mM Tris-HCl pH 7.5, 5 mM EDTA; (25–28) four washes of 200 mM EPPS pH 8.5 in nuclease-free water. After the final wash the beads were moved to fresh PCR strip tubes, aspirated, and resuspended in 50 µl 200 mM EPPS pH 8.5.

Samples were transferred to a Mixer HC unit (USA Scientific) outfitted with custom "thermal couples" fashioned by wrapping a small square of aluminum foil around the end of a 1 mL pipette tip. Samples were supplemented with TCEP (Thermo Fisher; 77720) to a final concentration of 10 mM, and incubated at room temperature for 1 hour, with shaking at 700 rpm. Samples were alkylated with iodoacetamide (Sigma; I1149-5G) supplemented to a final concentration of 20 mM, and incubated in the dark for one-hour at room temperature with shaking at 700 rpm. Alkylation was quenched by the addition of dithiothreitol (Sigma; 43816) to 5 mM and incubation at room temperature for 15 minutes, with shaking at 700 rpm. To each sample was added 300 ng of mass spectrometry grade LysC (FujiFilm; 121-05063), and samples were incubated at 37°C for three hours, with shaking at 700 rpm. Samples were then supplemented with 300 ng sequencing grade modified trypsin (Promega; V5113) and incubated for exactly 16 hours at 37°C, with shaking at 700 rpm.

Beads were removed by magnetic separation, and the clarified samples were moved to fresh PCR strip tubes and frozen at –80 °C to inactivate LysC and trypsin. Samples were stored at –80°C for no longer than three days. Samples were thawed to room temperature, and to each were added 0.5 µL of 1% HPLC-grade formic acid to achieve pH < 3, as measured using pH strip paper. Samples were then desalted using C18 StageTips, largely as described previously^41^. Briefly, each StageTip was activated with 50 µL of HPLC-grade methanol, followed by addition of 50 µL of Stage Tip B (80% aq. acetonitrile; 0.1% trifluoroacetic acid) and then 50 µL of Stage Tip A (0.1% TFA). Samples were then loaded, washed with 50 µL of Stage Tip A, and stored at 4°C until use.

### O-MAP-MS data acquisition and analysis

Peptides were eluted from StageTips by addition of 50 µL 45% acetonitrile, 0.1% TFA, into a 96-well plate. Samples were dried and resuspended in 10 µL Stage Tip A. Peptides were separated on an EASY-nLC 1200 System (Thermo Fisher Scientific) using 20 cm long fused silica capillary columns (100 μm ID, laser pulled in-house with Sutter P-2000, Novato CA) packed with 3 μm 120 Å reversed phase C18 beads (Dr. Maisch, Ammerbuch, DE). The liquid chromatography (LC) gradient was 90 min long with 5–35% B at 300 nL/min. LC solvent A was 0.5% (v/v) aq. acetic acid and LC solvent B was 0.5% (v/v) acetic acid, 80% acetonitrile. MS data were collected on a Thermo Fisher Scientific Orbitrap Fusion Lumos using a data-independent acquisition (DIA) method with a 120K resolution Orbitrap MS1 scan and 12 m/z isolation window, 15K resolution Orbitrap MS2 scans for precursors from 400-1000 m/z.

Data .raw files were converted to .mzML using MSConvert 3.0.21251-d2724a5 and spectral libraries built using MSFragger-DIA^132^ (FragPipe version 19.1) with quantification through DIA-NN version 1.8.232. The database search was against the UniProt human database (updated September 17th, 2021) containing 20,420 sequences and 20,420 reverse-sequence decoys. For the MSFragger analysis, both precursor and initial fragment mass tolerances were set to 20 ppm. Spectrum deisotoping, mass calibration, and parameter optimization were enabled. Enzyme specificity was set to “stricttrypsin” and up to two missed trypsin cleavages were allowed. Oxidation of methionine, acetylation of protein N-termini, −18.0106 Da on N-terminal Glutamic acid, and −17.0265 Da on N-terminal Glutamine and Cysteine were set as variable modifications. Carbamidomethylation of Cysteine was set as a fixed modification. Maximum number of variable modifications per peptide was set to 3.

FragPipe/DIA-NN output files were processed and analyzed using the Perseus software package v1.5.6.0. Expression columns (protein MS intensities) were log2 transformed and normalized by subtracting the median log2 expression value from each expression value within each MS run. For statistical testing of significant differences in expression, a two-sample Student’s t test was applied. Gene set enrichment analysis (GSEA, **Fig. 6c**) was performed using the clusterProfiler R package, ranking proteins detected in the pulldown by the T-statistic comparing RBM20 KO and WT samples. Enrichment was evaluated for Gene Ontology biological process (GO:BP) terms with Benjamini–Hochberg correction for multiple testing. Selected enrichment plots were generated using the enrichplot package.

### SAFB knockdown

Third-generation lentiviral plasmids encoding SAFB-targeting shRNAs (**Supplementary Fig. 13a**) were designed using the VectorBuilder platform (VectorBuilder, Chicago, IL, USA), based on shRNA hairpins derived from the RNAi Consortium (TRC) database. The four top-scoring hairpins were individually cloned and screened for *SAFB* knockdown in HEK293T cells by quantitative RT-PCR, in comparison to a standard scrambled shRNA control (VectorBuilder; #Ecoli(VB010000-9549mjn)). Vectors were designed with an internal eGFP reporter, rather than a drug-selectable marker, to avoid the cardiotoxic effects of drug selection. Plasmids were synthesized by VectorBuilder and delivered as transformed *E. coli* glycerol stocks. These were expanded in LB supplemented with 100 µg/mL Ampicillin (at 32°C to avoid recombination), purified using the ZymoPURE II plasmid midiprep kit (Zymo Research; #D4201), and quantified using a NanoDrop One.

To generate lentiviral particles, 1x10^6^ HEK293T cells were seeded onto 6 cm plates in 5 mL growth medium (DMEM, supplemented with 10% (v/v) fetal bovine serum, 1x Penicillin-streptomycin, 1x Glutamax supplement, Thermo Fisher). One day thereafter, cells were transfected using the X-tremeGENE transfection reagent (Roche/Sigma; # XTG9-RO), according to the manufacturer’s instructions. Transfection mixes comprised 100 ng packaging plasmid VSV-G, 900 ng packaging plasmid psPax2, and 1 µg shRNA expression construct, in a final volume of 150 µL DMEM. Viral supernatants were collected 48 hours post-transfection, filtered through 0.45 µm syringe-mounted filters, and stored at –80 °C. Viral titers were determined by transduction of HEK293T cells and quantification of eGFP-positive cells by flow cytometry. Briefly, 1x10^5^ cells were seeded into wells of a 12-well plate and inoculated with 5–800 µL lentivirus, in the presence of polybrene. Cells were cultured at 37°C for 48–72 hours, harvested by trypsinization, and the percentage of eGFP-expressing cells was quantified (BD FACSDiscover S8 Sorter; UW SLU Flow and Imaging Core). Based on follow-up qRT-PCR analysis of *SAFB* mRNA levels, hairpin three (VectorBuilder; #Ecoli(VB250304-1487zrz)) achieved the most robust *SAFB* knockdown and was used in all subsequent experiments (**Fig. 7** and **Supplementary Fig. 13**).

For knockdown experiments, hESC-CMs were plated and cultured as described above, and transduced using a calculated multiplicity of infection (MOI) of 1.0. Following 72 hours of viral exposure, medium was replaced with fresh cardiomyocyte growth medium (RPMI, supplemented with 1x each B-27 and Penicillin-Streptomycin) and subsequently exchanged every two days. Cells were typically collected for downstream analyses (*e.g.*, imaging, RNA-seq, western blotting) 5–6 days post-infection.

### Immunoblotting

Whole-cell protein lysates were prepared from hESC-CMs in RIPA buffer (25 mM Tris•HCl, pH 7.6; 150 mM NaCl; 1% (v/v) NP-40; 1% (v/v) sodium deoxycholate; 0.1% (v/v) sodium dodecyl sulfate). Cells (∼3 × 10⁵ per well of a 6-well plate) were lysed *in situ* by addition of 1 mL RIPA buffer, followed by incubation for 5 minutes on ice with gentle swirling. Extracts were clarified by centrifugation at 14,000 × g for 10 min at 4 °C, and soluble protein concentration was measured via BCA assay (Thermo Fisher; #A55864). For each sample, 10 µg of protein was precipitated with four volumes of ice-cold acetone, incubated for 60 minutes at –20 °C, and centrifuged at 14,000 x g for 10 min. Pellets were dried using a Speed-Vac (Savant), resuspended in water, supplemented with 6× Laemmli sample buffer, and denatured at 95 °C for 5 min. Samples were resolved on 10% SDS–PAGE gels and transferred to PVDF membranes by wet transfer overnight at 4 °C (0.04 A). Membranes were blocked in 5% BSA in TBS-T (20 mM Tris, pH 7.5, 150 mM NaCl, 0.1% (v/v) Tween-20) for 1 hour and incubated with primary antibodies against SAFB (AbClonal; #A7927) at 1:1000 dilution, for 1 hour at room temperature. GAPDH (Cell Signaling Technology; 5174S) was used as a loading control. Membranes were extensively washed in TBS-T and probed with HRP-conjugated anti-rabbit secondary antibody (1:10,000 dilution) in TBS-T supplemented with 5% (w/v) BSA, for 1 hr at room temperature. After extensive washing in TBS-T, membranes were developed using the SuperSignal West Pico PLUS Chemiluminescent Kit (Thermo Fisher 34580) and imaged on an iBright chemiluminescence imager.

### Image acquisition

Fluorescence widefield microscopy was performed on a Leica DM IL, equipped with a HC Fluotar 100x oil immersion objective with a 1.32 numerical aperture and planar correction (Leica; 11506527), a white LED light source (Leica; EL6000) and a DFC365 FX digital camera (Leica; 11547004). The following filter cubes were used: Texas Red (Leica TX2 ET; 11504180; used with Dylight-550 conjugates), Cy5 (Leica Y5 ET; 11504181, used for Alexa Fluor 647), GFP (Leica GFP ET; 11504174, used for Atto-488 and AF488), and DAPI (Leica DAPI ET; 11504204). Illumination intensity was adjusted using the light source manual control; acquisition times ranged from 40–1000 ms, as controlled by the Leica LASX software. Fluorescence confocal microscopy was performed on a Leica SP8X microscope (UW Keck Imaging Center), outfitted with a HC CS2 63x oil immersion objective, with 1.40 numerical aperture with both planar and apochromatic correction. The pixel size was 0.18 μm for fixed cell distance quantification, 0.06 µm for fixed cell representative images, and 0.045 µm for FFPE distance quantification and representative images. Samples were illuminated using a 470–670 nm tunable White Light Laser system, with a typical laser power of 0.1% for DAPI, 3% for 550 nm, and 10% for 647 nm. Gain and offset settings were adjusted to avoid pixel saturation. Images were line-averaged twice, with an average pixel dwell time of 1.58 μs. A bit-depth of 16 was used and a zoom factor of 1-3 was used for all images. Structured Illumination Microscopy (SIM) images (**Fig. 7c,d**) were acquired using a ZEISS Elyra 7 super-resolution microscope equipped with a Plan-Apochromat 63×/1.4 Oil DIC M27 objective. Excitation wavelengths were set at 488 nm (power 100), 561 nm (power 100), and 405 nm (power 50). Structured Illumination Microscopy (SIM) reconstruction was performed using Zeiss ZEN Blue 3.0 software with the "weak" setting optimized for fixed cells. Imaging experiments were performed across at least three independent biological replicates, with replicate cultures thawed, processed, and imaged on separate days. Reported n values indicate the total number of individual cells analyzed across these independent experiments.

### Image processing

Images were processed using Fiji^133^ and ImageJ^48^ and multicolor overlays were made using the screen setting in Adobe Photoshop. All confocal images are maximum projections of z-stacks. Brightness and contrast were adjusted for display purposes using Fiji and ImageJ or Adobe Photoshop. In all cases, contrast adjustment was applied to improve signal visibility, by changing the minimum (black) and maximum (white) values only. Automated despeckling was applied when necessary (e.g. in DNA-FISH images with weak, diffuse speckling between cells) to reduce residual background signal. To measure the 3D distances between DNA-FISH puncta for the TTN locus and DNA-FISH puncta for an experimental TID, nuclei were first segmented in Fiji. Each nucleus was then duplicated in 3D retaining all Z-slices for DAPI and for FISH signal. Then, for each 3D composite image, the minimum edge-edge distances between *TTN* FISH loci and TID FISH loci were measured using the object-based ImageJ Distance Analysis (DiAna) plug-in^134^, using the Gaussian-filtering setting to remove noise. At least 50 cells per condition were analyzed for the distance measurements; the exact number is listed in each figure. The same pipeline was used to measure the distance between nascent *TTN* O-MAP signal and interacting RNA-FISH signal. DiAna was also used for object-based colocalization, and the shuffle function was used to calculate the statistical significance of object-based colocalization and to generate the cumulative distribution plots. Surface analysis of SIM images (**Fig. 7c,d**) was performed on cardiomyocytes from two differentiations using fixed intensity thresholds of 3.4 × 10⁴ and 2.8 × 10⁴ for SAFB and QKI signals, respectively. The number of foci per cell was compared among the 4 genotypes using a linear mixed model with the “lmer” function (lme4 R package)^135^, with genotype treated as a fixed effect and differentiation batch included as a random effect.

### Docker environment

To ensure reproducibility and transparency, all bioinformatic analyses were performed within publicly available Docker environments, each of which can be retrieved directly from Docker Hub using the image names listed below (docker pull <image_name>). Download commands and analysis workflows are available on the Bertero lab’s GitHub repository (https://github.com/sara-bianchi/TTN_O_MAP_data_analysis). Hi-C and chromatin data visualization were performed within the image hedgelab/hic_pipeline:image15. Bulk RNA-Seq processing and visualization were performed with the image hedgelab/bulk_image:image3. Splicing variant identification was performed with the hedgelab/rmats:image1 image and these results were visualized with the hedgelab/sashimiplot:image1 image.

