## Supplementary material for "Nascent transcript O-MAP reveals the molecular architecture of an RBM20-regulated single-locus subnuclear compartment organized around the *TTN* RNA": Compiled Supplementary Materials

**Supplementary Discussion**

**Supplementary Figure 1.** Genotype-specific Hi-C compartmentalization analysis and correlations with O-MAP data.

**Supplementary Figure 2.** Validation of *TTN*–CID and *TTN*–TID proximity

**Supplementary Figure 3.** *Trans*-chromosomal contact discovery with Trans-C and *Signature*

**Supplementary Figure 4.** Further characterization of TID-localized genes.

**Supplementary Figure 5.** Loss of the TTN RNA partially phenocopies the loss of RBM20.

**Supplementary Figure 6.** Further characterization of the TTN RNA factory transcriptome

**Supplementary Figure 7.** Validation of *TTN* factory resident transcripts identified by O-MAP-Seq

**Supplementary Figure 8.** Loss of RBM20 induces pervasive alternative splicing events beyond its direct target transcripts.

**Supplementary Figure 9.** Validation of the RBM20 KO–Δ*TTN* Pro Double Knockout line.

**Supplementary Figure 10.** Loss of the *TTN* RNA parallels RBM20-dependent splicing changes more closely than does loss of the Titin protein.

**Supplementary Figure 11.** RBM20 KO-induced splicing events attenuated after loss of the *TTN* RNA

**Supplementary Figure 12.** Concordance of RNA-binding proteins and their cognate motifs in the *TTN* RNA factory proteome and transcriptome.

**Supplementary Figure 13.** Validation of SAFB knockdown.

### SUPPLEMENTARY DISCUSSION

#### *Regarding O-MAP's labeling radius and its capture of compartmental interactions*

It may seem counterintuitive that O-MAP—like all tyramide/peroxidase-based proximity-labeling approaches—nominally labels molecules within ~50 nm of its molecular target, and yet the novel compartmental factors the technique has identified (*e.g.* CIDs, TIDs, factory transcripts) appear to localize at markedly higher distances. We attribute this apparent discrepancy to two principal factors.

First, the effective labeling radius should be regarded as an approximate operating range rather than a fixed physical constant. Indeed, reported diffusion distances remain surprisingly variable, ranging from ~5 nm[1] to several hundred nanometers[2], and are likely influenced by experiment-specific parameters like solvent conditions and local molecular crowding within the target subcellular microenvironment[3]. Our quoted ~50 nm value is based on what we believe to be the most rigorous experimental evidence to date. This includes work from the Macmillan group, who used an *in vitro* tyramide labeling assay, combined with super-resolution imaging, to measure radical diffusion in an uncrowded aqueous solvent, yielding an average radius of 50–75 nm[4]. Furthermore, recent work from the Zhou and Chen labs applied a clever *in vitro* assay that uses tethered substrates on a DNA origami array to measure the tyramine labeling radius via a sequencing-based readout[5]. This yielded an average diffusion radius of ~11 nm, and a maximum labeling envelope of ~55 nm—similar to the Macmillan measurement. These numbers seem reasonable, given our own qualitative imaging results, since we often observe a "cloud" of O-MAP biotinylation that is slightly larger than the target RNA's FISH signal[6] (**Supplementary Discussion Fig. 1**). Because this size difference is larger than the diffraction limit of confocal imaging, it is likely that, under our preferred labeling conditions, diffusion may be modestly larger than the quoted Macmillan and Zhou/Chen measurements.

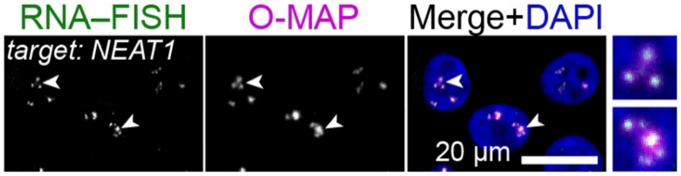


**Supplementary Discussion Figure 1. Qualitative assessment of biotin radical diffusion during O-MAP labeling.** Combined O-MAP and RNA-FISH targeting the noncoding RNA *NEAT1*, in HeLa cells. Note that biotinylation puncta are slightly larger and more diffuse than the FISH signal (examples noted by arrowheads and shown at ~3x zoom on the right). Data adapted from Tsue, Kania *et al* (2024), *Nature Methods*[6].

A second important consideration is the organizational nature of the *TTN* factory itself, which does not seem to form a rigid, precisely defined structure, but instead may behave like a liquid-like biomolecular condensate that varies dynamically across cells[7]. This means that, while CIDs, TIDs, and compartmental transcripts may exhibit a median separation of ~1 µm from nascent *TTN*, in WT CMs there is a significant population of cells in which these molecules are located close enough to be captured by O-MAP. As factors become more distal, the probability that these molecules localize within this capture radius decreases (**Supplementary Discussion Fig. 2a**). Although our imaging experiments (*e.g.,* **Supplementary Figs. 2, 7**) suggest that only a small fraction of cells fall within this labeling boundary, our omics data strongly suggest that this signal is sufficient to reveal differential enrichment (**Supplementary Discussion Fig. 2b**). Thus, O-MAP does not require that a compartmental factor reside within the biotinylation radius in every cell. Rather, factors that are preferentially enriched near the target RNA have a higher probability of entering this capture radius within some proportion of the cell population, producing measurable enrichment even when their average separation is large.

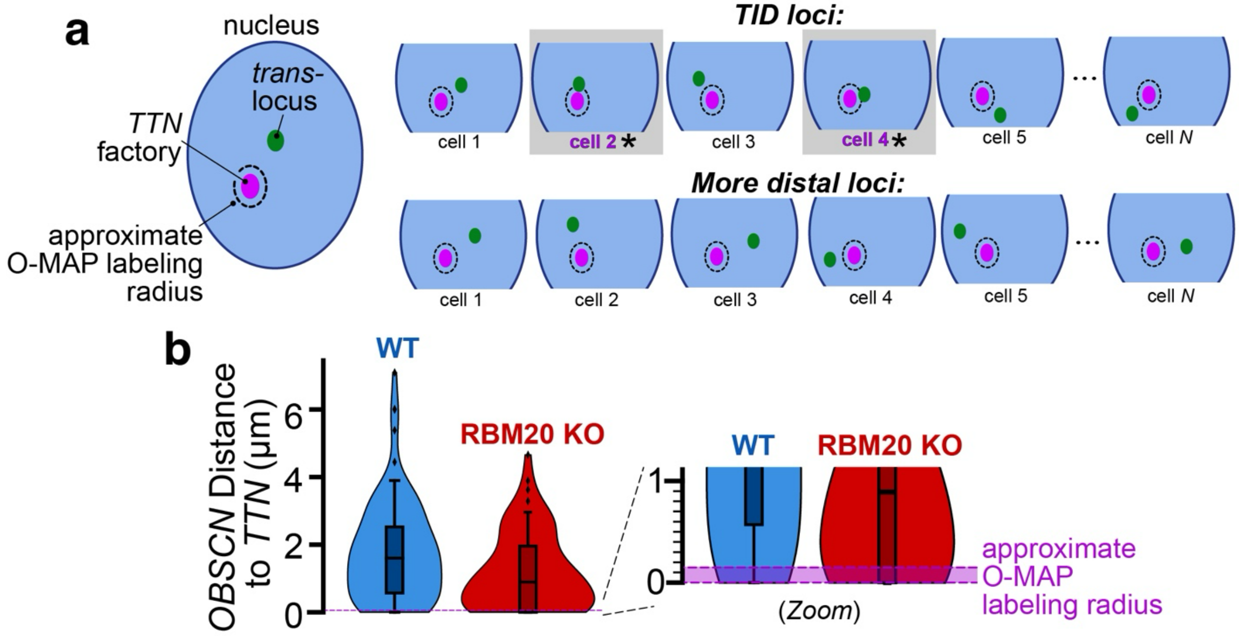


**Supplementary Discussion Figure 2. Model for O-MAP capture of micron-scale compartmental interactions. a**, Factors that are enriched near the *TTN* RNA factory have a higher probability of localizing within the O-MAP biotinylation radius than do more distal factors. The illustrated example depicts a TID locus (*top*) vs. more distal loci (*bottom*), but the same principal also applies to factory-enriched transcripts and proteins. **b,** We hypothesize that O-MAP captures factory-resident components from only a small proportion of cells. Data taken from (**Supplementary Fig. 7a**).

We note that our imaging-based analyses indicate an increase in the average spatial distance between *TTN* and TID loci in individual nuclei that is not mirrored by a comparable loss of population-averaged Hi-C contacts (**Supplementary Fig. 2**). We do not view these observations as contradictory, because DNA-FISH and Hi-C interrogate distinct aspects of nuclear organization: FISH measures physical separation between loci in single cells, whereas Hi-C reports ensemble-averaged contact frequencies and is most sensitive to relatively close-range ligation events. Thus, an increase in the typical spatial separation of TTN and TID loci can coexist with preservation of a subset of close contacts detectable by Hi-C. Taken together, these orthogonal measurements therefore support the more conservative interpretation that RBM20 loss weakens or relaxes, rather than completely abolishes, the underlying TTN-centered chromatin architecture.

#### *How are proteins and RNAs recruited to the TTN RNA factory?*

While our studies have established a rich molecular characterization of the *TTN* factory, questions remain as to how cardiomyocytes assemble this complex interactome. For example, it may seem counterintuitive that the factory—which in WT cells is defined by prominent puncta of RBM20[8]—is relatively devoid of RBM20 target transcripts other than *TTN* (**Supplementary Fig. 6**). However, we do not view this as inconsistent with RBM20's proposed architectural role. Rather, based on our data and prior biophysical studies[7], we propose that the *TTN* factory adopts an RBM20-dependent core-shell structure, similar to the nested compartmental architecture observed in several other RNA-centric nuclear compartments[9]. In our model, RBM20 helps establish the factory's core through multivalent interactions nucleated along the nascent *TTN* transcript. This core then recruits a larger cohort of outer-shell proteins, which in turn mediate interactions with other transcripts and chromatin. In this regard, RBM20 would serve dual roles as both a key regulator of *TTN* biogenesis and an architectural hub that helps recruit additional factors to pattern compartmental composition.

This proposed assembly mechanism closely parallels those of other RNA-centric nuclear compartments, including Barr bodies and paraspeckles[10, 11]. These compartments are built around scaffolds of noncoding RNA (*XIST* and *NEAT1,* respectively) which organize assemblies of RNA-binding proteins at significantly higher multiplicity[12, 13]. To drive this super-stoichiometric recruitment, each nucleating RNA directly binds to a cohort of "inner-shell" RBPs, which in turn recruit higher-order assemblies of additional proteins that collectively comprise the compartment's "outer shell" structure[9]. This secondary recruitment often exploits biomolecular condensation processes like liquid-liquid phase separation (LLPS), as mediated by intrinsically disordered domains (IDRs) within compartmental proteins[14]. Importantly, RBM20 exhibits many of the hallmarks expected of such an inner-shell architectural factor: it binds nascent *TTN* RNA at high multiplicity (the unspliced *TTN* transcript harbors more than 400 eCLIP-validated RBM20 binding sites[7]), it contains multiple prominent IDRs[15], and it is capable of undergoing biomolecular condensation[7]. Our data have further demonstrated that such RBM20 assembles are required to recruit other compartmental factors like SAFB and QKI (**Fig. 7a–b**), and that loss of these RNP assemblies globally remodels *TTN's* interactions with RNA and chromatin. Collectively, these data support RBM20's role as a key inner shell factor, while suggesting that the RBM20-dependent proteome identified by O-MAP (**Fig. 6**) comprises much of the factory's outer shell.

How, then, is the factory's transcriptome established? In general, the pathways that govern RNA subnuclear compartmentalization are less well characterized than the analogous protein-centric mechanisms described above, and so our model remains largely speculative. Evidence suggests that RNA recruitment can be in part mediated through direct RNA–RNA interactions (RRIs)[16], which are themselves capable of forming multivalent assemblies that undergo LLPS[17, 18]. However, due to the size and complexity of the *TTN* factory transcriptome (**Fig. 4** and **Supplementary Table 4**) and the lack of robust informatic tools for identifying such RRIs, we have not attempted to assess how this mechanism contributes to transcript recruitment within the factory. Alternatively, a non-mutually exclusive mechanism posits that RNAs are recruited to the factory indirectly, as clients of factory-resident RNA-binding proteins. This model is perhaps best exemplified by nuclear speckles, which have been proposed to act as splicing "hubs" where pre-mRNAs accumulate through interactions between pre-mRNA intronic sequences and speckle-resident RBPs[12, 19, 20]. The apparent concordance of our O-MAP-Seq and O-MAP-MS data suggests that a similar model may contribute to the *TTN* factory's transcriptome (**Supplementary Fig. 12**). Our double KO experiments (**Fig. 5f–h**) also lend support to this model, suggesting that, in the absence of RBM20, remodeling of the *TTN*-proximal proteome in turn reshapes the factory transcriptome (**Fig. 4e**), with downstream consequences for RNA processing (**Fig. 5**). While we have not yet characterized these putative RBP-RNA interactions experimentally, we hope future studies will help clarify the mechanisms that organize the factory transcriptome, and how remodeling of this process contributes to RBM20-associated cardiomyopathy.

### SUPPLEMENTARY FIGURES

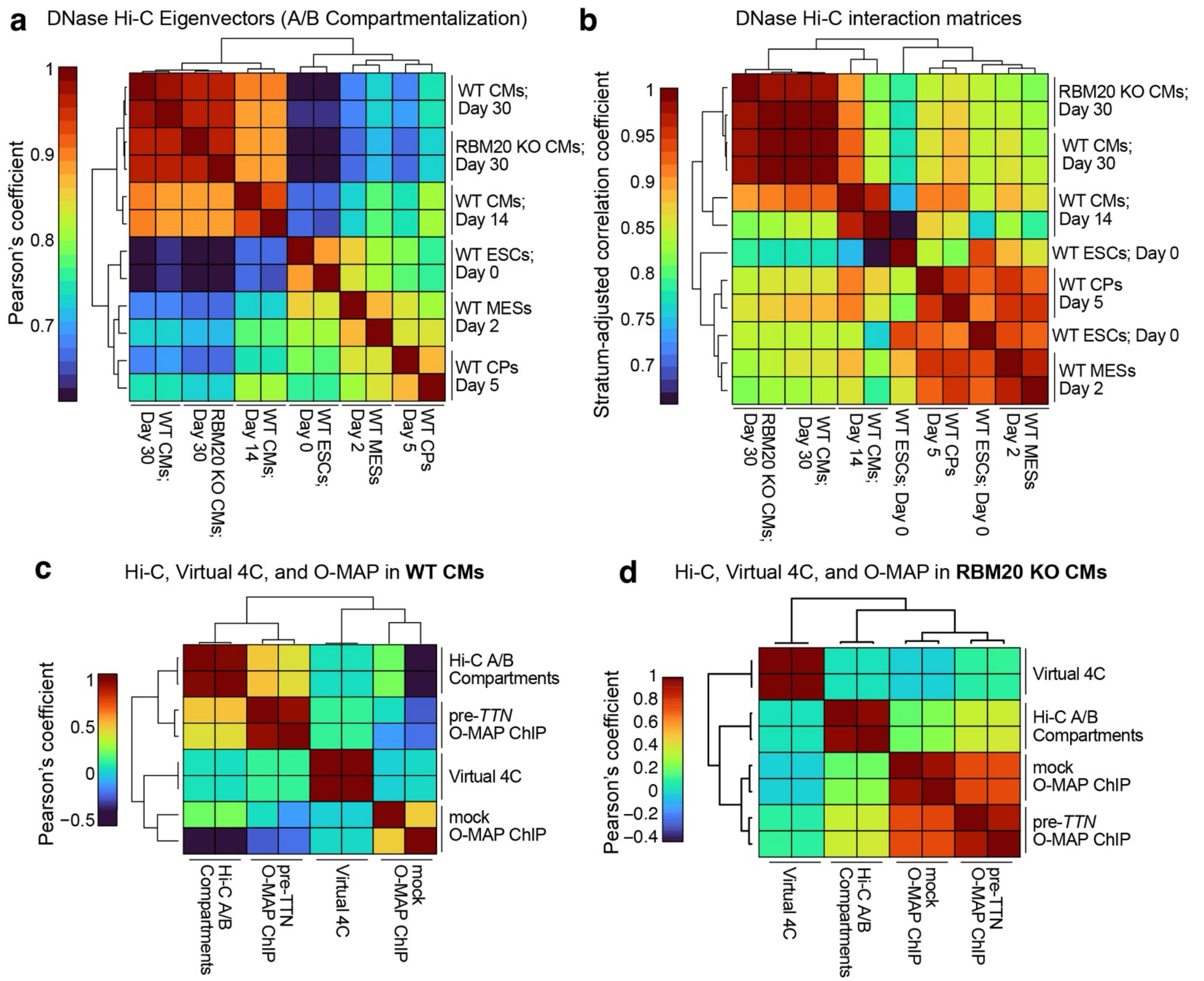


**Supplementary Figure 1. Genotype-specific Hi-C compartmentalization analysis and correlations with O-MAP data.**

**a–b,** Correlations between DNase Hi-C datasets. Newly obtained Hi-C datasets from day-30 WT and RBM20 KO embryonic stem cell derived cardiomyocytes (hESC-CMs) were compared to existing DNase Hi-C data from an *in vitro* differentiation course, comprising ESCs (Day 0), mesoderm (MESs; Day 2), cardiac progenitors (CPs; day 5), and early cardiomyocytes (CMs; Days 14). n = 2 for all conditions; individual replicates shown. **a,** Pearson correlation coefficients of genome-wide DNAse Hi-C eigenvectors. **b,** Stratum-adjusted correlation coefficients of the same comparisons. Note high correlations between WT and RBM20 KO CMs, indicating broadly stable chromatin architecture. **c–d,** Correlations between DNase Hi-C compartmentalization, Virtual 4C anchored on the *TTN* locus, pre-*TTN* O-MAP-ChIP, and mock O-MAP-ChIP. O-MAP data were processed as $\log_2$(fold change, Pulldown/Input); Virtual 4C data are the *TTN* promoter interaction profiles with bins on chromosome 2, normalized over the total number of *TTN cis*-contacts. n =2 for all conditions; individual replicates shown. **c,** WT CMs. Note the strong correlation between Hi-C A/B compartments and, to a lesser extent, Virtual 4C, with pre-*TTN* O-MAP-ChIP. **d,** RBM20 KO CMs. Note the loss of correlation between pre-*TTN* O-MAP-ChIP and Hi-C/Virtual 4C data, and the increased correlation between pre-*TTN* and mock O-MAP ChIP, indicating a nearly complete loss in pre-*TTN*–chromatin interactions.

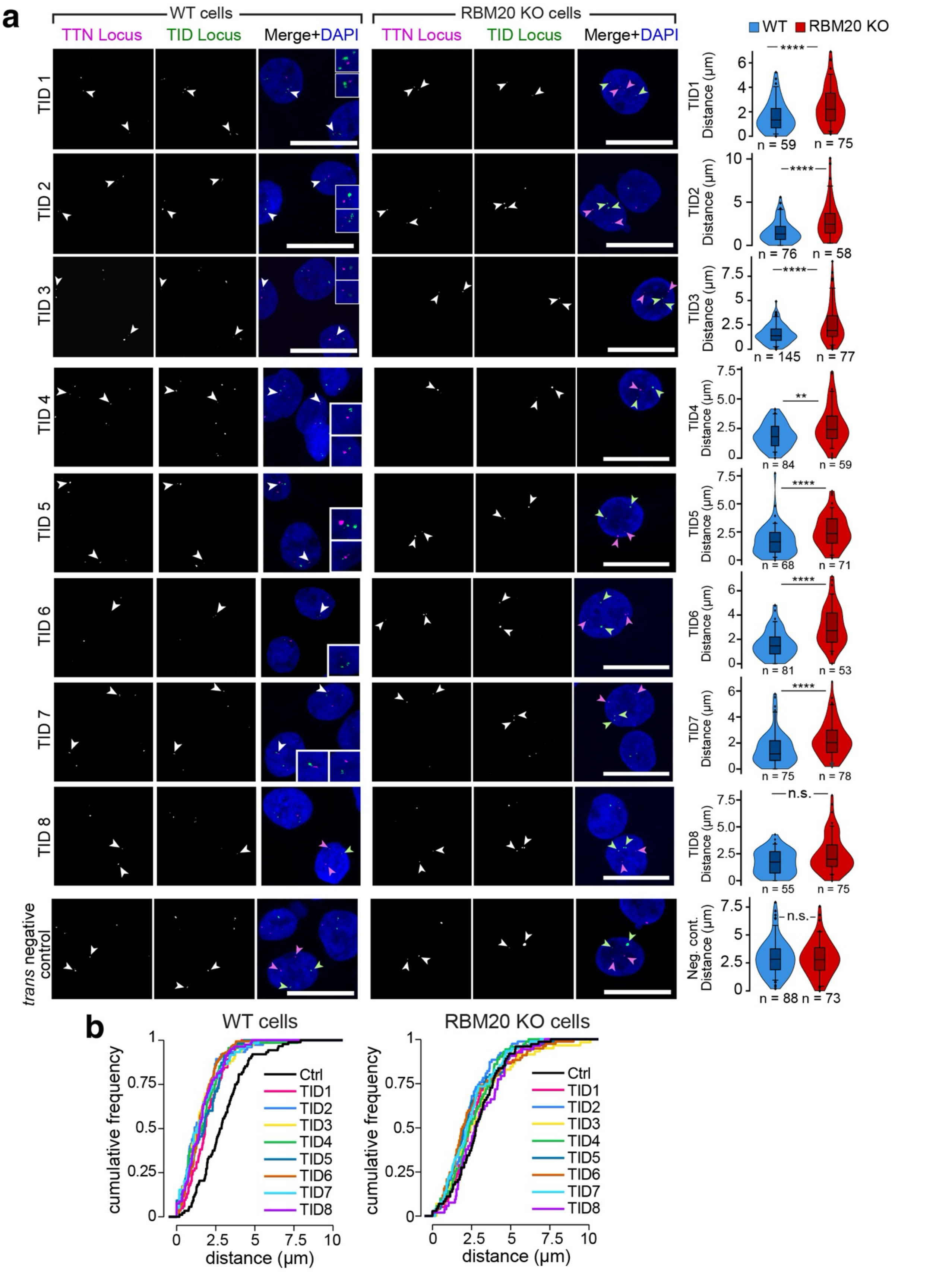


**Supplementary Figure 2. Validation of *TTN*–TID proximity. a,** Two-color DNA SABER-FISH for eight TID loci and one negative control (*bottom*). *Left:* WT cells. Proximal pairs of loci are indicated by arrowheads and highlighted in insets (TIDs1–3: ~3x zoom; TIDs 4–7: ~2.5x zoom). *Middle:* RBM20 KO cells. *Right:* Quantified minimal 3D distances between the *TTN* locus and other loci. For each condition, the number of quantified cells is listed below the corresponding graph. Violin plots represent aggregated data and indicate the number of cells, pooled from ≥ 3 independent biological replicates. Internal box and whisker plots indicate median, 25th, and 75th percentile and the 5th–95th percentile range. *P*-values: Kruskal-Wallis test followed by Dunn's multiple comparisons, vs WT. n.s.: > 0.05; ** < 0.01; *** < 0.005; **** < 0.0001. Scale bars: 20 µm. **b,** Cumulative distribution functions (CDFs) of the minimal 3D distances between the *TTN* locus and TID loci, in WT (*left*) and RBM20 KO (*right*) cardiomyocytes.

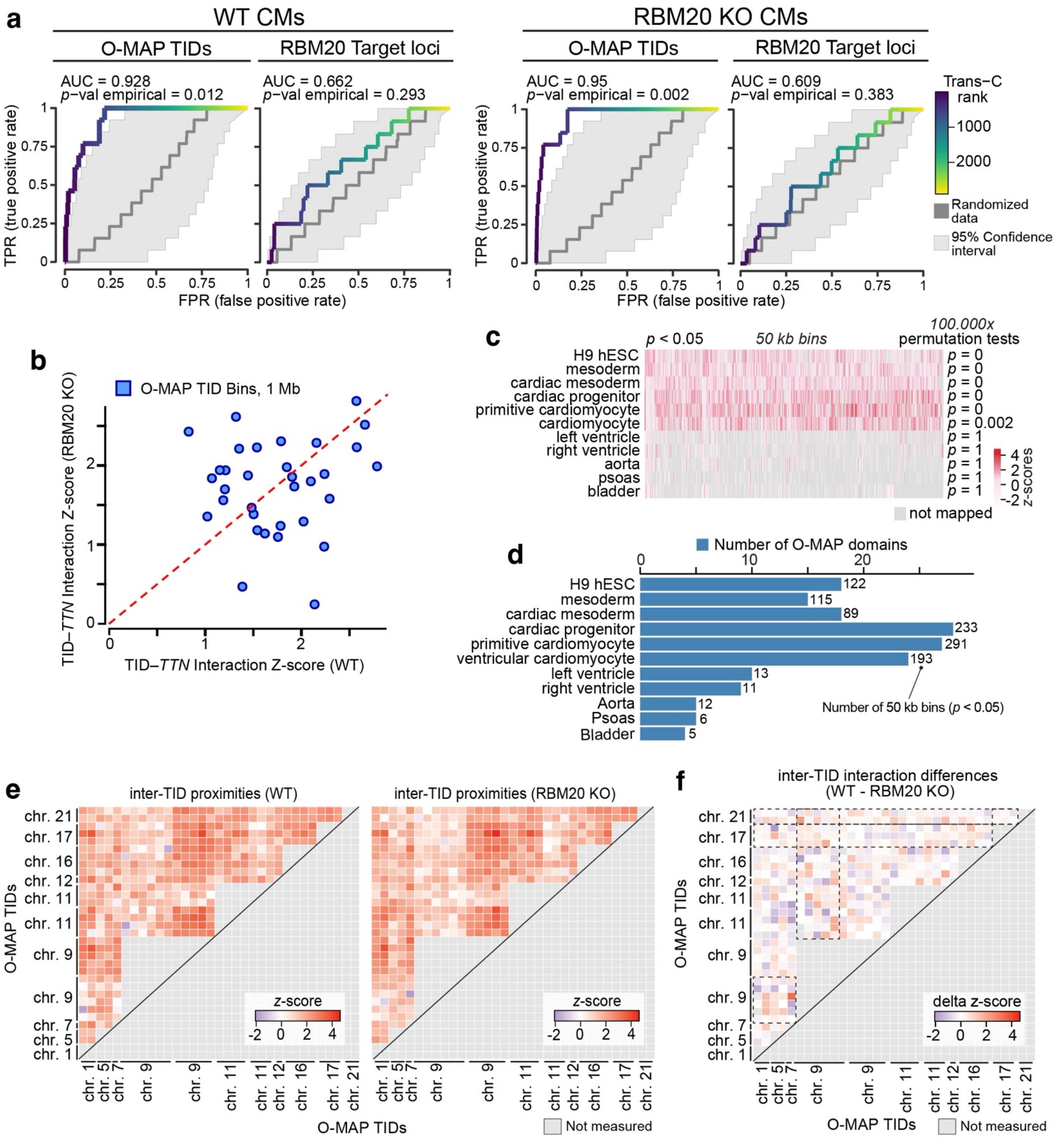


**Supplementary Figure 3. *Trans*-chromosomal contact discovery with Trans-C and Signature**
**a,** Evaluating classes of *trans*-chromosomal interactions using the Trans-C pipeline[21]. Receiver Operating Characteristic (ROC) analysis of O-MAP-TIDs and RBM20-target loci in WT CMs (*left*) and RBM20 KO CMs (*right*). Trans-C ranks were derived from a random walk, with restart on a DNase Hi-C-derived interchromosomal network, using the *TTN* locus as seed. In each analysis, true positives (TPs) were defined as loci within the indicated list of putative *trans*-interactors; false positives (FPs) were all other loci. Empirical *p*-values were calculated by comparing the observed Area Under the Curve (AUC) values to those obtained from "randomized data"—1000 random walks starting from other genomic seeds (dark gray bar = 50th percentile of random walks; light-gray shade, 5%–95% confidence interval). **b,** Signature[22] analysis of the interaction strengths (measured

as Z-scores) of *trans*-chromosomal proximities between the *TTN* locus and each TID bin. Scatterplot visualizes changes in these scores between WT and RBM20 KO CMs. Note that scores remain positive for all bins in both conditions, with no significant global shift in the population. **c,** *Signature* analysis of publicly available Hi-C datasets across multiple cell types reveals the developmental trajectory of TTN–TID *cis*-chromosomal proximities (bin size: 50 kb, chromosome 2). Empirical *p*-values shown at right indicate enrichment of TTN–TID *cis*-chromosomal interaction strength relative to the genome-wide *cis*-chromosomal proximities. **d,** Number of significant *TTN*-TID chromosomal proximities shown in (**c**). **e,** *Z*-scores of inter-TID *trans*-chromosomal proximities in WT and RBM20 KO, as measured by *Signature*. **f,** Changes in *z*-scores (WT - RBM20 KO) are shown for all *inter*-TID proximities. Dashed boxes indicate significant changes of TID–TID proximities (Mann–Whitney tests: chr. 9 $p = 0.01147024$; chr. 17 $p = 0.03174427$; chr. 21 $p = 0.01506076$).

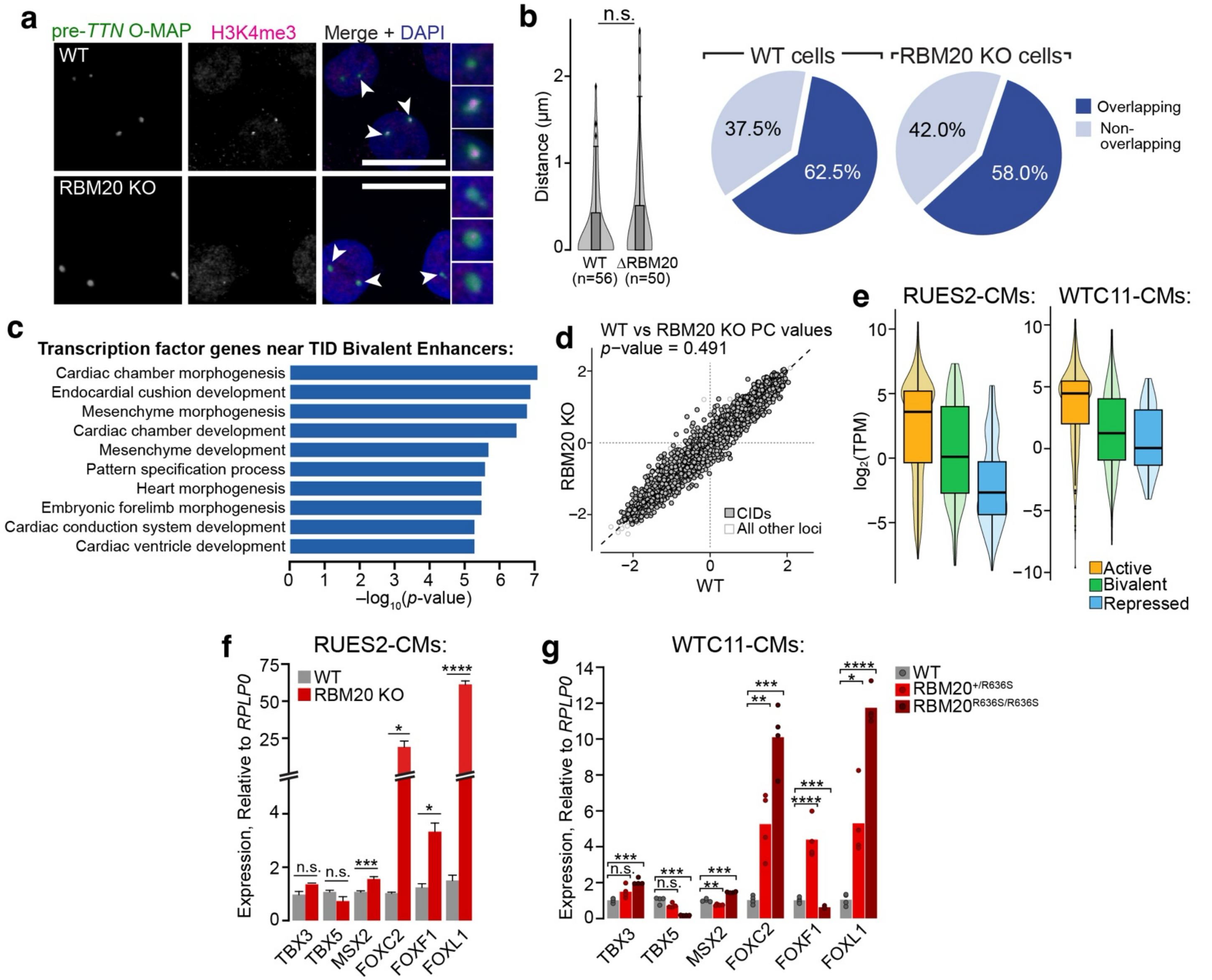


**Supplementary Figure 4. Further characterization of TID-localized genes. a,** Nascent *TTN* frequently colocalizes with prominent puncta of Histone 3 Lysine 4 trimethylation (H3K4me3), an activating epigenetic mark. *TTN* visualized by O-MAP; H3K4me3 by immunofluorescence. Scale bars, 20 μm. Insets at ~3x zoom. **b,** Nascent *TTN* and H3K4me3 puncta (when present) were essentially superposable within the detection limit of confocal microscopy (*left*; depicting minimal distances between H3K4me3 and pre-*TTN*), and observed in nearly two-thirds of nuclei, irrespective of RBM20 status (*right*). For each condition, the number of quantified cells is listed below the corresponding graph, as pooled from ≥ 3 independent biological replicates. **c,** TIDs contain transcription factor genes that appear regulated by bivalent enhancers in an RBM20-dependent manner. Gene Ontology (GO) analysis for transcription factor genes near TID-localized bivalent enhancers. The top ten enriched terms are shown. **d,** Scatterplot of DNase Hi-C eigenvectors for WT and RBM20 KO CMs. Bins belonging to the CIDs are highlighted. No significant shift in the distribution can be observed between conditions (two-sided Wilcoxon signed-ranked test). **e,** Gene expression distribution for genes belonging to each epigenetic state described in (**Fig. 3e**). *Left:* WT RUES2 CMs, and *Right:* WT WTC11 CMs. Gene expression is plotted as the logarithm of TPM (Transcripts Per Million). Boxplots indicate the median, 25th, and 75th percentiles. **f–g,** Forkhead Transcription Factors are upregulated upon loss of RBM20. **f,** Quantitative RT-PCR (qRT-PCR) for TID-encoded transcription factors in WT (gray) and RBM20 KO (red) RUES2-derived cardiomyocytes (RUES2-CMs). **g,** Similar changes in gene expression were observed in WTC11-derived CMs bearing the disease-associated RBM20 point mutation R636S. Data for WT, heterozygous ($RBM20^{+/R636S}$), and homozygous ($RBM20^{R636S/R636S}$) lines are shown. Note dose-dependent increase in *FOXC2* and *FOXL1* expression. In all qRT-PCR experiments, expression was quantified relative to *RPLP0* (four biological replicates each measured in technical quadruplicate). *p*-values: 2-tailed Student's t-test. n.s.: > 0.05; ** < 0.01; *** < 0.005; **** < 0.0001.

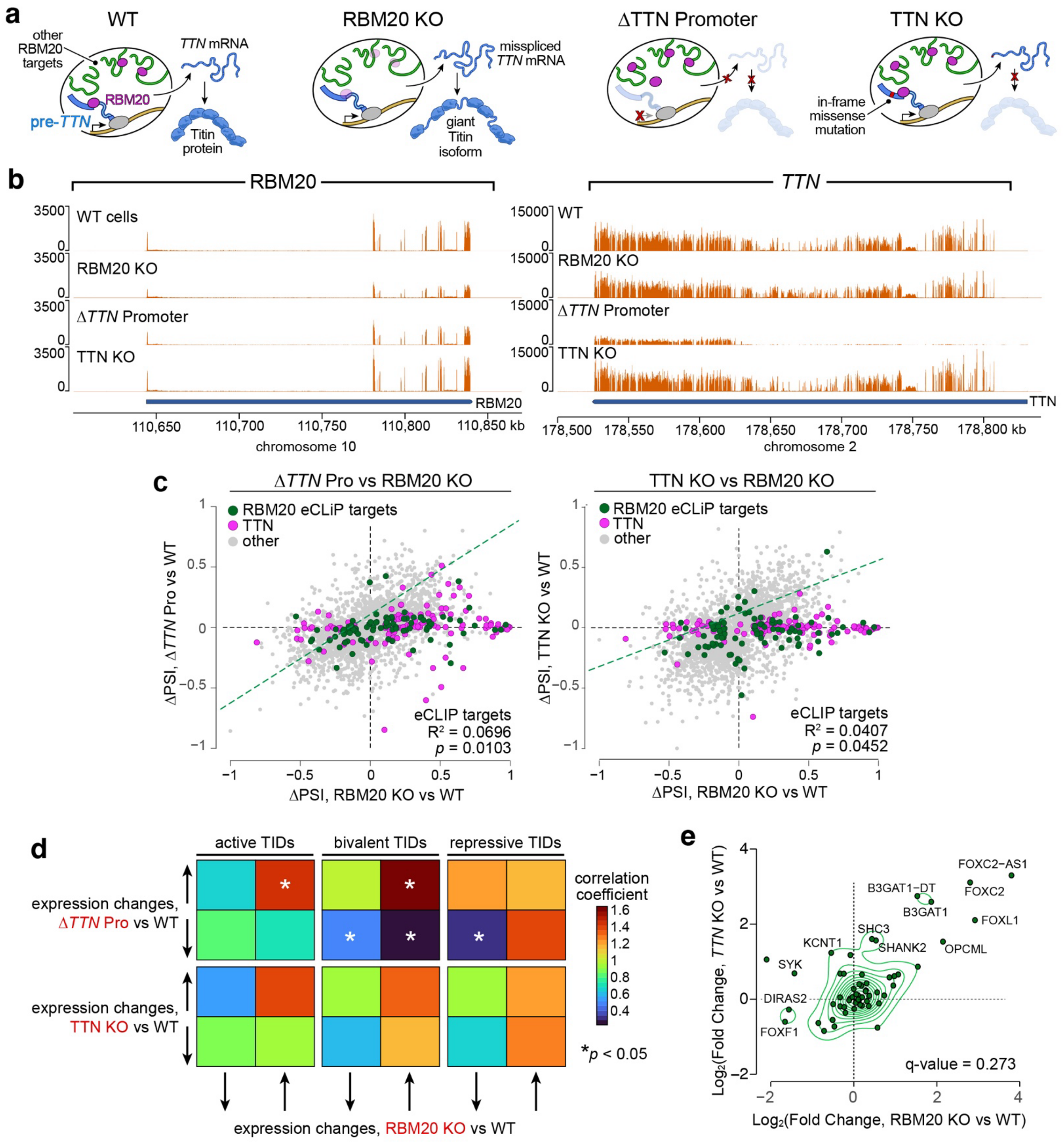


**Supplementary Figure 5. Loss of the *TTN* RNA partially phenocopies the loss of RBM20.**
**a,** Schematic of the model. Homozygous loss of RBM20 (frameshift mutation, "RBM20 KO," *left*), loss of the *TTN* transcript (removed by deleting the *TTN* promoter, "Δ*TTN* Pro," *middle*), or of the Titin protein (frameshift mutation, "TTN KO," *right*) will impair sarcomere formation. If the *TTN* transcript plays a structural role in the *TTN* RNA factory, then the gene expression changes induced by its loss—and not the absence of the Titin protein—should phenocopy those induced by RBM20 knockout. **b,** Cell line validation. RNA-seq reads at *RBM20* and *TTN* loci, binned at 100 bp. Loss of *TTN* does not influence RBM20 expression levels, or *vice versa*. Note that RBM20 frameshift mutation leads to only partial nonsense-mediated decay, resulting in detectable RBM20 mRNA that is, however, not translated[8]. Also note that, in ΔTTN Pro cells, the internal promoter for the shorter Cronos isoform produced the expected shorter transcripts originating near the 3´ end of the gene[23]. These transcripts lack the RBM20 binding site-rich 5´ region, as noted previously[7, 8]. **c,** Scatterplots of global ΔPSI

distributions in (*left*) Δ*TTN* Pro and (*right*) TTN KO cells (hESC-derived cardiomyocytes) relative to RBM20 KO cells. Shown are splicing events significantly changing in at least one of three perturbations (FDR < 0.5 and |ΔPSI| > 0.10). Dashed lines indicate linear fits of the splicing events in RBM20 target transcripts, excluding *TTN*. Note higher correlation between Δ*TTN* Pro and RBM20 KO cells (*left*)**. d,** Enrichment analysis for genes in each state, comparing classes of transcript that are up- or down-regulated in Δ*TTN* Promoter (*top*) and TTN KO (*bottom*) cells, to those that are likewise up- and down-regulated in *RBM20 KO* cells. Genes were classified as up or downregulated upon each perturbation based on the sign of the $\log_2$(fold-change), relative to WT. *p*-values were evaluated with a two-sided Fisher's exact test and adjusted with Benjamini-Hochberg's multiple comparisons correction (* if $p_{adj} < 0.05$). Note that the strongest observed enrichment was in bivalent TID-encoded genes that were up-regulated in both Δ*TTN* Pro and RBM20 KO cells (**Fig. 3f**), while changes in gene expression for the same genes in TTN KO cells were insignificant. **e,** Gene expression changes upon perturbation of the RBM20 splicing factory and the TTN KO control. The enrichment of genes belonging to the upper right corner (concordantly upregulated—positive log2FC for both the RBM20 KO and TTN KO) was evaluated as the ratio between the percentage of bivalent and total double upregulated. The q-value was calculated with a two-sided Fisher's exact test.

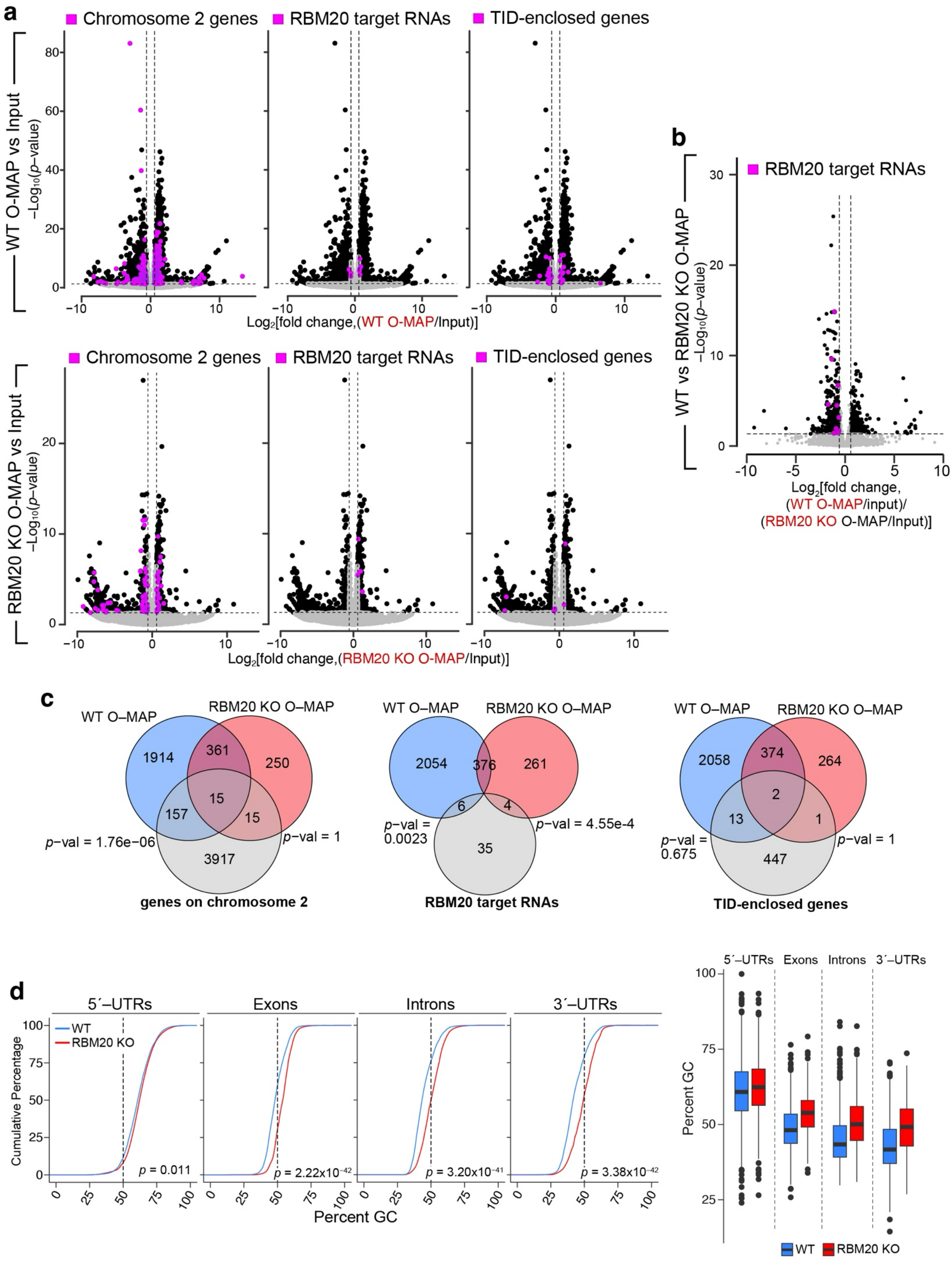
a
Chromosome 2 genes
RBM20 target RNAs
TID-enclosed genes
WT O-MAP vs Input
$-Log_{10}(p-value)$
$Log_2$[fold change,(WT O-MAP/Input)]
RBM20 KO O-MAP vs Input
$Log_2$[fold change,(RBM20 KO O-MAP/Input)]
b
RBM20 target RNAs
WT vs RBM20 KO O-MAP
$Log_2$[fold change,
(WT O-MAP/input)/
(RBM20 KO O-MAP/Input)]
c
WT O–MAP
RBM20 KO O–MAP
1914
361
250
15
157
15
3917
p-val = 1.76e−06
p-val = 1
genes on chromosome 2
2054
376
261
6
4
35
p-val = 0.0023
p-val = 4.55e-4
RBM20 target RNAs
2058
374
264
2
13
1
447
p-val = 0.675
p-val = 1
TID-enclosed genes
d
5´–UTRs
Exons
Introns
3´–UTRs
WT
RBM20 KO
Cumulative Percentage
p = 0.011
$p = 2.22x10^{-42}$
$p = 3.20x10^{-41}$
$p = 3.38x10^{-42}$
Percent GC

**Supplementary Figure 6** (*previous page*)**. Further characterization of the *TTN* RNA factory transcriptome.** **a,** pre-*TTN*-proximal transcripts exhibit modest, RBM20-dependent enrichment for other genes on chromosome 2 (*left*), but not RBM20 eCLIP targets (middle) or TID-enclosed genes (*right*), each highlighted in magenta. Volcano plots of the pre-*TTN* O-MAP-seq data in (*top*) WT CMs and (*bottom*) RBM20 KO CMs. Significant genes: $p_{adj.}$ <0.05 and |fold change| > 1.5. **b**, Few RBM20 target transcripts are enriched in the differential *TTN*-proximal transcriptome. **c,** Venn diagrams summarizing the data in (**a**). *p*-values: one-sided Fisher's exact test. Note the significant, though modest, selective enrichment of transcripts arising from chromosome 2-encoded genes in WT cells. **d,** RBM20 KO-specific pre-*TTN*-proximal RNAs exhibit higher GC content than WT-specific transcripts, distributed across most transcript features. *Left*: cumulative distribution plots. *Right*: Box-and-whisker plots. Note the markedly higher GC content among RBM20 KO-specific transcript exons, introns, and 3´–UTRs. *p*-values, Kolmogorov–Smirnov test.

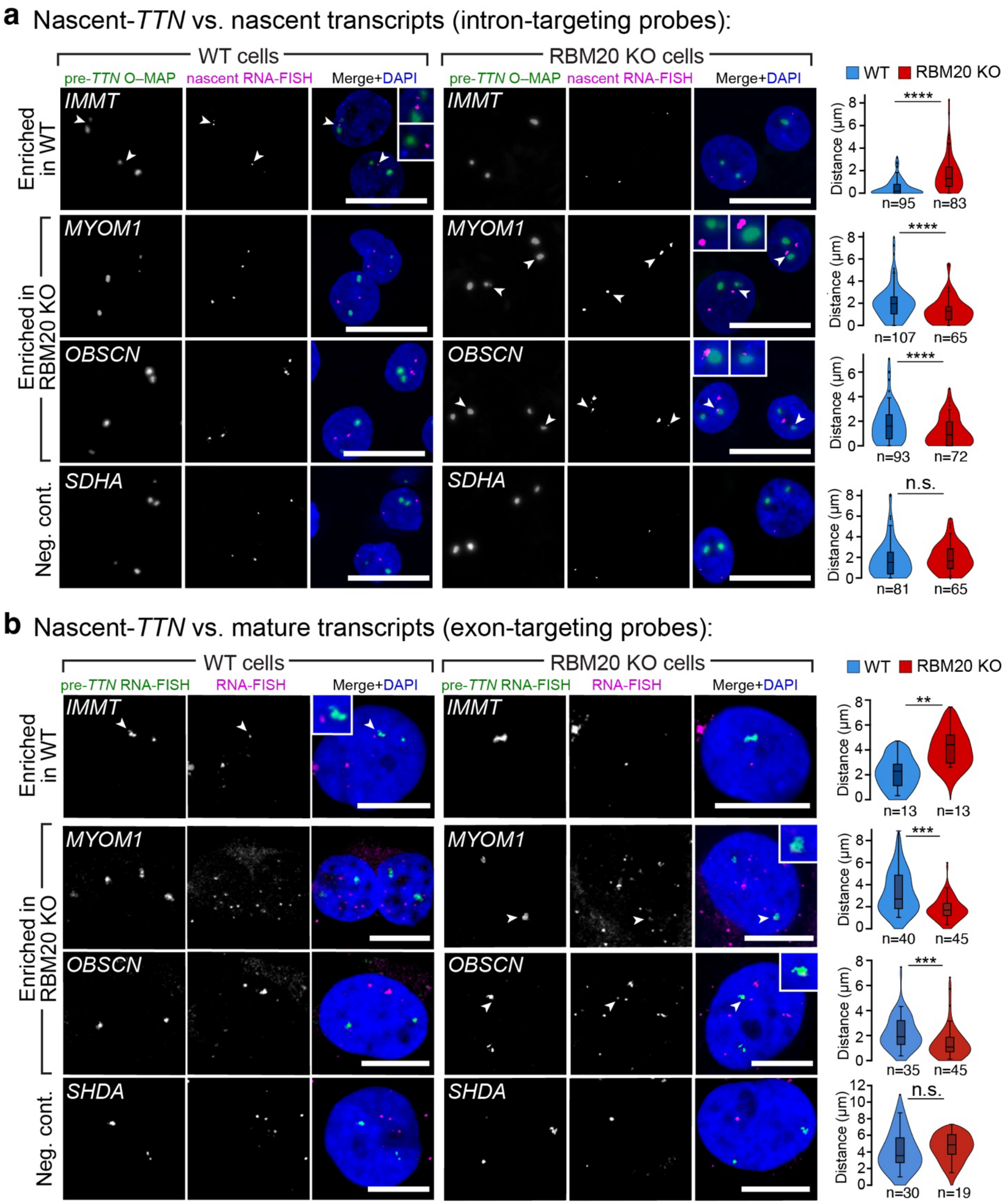


**Supplementary Figure 7. Validation of *TTN* factory resident transcripts identified by O-MAP-Seq.** Colocalization studies of pre-*TTN* and novel interacting transcripts. *IMMT* is a WT-specific pre-*TTN* interactor; *MYOM1* and *OBSCN* are RBM20 KO-specific; *SDHA* is a negative control. O-MAP Imaging[6] was used to visualize pre-*TTN* (*green*), and RNA-FISH to visualize candidate RNAs (*magenta*). **a**, Proximity of pre-*TTN* with nascent transcripts, imaged using intron-targeting FISH probes. Scale bars, 20 µm. **b**, Proximity of pre-*TTN* with mature transcripts, imaged using exon-targeting FISH probes. Arrows denote co-compartmentalization, highlighted in ~2.5–3x zoomed images. Scale bars: 10 µm. Minimal distances between *TTN* and the nearest punctum of the putative interacting RNA were quantified and plotted on the right; the number of cells in each condition is indicated below, pooled from ≥ 3 independent biological replicates. Internal box and whisker plots indicate median, 25th, and 75th percentile, and the 5th-95th percentile range. *p*-values by Kruskal-Wallis test followed by Dunn's multiple comparisons *vs* WT. n.s. > 0.05; ** < 0.01; *** < 0.001; **** < 0.0001.

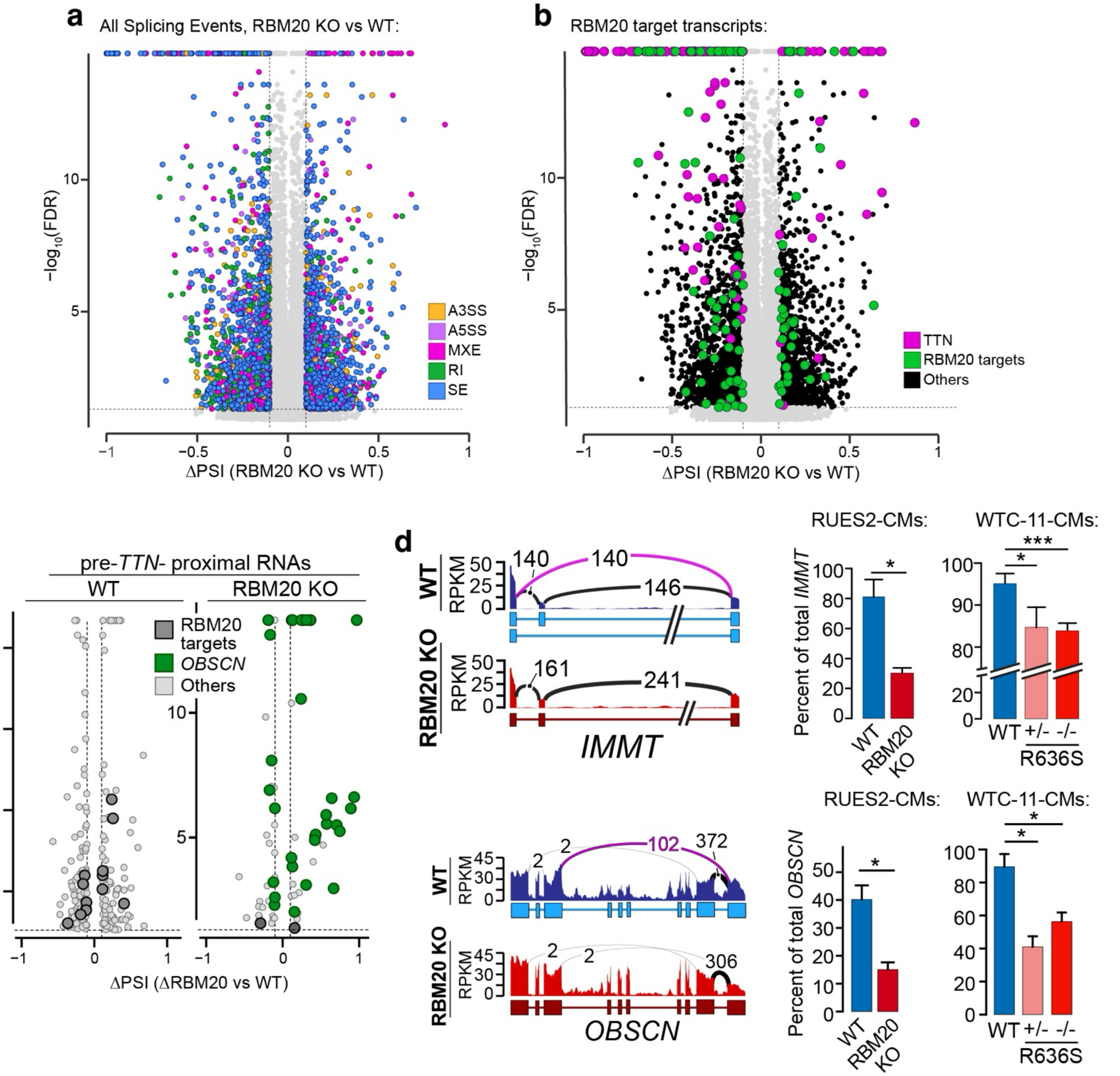


**Supplementary Figure 8. Loss of RBM20 induces pervasive alternative splicing events beyond its direct target transcripts. a,** Volcano plot of all alternative splicing events observed between RBM20 KO and WT CMs, colored according to event-type (**Fig. 5a**). **b,** Same data as in (**a**), highlighting known RBM20 target transcripts[7] (*green*), including *TTN* (*magenta*). Note the large population of alternative splicing events in the broader, non-RBM20-targeted transcriptome (*black*). Data from RUES2 CMs. **c,** Absolute splicing changes among genotype-selective *TTN* proximal transcripts are more pronounced in RBM20 KO cells. WT factory-resident transcripts (*left*) exhibited splicing effect sizes lower than the broader, non-resident transcriptome (0.731% among resident transcripts vs. 0.945% among non-residents), while RBM20 KO resident transcripts harbored some of the most pronounced splicing changes observed (0.974% among residents vs. 0.915% in non-residents). Much of this signal was driven by *OBSCN*, a giant, heavily spliced RBM20 target that preferentially compartmentalizes with *TTN* upon RBM20 loss. In contrast, the alternatively spliced WT-resident transcriptome was generally depleted of RBM20 targets, suggesting that factory-resident splicing regulation in WT cells is not dominated by direct RBM20 interactions. **d,** Examples of altered splicing in a WT-specific SE event (*IMMT, top*) and an RBM20 KO-specific MXE event (*OBSCN, bottom*) in pre-*TTN*-proximal transcripts. *Left:* Sashimi plots visualizing splicing patterns in each cell type, depicted as in (**Fig. 5e**). Splicing events highlighted in magenta were targeted for qRT–PCR quantification. *Right:* qRT–PCR analysis of SE event between *IMMT* exons 5–7, and an MXE event between OBSCN exons 48–55, in female ESC-derived RBM20 KO CMs and male iPSC-derived CMs bearing DCM-associated point mutations (n=4 biological replicates, run in technical quadruplicate; p-value: paired two-sided Student's t test).

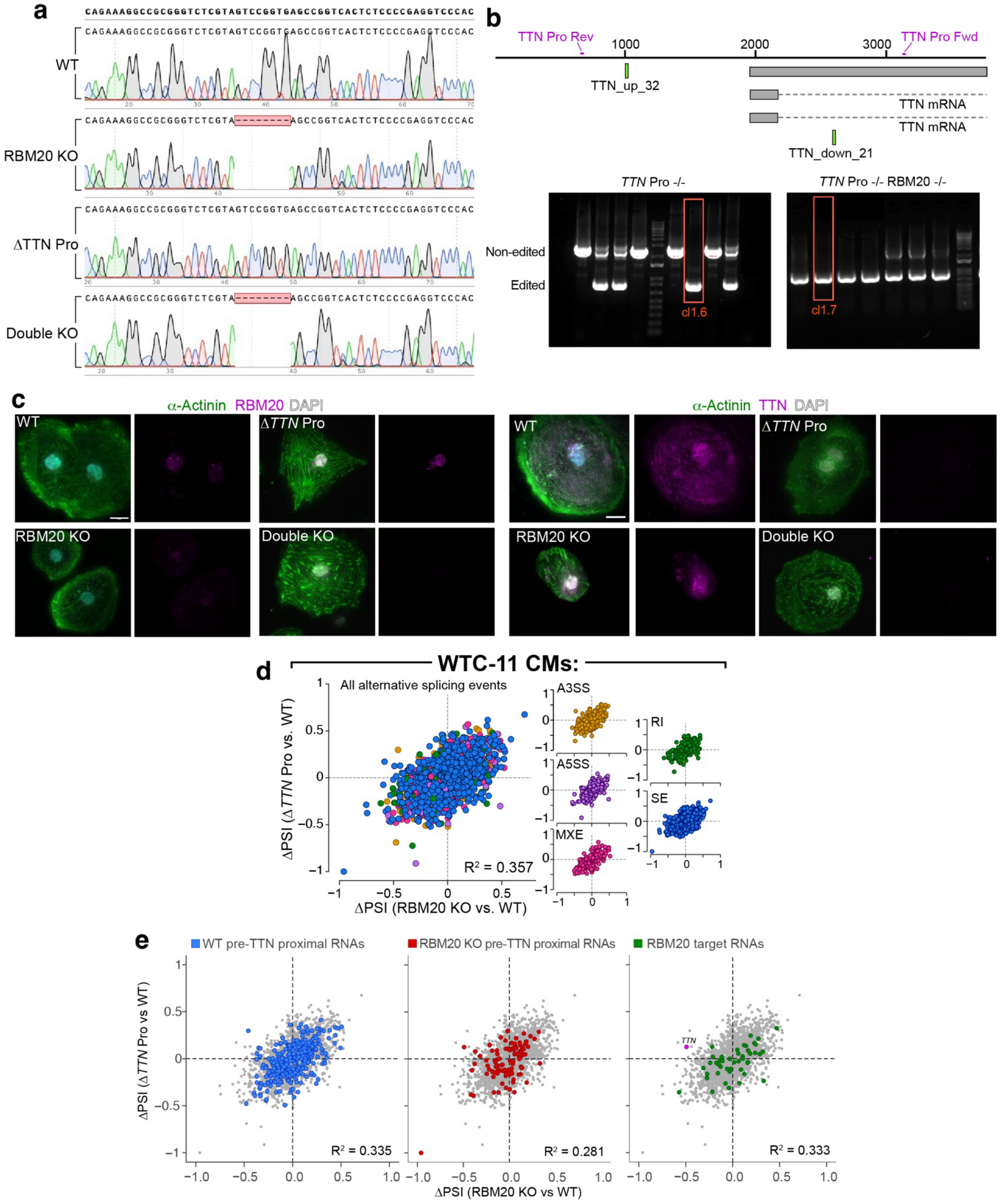


**Supplementary Figure 9. Validation of the RBM20 KO–Δ*TTN* Pro double knockout line. a–b,** RBM20 KO validation. **a,** Targeted genomic sequencing of the RBM20 gene region surrounding the CRISPR editing site[8]. Data are shown for WT WTC-11 iPSCs, and RBM20 KO, Δ*TTN* Pro, and Double KO cells derived from this line. Note the homozygous 8 bp deletions exclusive to RBM20 KO and Double KO cells. **b,** Δ*TTN* Pro validation. *Top*:

genotyping strategy showing the positions of the forward and reverse primers (*blue*) surrounding the *TTN* promoter, and of the two CRISPR/Cas9 sgRNAs used to delete it (*green*). *Bottom:* diagnostic agarose gel, demonstrating targeted homozygous deletion in screened clonal cell lines. Selected clones for the Δ*TTN* Pro and Double KO lines are indicated. **c**, Immunofluorescence (IF) validation of CMs differentiated from each cell line. α-actinin (*green*) is a marker of cardiomyocytes, indicating that the targeted edits did not impair differentiation. RBM20 and TTN are labeled in magenta; nuclei are counterstained with DAPI. Scale bars = 20 μm. **d–e,** Global splicing changes in Δ*TTN* Pro WTC-11 CMs parallel those in RBM20 KO WTC-11 CMs. Splicing events are only shown if they significantly differ (FDR < 0.05 and |ΔPSI| > 0.10) with at least one of the 3 perturbations. **d,** Loss of pre-*TTN* in WTC-11 CMs induces splicing changes that parallel those observed in RBM20 KO cells. PSI, Percent Spliced In. Only events significantly altered in at least one perturbation relative to WT are shown. **e,** Loss of the *TTN* promoter partially paralleled the effects of RBM20 loss across many transcript classes, including pre-*TTN* proximal transcripts in WT (*left*) and RBM20 KO (*middle*) cells, and known RBM20 targets.

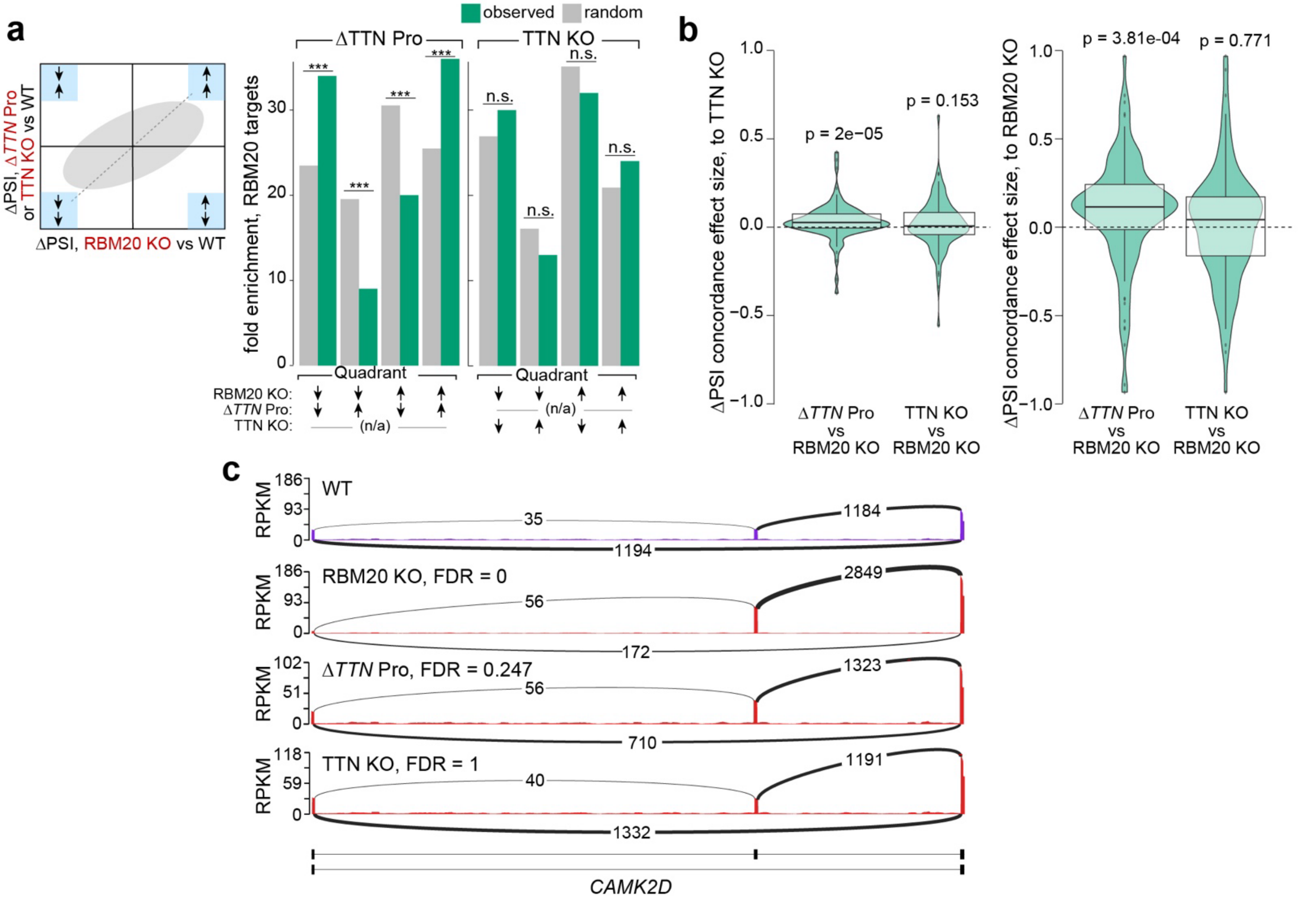


**Supplementary Figure 10. Loss of the *TTN* RNA parallels RBM20-dependent splicing changes more closely than loss of the titin protein. a,** *left:* schematic summarizing the quadrant computational approach. Splicing changes in Δ*TTN* Pro and TTN KO cells were grouped based on their directional concordance or discordance with splicing changes in RBM20 KO cells. *Middle:* RBM20 KO-induced splicing changes correlate with those induced in Δ*TTN* Pro cells, but not TTN KO cells (*right*). Events in each quadrant are plotted separately, depending on whether they are upregulated (+) or downregulated (-) in RBM20 KO, Δ*TTN* Pro, and TTN KO cells. *p*-values were calculated empirically relative to a random model (*gray*) and adjusted with Benjamini Hochberg correction for multiple comparisons. *** $p_{adj} < 0.001$; ** $p_{adj} < 0.01$, * $p_{adj} < 0.05$. **b,** *Left-hand graph:* ΔPSI distributions of the Δ*TTN* Pro vs. WT, and TTN KO, vs WT, multiplied by the sign of the corresponding ΔPSI changes in the RBM20 KO vs WT comparison. If both perturbations (*e.g.*, Δ*TTN* Pro and RBM20 KO) alter PSI in the same direction then the result will be positive; opposing results will be negative. A null model in which splicing differences vary randomly between the two samples will have a mean of 0. Note the positive and significant ($p = 2 \times 10^{-5}$) correlation between Δ*TTN* Pro and RBM20 KO (*left*) but not between TTN KO and RBM20 KO (mean = 0, $p = 0.153$; *right*). *Right-hand graph:* similar calculation, beginning with the ΔPSI values from RBM20 KO vs WT, and multiplying by the signs of the corresponding (Δ*TTN* Pro vs WT, *left*) and (TTN KO vs WT, *right*) comparison. Again note the positive and significant ($p = 3.81 \times 10^{-4}$) correlation between Δ*TTN* Pro and RBM20 KO (*left*), but not between TTN KO and RBM20 KO ($p = 0.771$, *right*). *p*-values were calculated with a one-sample Wilcoxon test ($\mu = 0$). **c,** Representative sashimi plots of a skipped exon event in the RBM20 target transcript *CAMK2D*, across cell lines. FDRs were calculated in relation to the WT using the rMATS pipeline[24]. Note that Δ*TTN* Pro parallels RBM20 KO while TTN KO does not, indicating that this effect is associated with loss of the *TTN* pre-mRNA and not just the Titin protein.

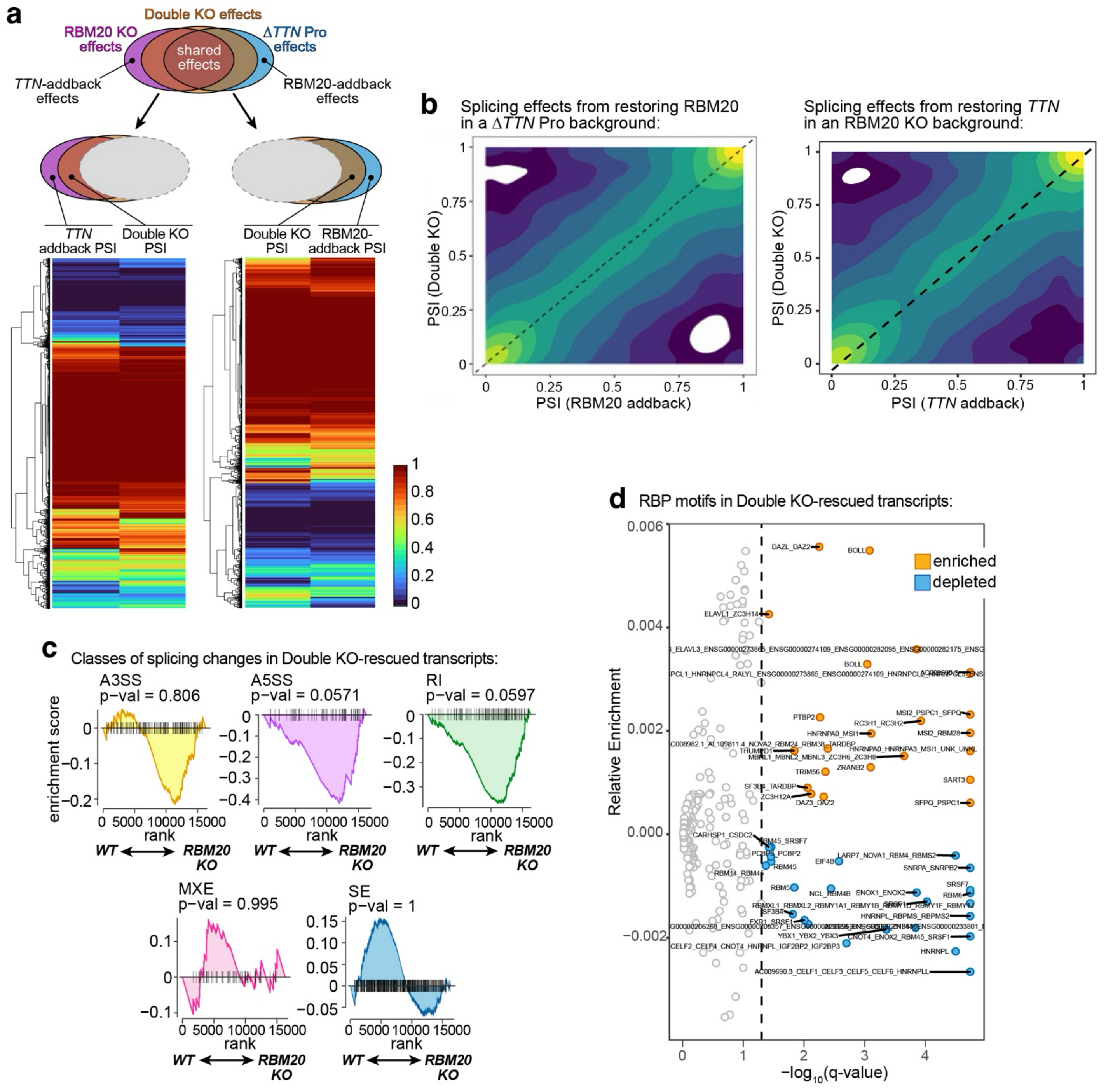


**Supplementary Figure 11. RBM20 KO-induced splicing events attenuated after loss of the *TTN* RNA. a,** Epistasis framework for dissecting RBM20- and TTN-dependent splicing effects. Splicing changes (PSIs) were quantified in Δ*TTN* Pro, RBM20 KO, and Double KO cardiomyocytes. *Top Left:* Events altered in the Δ*TTN* Pro single-mutant (FDR < 0.05) were removed to isolate splicing changes specific to RBM20 loss, in the presence or absence of the *TTN* RNA. Comparing these two residual sets revealed splicing events induced by "adding back" *TTN* in the absence of RBM20. *Top right:* The analogous analytical pipeline was used to identify splicing effects induced by "adding back" RBM20 in the absence of *TTN* expression. Heatmaps (*bottom*) show PSI values for these event sets, hierarchically clustered and combining all splicing event types. These data correspond to the axes in panel (b). **b,** Deletion of the *TTN* promoter is epistatic to RBM20 loss. *Left:* 2D density plot of PSI values comparing RBM20 addback and Double KO lines, after removing Δ*TTN* Pro single-mutant events. Restoring RBM20 in a Δ*TTN* Pro background has minimal effect on global splicing levels, as indicated by strong diagonal symmetry. *Right:* The same analysis for *TTN* "addback." Restoring *TTN* in the absence of RBM20 induces comparably broader shifts in PSI, reflected by off-diagonal skewing. **c–d,** Analysis of splicing events dysregulated in RBM20 KO cells and attenuated in the Double KO (RBM20 KO/Δ*TTN* Pro) cells. **c,** Attenuated events are not enriched for specific splicing classes. Gene Set Enrichment Analysis (GSEA) was performed for each category of splicing event, ranking genes based on $log_2$[(WT pulldown/Input) / (RBM20 KO pulldown / Input)]. All genes that are differentially spliced (FDR < 0.05 and $|\Delta PSI| > 0.10$) in RBM20 KO cells but rescued in Double KO cells were analyzed. Although most distributions show a modest rightward shift, suggesting preferential enrichment in RBM20 KO cells, none reached statistical significance. **d,** Candidate RNA-binding protein (RBP) motifs enriched in rescued transcripts. Volcano plot of Transite[25] motif-enrichment among transcripts that are dysregulated in RBM20 KO cells and attenuated in the Double KO line. The relative enrichment is computed by normalizing the absolute enrichment over the k-mer abundance in the reference set; *p*-values were adjusted for multiple comparisons with the Benjamini-Hochberg correction. Similar motifs were aggregated based on TOM-TOM[26] similarity.

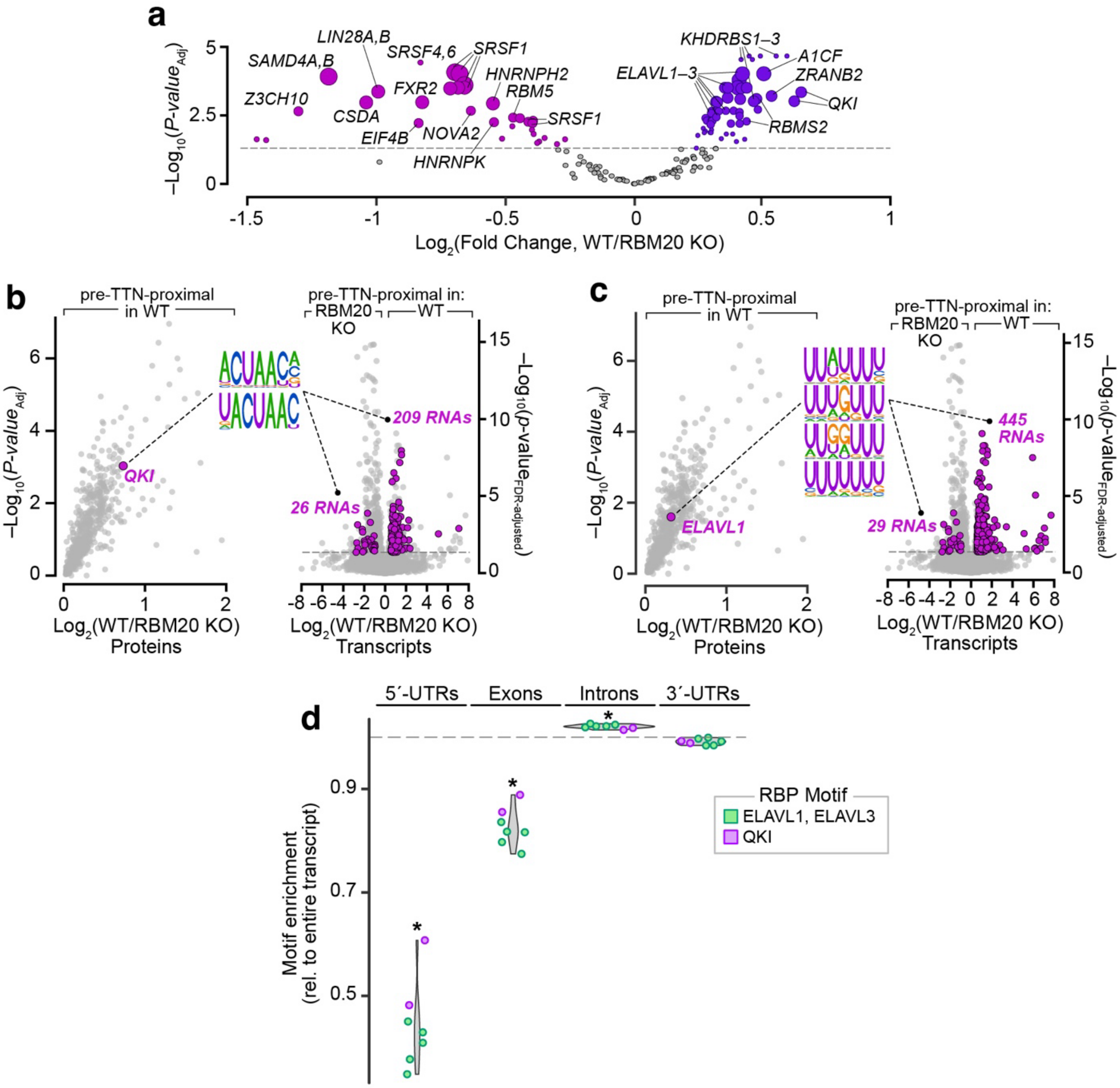


**Supplementary Figure 12. Concordance of RNA-binding proteins and their cognate motifs in the TTN RNA factory proteome and transcriptome. a,** Differential enrichment of RNA-binding protein recognition motifs in WT- and RBM20 KO-specific pre-TTN-proximal transcripts. The size of each motif's circle is proportional to the total number of transcripts harboring that motif. Significant differential motifs are highlighted in purple ($\log_2$(fold change) > 0.58; FDR < 0.05). **b–c**, The factory-proximal proteome and transcriptome show concordant enrichment of selected RBPs and their recognition motifs. *Left*: significant enrichment of each protein (*highlighted in magenta*) in O-MAP-MS. Data are the same as in (**Fig. 6b**). *Middle*: Logos for the major RNA recognition motifs for each RBP. *Right*: Incidence of these motifs within the WT- and RBM20 KO pre-*TTN*-proximal transcriptomes, observed by O-MAP-Seq (*highlighted in magenta*). Data are the same as in (**Fig. 4e**). The number of transcripts bearing each binding motif is indicated. **b,** QKI; **c,** ELAVL1. **d**, ELAVL1 and QKI motifs are predominantly localized in introns. *Transite* analysis discretely applied to mRNA functional domains (5´–UTR, exons, introns, 3´–UTR), comparing WT to RBM20 KO-enriched transcripts. Asterisks denote significantly enriched or de-enriched RNA domains ($p < 0.05$).

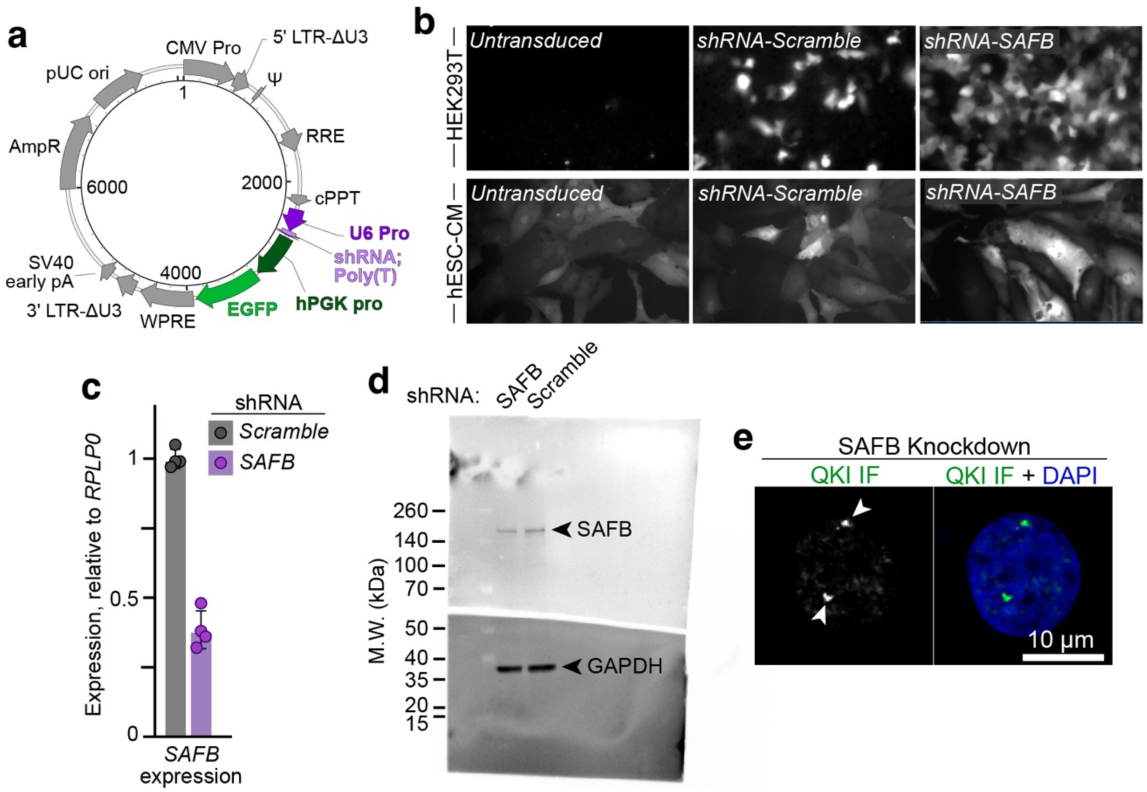


**Supplementary Figure 13. Validation of SAFB knockdown. a,** Lentiviral shRNA vector design (VectorBuilder Inc., Chicago, IL, USA). Transduction efficiency was assessed by eGFP expression rather than antibiotic selection to avoid toxicity in cardiomyocytes. **b,** Representative images illustrating high transduction efficiency in HEK293T cells (*top*) and hESC-derived cardiomyocytes (*bottom*). **c,** Validation of shRNA targeting. HEK293T cells were transduced with lentivirus encoding shRNA-SAFB (*purple*), or a scrambled negative control (*grey*), and *SAFB* expression was quantified by RT–qPCR. Bars represent mean ± s.d. (n = 4 technical replicates). **d,** Western blot validation demonstrates specific, but modest depletion of SAFB following shRNA-SAFB treatment (GAPDH loading control). Uncropped images of the full membrane are shown; the membrane was split to enable simultaneous probing of both targets. **e,** Representative images do not reveal an obvious change in QKI localization after partial SAFB depletion. Scale bars, 10 µm.
